# Kar4 acts as a Ste12 regulator in *Saccharomyces cerevisiae*, promoting Ste12 binding to a specific DNA motif genome-wide

**DOI:** 10.64898/2025.12.09.693310

**Authors:** Jason V. Rogers, Amanda Yeo, Val Meleshkevich, Hernan Lorenzi, Mark D. Rose, Orna Cohen-Fix

## Abstract

Kar4 is a putative transcription factor required for efficient mating in the budding yeast *Saccharomyces cerevisiae*. Kar4 functions with Ste12, the master transcriptional regulator of the yeast mating pheromone response, to promote the transcription of a subset of Ste12 targets required for mating. However, the mechanism by which Kar4 modulates Ste12 activity has remained uncertain. Here, we examined Kar4’s function at the levels of transcription (RNA-seq) and Ste12 DNA binding (ChIP-exo). We show that Kar4 promotes Ste12 binding to nearly all Ste12 DNA-binding sites associated with transcriptionally-upregulated genes, even if their upregulation is not dependent on Kar4. We further found that the majority of Ste12-binding sites have two pheromone response elements (PREs) separated by four nucleotides in a head-to-tail orientation (H-T 4 motif). Sites associated with Kar4-dependent transcription have PREs with more mismatches and substantially lower Ste12 occupancy in *kar4*Δ cells than sites linked to Kar4-independent transcription. During the pheromone response in *kar4*Δ cells, Ste12 exhibits increased binding to non-H-T 4 motifs, particularly T-T 3 motifs, resulting in the abnormal upregulation of many genes not transcribed during the wild-type pheromone response. Therefore, we propose that Kar4 functions by increasing the DNA-binding specificity of Ste12 globally, promoting its binding primarily to H-T 4 motifs. Our model is consistent with previously-observed slower induction kinetics of Kar4-dependent genes via a feed-forward mechanism. Lastly, we uncovered several novel aspects of the pheromone response, including a broad role for the Crz1 transcription factor, induction of stress responses, and the identification of pheromone-responsive intergenic transcripts.

## Introduction

Regulation of gene expression is a critical part of many cellular and developmental processes. One aspect of this process is the mechanism by which transcription factors recognize their intended targets. The present study uses the transcriptional response to pheromone in the budding yeast *Saccharomyces cerevisiae* to explore this question. Budding yeast can exist as haploid cells of one of two mating types: *MAT***a** or *MAT*α (Bardwell 2005). Cells of opposite mating types can mate through a process that is initiated by the secretion of a mating-type-specific pheromone (a-factor in *MAT***a** cells and α-factor in *MAT*α cells) and expression of a receptor for the pheromone of the opposite mating type (e.g., α-factor receptor in *MAT***a** cells). When cells of opposite mating types are in close proximity, pheromone binding to the cognate receptor activates a MAPK-signaling cascade, ultimately resulting in G1-arrest, cell polarization toward the direction of pheromone, and activation of the Ste12 transcription factor. Several hundred genes downstream of Ste12 activation mediate the remainder of the mating process, including mating projection (“shmoo” tip) formation, cell fusion, nuclear congression, and nuclear fusion, ultimately resulting in a single diploid cell (Roberts *et al*. 2000; Sieber *et al*. 2023).

Ste12 is expressed constitutively (Esch *et al*. 2006) and preemptively binds the promoters of a subset of genes that are upregulated in response to pheromone (Zeitlinger *et al*. 2003; Rossi *et al*. 2021). However, Ste12 does not induce transcription before pheromone is detected, as the Ste12 transcriptional activation domain is inhibited by Dig1 and Dig2 (Pi *et al*. 1997; Tedford *et al*. 1997). The prevailing model is that, in the presence of pheromone, MAPK-signaling results in Fus3-mediated phosphorylation of Dig1 and Dig2, thereby relieving repression of Ste12 (Cook *et al*. 1996), leading to immediate transcriptional activation for a subset of pheromone-induced genes to which Ste12 was pre-bound. For other genes, Ste12 appears to bind only after pheromone treatment (Zeitlinger *et al*. 2003), leading to slower induction kinetics. In addition, the transcription of Ste12 itself is modestly upregulated by pheromone induction (Roberts *et al*. 2000), although Ste12 protein levels decline during the pheromone response (Esch *et al*. 2006).

Ste12 binds DNA at pheromone response elements (PREs), optimally 5’-TGAAACA-3’ (Kronstad *et al*. 1987; Van arsdell and thorner 1987; Hagen *et al*. 1991; Yuan and fields 1991). In vitro, the purified DNA-binding domain (residues 1-215) has been shown to bind with low affinity to a single PRE, or cooperatively with higher affinity to a pair of PREs, suggesting that in vivo Ste12 binds as a dimer (Yuan and fields 1991). Further, in vitro Ste12 can bind multiple PRE orientations and spacings (Yuan and fields 1991). Intriguingly, however, in vitro HT-SELEX (High-Throughput Systematic Evolution of Ligands followed by EXponential enrichment) experiments on Ste12 demonstrated that it has the highest affinity toward PRE pairs in a tail-to-tail orientation separated by 3 nucleotides (T-T 3 motif, specifically GTTTCA-NNN-TGAAAC) (Dorrity *et al*. 2018).

In vivo, pheromone-induced gene promoters contain multiple PREs and PRE-like sequences (imperfect PREs) in a wide variety of orientations and spacings (Chou *et al*. 2006; Su *et al*. 2010; Aymoz *et al*. 2018). However, few rules regarding Ste12 binding in vivo have been deciphered. It has been demonstrated that the strength of transcriptional induction is correlated with relative PRE affinity using β-Gal assays (Su *et al*. 2010; Pinheiro *et al*. 2025). Additionally, there appear to be many possible orientations and spacings which are functional (Hagen *et al*. 1991; Su *et al*. 2010; Pinheiro *et al*. 2025), often with some redundancy between PREs when more than 2 are present (Su *et al*. 2010; Aymoz *et al*. 2018). A recent study using synthetic promoters found that, while all PRE pair orientations can be functional, only a subset of possible spacings are utilized in vivo, generally either 3 or 4 bp spacing (depending on the orientation) followed by 10 bp increments (one helical turn; e.g., 4, 14, 24, etc.) (Pinheiro *et al*. 2025). The genomic context of the PREs also matters, as PREs positioned in nucleosome-depleted regions give rise to faster induction and a higher transcriptional output (Aymoz *et al*. 2018; Pinheiro *et al*. 2025).

In addition to Ste12, Kar4 is essential for the full transcriptional response to pheromone. Kar4 is essential for mating, with *kar4Δ* mutants exhibiting nuclear congression defects, resulting in unfused nuclei (Kurihara *et al*. 1994). This specific defect can be rescued by simultaneously overexpressing *KAR3* and *CIK1*, two proteins involved in the microtubule-dependent movement of nuclei (Kurihara *et al*. 1996). By microarray analysis, Kar4 was shown to be required for the full pheromone-mediated transcriptional induction of 70 genes (of which 29 were induced in wild-type ≥ 2.5-fold), including but not limited to *KAR3* and *CIK1* (Lahav *et al*. 2007). A class of genes was also identified which appeared repressed by Kar4, as they were induced by pheromone only in *kar4*Δ mutants (Lahav *et al*. 2007). These findings led to a model in which Kar4 and Ste12 cooperatively bind at the promoters of Kar4-dependent genes. As Kar4 itself is induced by pheromone (although it also has moderate expression prior to pheromone), it was proposed that Kar4 functions in a feed-forward loop to mediate expression of many genes expressed later during the pheromone response, whereas genes induced earlier and directly by Ste12 are Kar4-independent (Lahav *et al*. 2007). Kar4 also has non-mating roles, functioning in the N6-methyladenosine methyltransferase complex (MTC) during early meiosis to regulate gene expression via RNA methylation (Ensinck *et al*. 2023; Park *et al*. 2023b; Park *et al*. 2023c). However, as numerous alleles were isolated for Kar4 which disrupt only its mating or meiotic functions, its roles in these two processes are separate (Park *et al*. 2023c).

Few studies since Lahav et al. (2007) have investigated the role of Kar4 in the transcriptional response to pheromone. Aymoz et al. (2018) studied the promoters of *AGA1* and *FIG1* as model early (and Kar4-independent) and late (and Kar4-dependent) promoters, respectively. Chromatin IP (ChIP) revealed that Ste12 binds more slowly to the p*FIG1* promoter during the pheromone response, as expected. However, Kar4 exhibited promoter binding kinetics that were identical to Ste12 for both p*AGA1* and p*FIG1*. Further, in both cases Kar4 promoter binding was detectable before pheromone treatment, and the binding was Ste12-dependent. That Kar4 broadly binds to the same sites as Ste12 was later verified genome-wide, although it was only tested in asynchronous cells not responding to pheromone (Rossi *et al*. 2021). Ste12 and Kar4 were also shown to physically interact by co-immunoprecipitation, both before and after pheromone treatment (Aymoz *et al*. 2018), consistent with one-hybrid data (Lahav *et al*. 2007). Importantly, Aymoz et al. (2018) demonstrated that by mutating a suboptimal PRE (TGAcACA) to an optimal PRE (TGAAACA), p*FIG1* could be made transcriptionally Kar4-independent. Therefore, they concluded that rather than Kar4 uniquely binding at promoters of Kar4-dependent genes, Kar4 functions as a general Ste12 stabilization factor, effectively increasing Ste12 affinity for all Ste12-binding sites. In support of this model, later work using synthetic promoters with two PREs found no difference in transcription in *kar4*Δ cells when the PREs were optimal, but slower induction (albeit higher final transcriptional output) when the PREs contained mutations (Pinheiro *et al*. 2025).

Outside of mating, Ste12 regulates filamentous growth together with Tec1, where they form a heterodimer at promoters containing an adjacent PRE and a Tec1-binding site (Wong sak hoi and dumas 2010; Cullen and sprague 2012). Similarly, Ste12 can also bind cooperatively with the Mcm1 transcription factor, regulating genes relevant to mating before the pheromone response, including *MAT***a**-specific genes and *FAR1* (Oehlen *et al*. 1996; Wong sak hoi and dumas 2010).

Despite this prior work, our understanding of the role of Kar4 during mating remains incomplete. In this study, we sought to gain a deeper understanding on how Kar4 affects Ste12 function. To this end, we compared the pheromone response between wild-type and *kar4*Δ cells by both RNA-seq and ChIP-exo. We found that Kar4 promotes Ste12 binding to virtually all of the Ste12 DNA-binding sites associated with transcriptionally-upregulated genes during the pheromone response, even if their upregulation does not require Kar4. Furthermore, we found that Ste12 binding has a preference for PREs separated by four nucleotides in a head-to-tail orientation (H-T 4 motif) present in a large majority of Ste12-bound pheromone-induced gene promoters. H-T 4 motifs associated with Kar4-independent transcription typically contained higher affinity PREs with fewer mismatches, whereas those linked to Kar4-dependent transcription contained more mutations, often in the first 2 bp of the PRE. In the absence of Kar4, Ste12 exhibited increased binding to a range of non-H-T 4 motifs, particularly tail-to-tail PREs separated by 3 nucleotides (T-T 3 motifs). In many cases the absence of Kar4 resulted in pheromone-induced gene transcription where none is observed in wild-type cells. Together, we propose that Kar4 functions as a general Ste12 regulator, regulating its DNA-binding specificity genome-wide.

## Results

### Analysis of the transcriptional response to mating pheromone using pre-synchronized cells

The overarching goal of this study was to understand how Kar4 affects Ste12’s function. To this end, we took two orthogonal approaches: the first was to re-examine the effect of Kar4 deletion on the transcriptional response to pheromone. The second was to determine the locations of Ste12’s DNA binding in the presence and absence of Kar4. Although gene expression during the pheromone response in the absence of Kar4 activity has been measured previously by microarray (Lahav *et al*. 2007), we sought to revisit this topic using several improved methodologies. First, we used RNA-sequencing (RNA-seq), which enables the quantification of all RNA transcripts, including non-coding RNAs. Second, we pheromone-treated cells pre-synchronized in G1 (**Figure 1a** and see below), rather than asynchronous cultures, as was done previously. This removed any confounding gene expression effects due to changes in cell cycle rather than a genuine pheromone response. Moreover, in asynchronous cultures exposed to pheromone, cells will take varying amounts of time to fully respond (Colman-lerner *et al*. 2005), potentially decreasing the signal-to-noise ratio. Our experimental design enabled us to discover several new aspects of the yeast mating pheromone response unrelated to Kar4, including Crz1-dependent gene expression, accurate measurements of gene downregulation in response to pheromone, the activation of a stress response, the identification of a Ste12-independent pheromone response, and the identification of pheromone-responsive intergenic transcripts, many of which are likely to be non-coding. These data are presented and discussed in the accompanying Appendix. Here, we focus primarily on the role of Kar4 during the pheromone response and the identification of a prevalent Ste12 binding motif.

**Figure 1.**
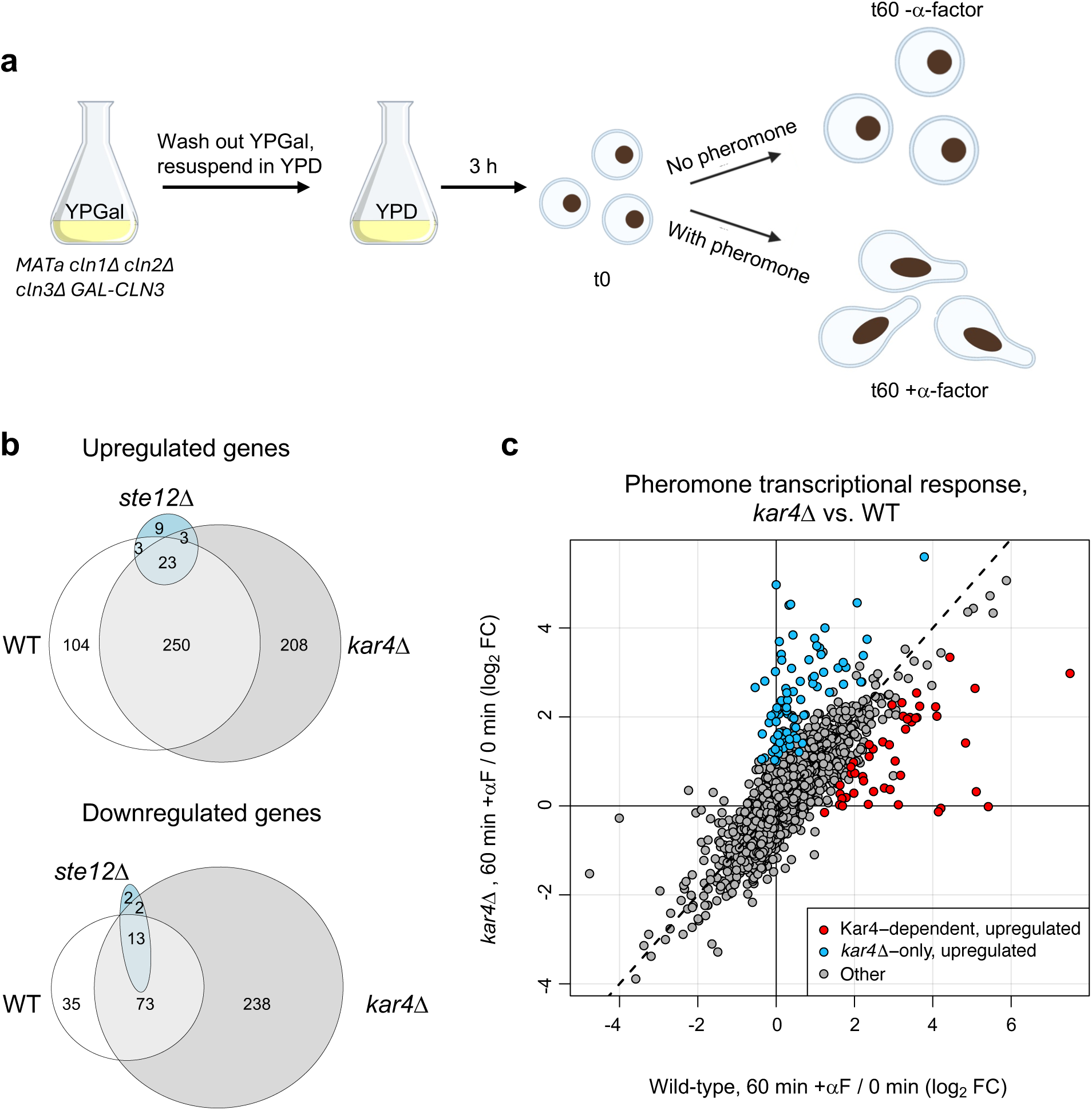
Analysis of the transcriptional response to mating pheromone using pre-synchronized cells. a) Experimental design used for RNA-seq. A strain with all G1 cyclins deleted and with *CLN3* under a galactose-inducible promoter was transferred from growth media with galactose (YPGal, “*CLN3* on”) to media with glucose (YPD, “*CLN3* off”) to induce a G1 arrest. Arrested cells were then used for RNA-seq either without (time 0 or time 60 - α-factor) or with pheromone for 1 hour (time 60 + α-factor). Image created using BioRender. b) Venn diagram of significantly up and down-regulated genes (and at least a 2-fold change compared to time 0) after pheromone treatment for the indicated genotypes. c) Scatterplot comparing gene expression log_2_ fold-changes (FC) after pheromone treatment for *kar4*Δ vs. wild-type cells. Dashed line indicates 1:1 line. Kar4-dependent upregulated genes and *kar4*Δ-only upregulated genes (see main text) are indicated with red and blue circles, respectively.

To re-examine the effect of Kar4 on pheromone-induced gene expression, we performed RNA-seq on *MAT***a** wild-type (WT), *ste12Δ*, and *kar4Δ* cells. To synchronize cells in G1, strains were deleted for the three G1 cyclin genes (*CLN1, CLN2 and CLN3*), and *CLN3* was expressed using a galactose-inducible promoter (Stone *et al*. 2000). Cells were arrested in G1 by switching from a galactose-containing medium to a glucose-containing medium, where *CLN3* expression is repressed. Arrested cells (t0, **Figure 1a**) were either treated for 60 minutes with alpha factor (αF) mating pheromone (t60 + α-factor) or left untreated for 60 minutes (t60 - α-factor). (**Figure 1a**). Samples were collected at t0 and t60. The t0 and t60 + α-factor conditions in all three genotypes were performed in biological quadruplicates: two biological replicates in two separate experiments each. Separately, samples for wild-type cells were taken at t60 with or without α-factor, in duplicates. Note that at this point, unless otherwise noted, we restricted analyses to nuclear protein-coding genes and excluded Ty elements (long terminal repeat retrotransposons) (see Methods).

To establish a control dataset, we compared changes in gene expression in response to pheromone in wild-type cells relative to either t0 or to t60 - α-factor (**Figure 1a)**. The correlation between these two comparisons was excellent (R = 0.85), indicating that the extra 60 minutes in G1 had little effect on overall gene expression (**Supplementary Figure S1a, Supplementary File S1 and Supplementary File S2**). Because the quality of the t0 data was better (more replicates), we used these data sets as our control. In the differential expression analyses below we removed genes whose expression was likely affected by the longer arrest in G1 (4 genes from the upregulated gene list and 25 genes from the downregulated one; **Supplementary Figure S1b, Supplementary File S3** and see Materials and Methods).

Next, we conducted quality control analyses to determine reproducibility and identify general patterns in the data (**Supplementary Figure S2a and b**). For the t0 and t60 + α-factor samples of all genotypes, our principal component analysis (PCA) indicated that the replicates were similar to each other. There was a small batch effect between the two experiments (PC2), although it did not impact the major effect of the response to mating pheromone (PC1**, Supplementary Figure S2a**; batch effect corrected in subsequent differential expression analyses, see Methods). As expected, *ste12Δ* cells did not respond to α-factor, but interestingly they grouped separately from other genotypes at time 0, likely due to genes that are controlled by Ste12 prior to pheromone treatment. Also as expected, the similarity plot showed that the samples were most strongly grouped by genotype and response to α-factor (**Supplementary Figure S2b**). While *ste12*Δ samples were most closely related to non-α-factor treated samples, they again were distinctly different (**Supplementary Figure S2b**).

### RNA-seq analysis of synchronized wild-type, *kar4*Δ, and *ste12*Δ cells reveals novel patterns of gene expression

To identify broad patterns in the RNA-seq data, we performed hierarchical clustering of gene expression changes for all genes compared to the average expression of samples from wild-type at t0 (**Figure 2; Supplementary File S1**). The largest clusters contained genes that were upregulated (cluster I) or downregulated (cluster II) in response to pheromone in both wild-type and *kar4*Δ cells, but not in *ste12*Δ cells. The appearance of down-regulated genes is interesting, as those have been previously attributed to cell cycle differences (Roberts *et al*. 2000). In our experimental setup, however, cell cycle differences can be discounted. This group of genes revealed the activation of a cellular stress response in response to pheromone (see Appendix). A subset of the upregulated genes also required Ste12 for their basal expression in G1-arrested cells prior to pheromone treatment (cluster I-A). As has been previously observed (Lahav *et al*. 2007), a small subset of upregulated genes required Kar4 for full induction during pheromone treatment (cluster I-B). We also observed a cluster of genes that were not normally regulated by pheromone, but became strongly upregulated in the *kar4*Δ mutant (cluster III; referred to as “*kar4Δ*-only” genes below). As with cluster I, a subset of these genes required Ste12 for their basal expression (cluster III-A), while the remainder did not (cluster III-B). Lastly, we observed a small cluster of genes that were upregulated by pheromone in a Ste12-independent manner (cluster IV, see Appendix).

**Figure 2.**
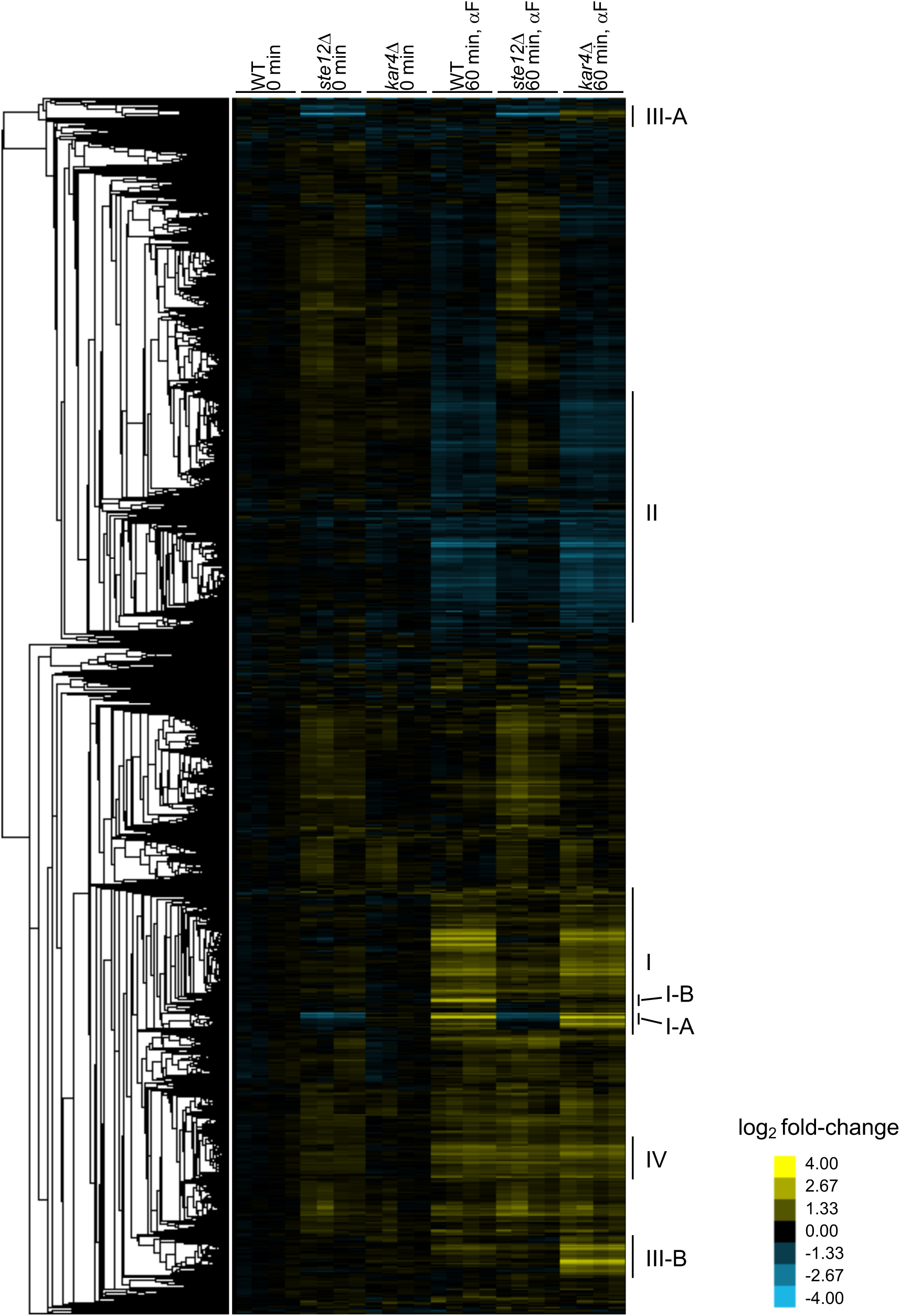
Global hierarchical-clustering overview of RNA-seq data. Log_2_ fold-changes for wild-type (WT), *kar4Δ*, and *ste12Δ*, at time 0 min or time 60 min with alpha factor (αF), were calculated against the average transcripts per million (TPM) values for WT 0 minutes (see Methods). Each column group contains 4 samples (biological quadruplicates). Broad cluster groupings are indicated and discussed in main text. Briefly, clusters are I (pheromone-induced genes), I-A (pheromone-induced and require Ste12 for basal expression), I-B (pheromone-induced and Kar4-dependent), II (pheromone-repressed genes), III (*kar4*Δ-only), III-A (*kar4*Δ-only and require Ste12 for basal expression), III-B (*kar4*Δ-only and do not require Ste12 for basal expression), and IV (pheromone-induced and Ste12-independent).

Differential gene expression analysis revealed that 380 genes were significantly upregulated after pheromone addition to wild-type cells (at ≥ 2-fold, here and for all remaining analyses, unless noted otherwise), while 121 genes were significantly downregulated (**Figure 1b; Supplementary File S2**). In the absence of Kar4, the number of genes affected by pheromone treatment was greater than in wild-type, with 484 genes upregulated and 326 genes downregulated (**Figure 1b**). As expected, there was considerable overlap between the two strains, with most wild-type-regulated genes forming a subset of genes regulated in *kar4*11 cells (**Figure 1b**). The large increase in the number of genes differentially affected by the absence of Kar4 appeared primarily due to a slightly increased global pheromone response, which was more evident for the downregulated genes (**Figure 1c**, note the shift off the diagonal line). Due to these differences in expression, several Gene Ontology (GO) terms were unique to the pheromone response in *kar4Δ* cells, including ribosome biogenesis (see below). As expected, the majority of gene expression changes were Ste12-dependent (**Figure 2**).

To compare our data to previously obtained datasets of pheromone-induced gene expression, we compared our list of wild-type differentially expressed genes to microarray data from asynchronous wild-type cells exposed to pheromone for 120 minutes, using the same 2-fold change cutoff (Roberts *et al*. 2000). 139 genes were upregulated in both datasets, which is a 37% overlap of the Roberts et al. data to ours and 37% overlap of our dataset compared to theirs, with an overall correlation of 0.53 (**Supplementary Figure S3a and b**). We suspect that the moderate overlap between the two experiments is largely due to the difference in using RNA from G1-arrested cells (our study) versus asynchronous cells (Roberts et al. study) as the control dataset.

Using these datasets, we determined pathways that were overrepresented in the transcriptional response to pheromone in the three genetic backgrounds (wild-type, *ste12Δ*, and *kar4Δ*) through standard Gene Ontology (GO) enrichment analysis using Saccharomyces Genome Database terms (Ashburner *et al*. 2000; Gene ontology *et al*. 2023). Only categories with 3 or more genes were considered. The analysis included upregulated genes and downregulated genes that met the differential expression cutoffs of fold-change ≥ 2 and adjusted p-value of < 0.05 (**Supplementary File S4**). In the presence of pheromone, pathways that were transcriptionally upregulated in wild-type cells were related to the mating process and to metabolism, in agreement with a previous analysis of the Roberts et al. (2000) dataset (Williams *et al*. 2016). In *ste12Δ* cells, genes that were upregulated in response to pheromone were also mostly related to metabolism (see Appendix). Excluding these categories from the wild-type and *kar4Δ* overrepresentation lists, the processes that were activated in a Kar4-independent manner included the expected categories of reproduction, cell wall organization, and karyogamy. As these categories were also identified in the *kar4Δ* strain, the requirement for Kar4 does not appear specific for a particular process, but instead Kar4 appears to regulate genes in all the mating-related categories. However, consistent with an increased global pheromone response as noted above, more processes were upregulated in the *kar4Δ* strain than in wild-type, including processes that are not obviously directly linked to mating, such as sporulation, selective autophagy, and detoxification.

When examining overrepresentation among the downregulated genes in response to pheromone, two observations stood out: first, in wild-type cells, the major pathways that are downregulated are involved in transport of ions and small molecules. Second, in the absence of Kar4, a large number of ribosome and translation-related genes are downregulated. Both patterns are reminiscent of a stress response (see Appendix), and the latter can be observed also in wild-type cells, albeit to a lesser degree (i.e. differential expression less than a log_2_ fold change of 1, **Supplementary File S2**), which is why it did not appear enriched in the GO enrichment analysis. The presence of pheromone-induced downregulated genes was previously attributed to differences in cell cycle states (Roberts *et al*. 2000; Williams *et al*. 2016), and we suspect that our experimental design, whereby the control group was cells arrested in G1, revealed gene categories that were masked when the control group was asynchronous cells.

### Identification of three gene-expression patterns related to Kar4: Kar4-independent, Kar4-dependent, and *kar4*Δ-only

We next defined the set of genes whose expression in response to pheromone was Kar4-dependent. Because our overall goal was to examine how Kar4 affects Ste12 function, we used more stringent criteria in defining this gene set. First, to qualify as Kar4-dependent, genes had to be induced at least 2-fold in wild-type relative to their expression at t0 (no pheromone). Second, these changes had to be Ste12-dependent, such that expression in wild-type pheromone-treated cells was at least 2-fold higher than in pheromone-treated *ste12Δ* cells (see Methods). This resulted in 215 genes (214 not including *KAR4*) that were strongly induced by pheromone in wild-type cells in a Ste12-dependent manner (**Supplementary Figure S4 and Supplementary File S3**).

We then identified a subset of 47 Kar4-dependent genes whose expression in pheromone-treated *kar4*Δ cells was at least 2-fold lower than in pheromone-treated wild-type cells (**Figure 1c and Supplementary Figure S4b**). The remaining 167 pheromone-induced genes were classified as Kar4-independent (**Supplementary Figure S4a**). Of note, 50% of the 214 pheromone-induced genes were significantly Kar4-dependent without also imposing a 2-fold cutoff (i.e., genes with a significant p-value even if the fold-change difference between wild-type and *kar4*Δ cells was small). We also identified 5 Kar4-dependent downregulated genes (not discussed further, but shown in **Supplementary Figure S4b**). By GO-term analysis, the Kar4-dependent genes shared no functional enrichment. Importantly, however, the Kar4-dependent gene list includes *KAR3* and *CIK1*, two genes whose pheromone-induction was previously shown to be Kar4-dependent and to underlie the karyogamy defect in *kar4*Δ cells (Kurihara *et al*. 1996). Intriguingly, the second-most Kar4-dependent gene was *IME4*, a known Kar4-interacting protein thought to function exclusively during meiosis (Ensinck *et al*. 2023; Park *et al*. 2023b; Park *et al*. 2023c) (and see below).

The final group of genes we identified were those whose pheromone-induced expression in *kar4Δ* cells was ≥ 2-fold higher compared to their expression in wild-type cells. This group, defined as “*kar4*Δ-only,” included 84 upregulated genes, which was unexpected as it suggested that Kar4 plays a negative role in regulating gene expression (**Supplementary Figure S4c**). This group also included 13 genes that were downregulated 2-fold more in *kar4Δ* cells than their downregulation in wild-type cells. A group of genes with a similar dependence was noted by Lahav et al. (2007). The *kar4Δ*-only group turned out to be informative of Kar4’s potential function and will be revisited later in this study. Together, these data demonstrate that loss of Kar4 function has at least three effects on the pheromone response: reduced upregulation of a subset of genes normally induced by pheromone in wild-type (Kar4-dependent genes), upregulation of a set of genes not normally induced during the pheromone response (*kar4Δ*-only genes), and a slightly increased global pheromone response for many of the remaining pheromone-regulated genes, particularly downregulated genes.

### The mRNA-methylating function of Kar4 is not involved in its regulation of pheromone-induced gene expression

Kar4 has three known activities: (a) regulating transcription during the mating response (Lahav *et al*. 2007), (b) being part of the *N6*-methyladenosine-methyltransferase complex (MTC) (Ensinck *et al*. 2023; Park *et al*. 2023b; Park *et al*. 2023c), and (c) completion of meiosis and sporulation (Kurihara *et al*. 1996; Park *et al*. 2023b). The MTC also includes Ime4, Mum2, Vir1, Slz1, and Dyn2, and the activity of Kar4 as part of the MTC is thought to be confined to meiosis (Ensinck *et al*. 2023; Park *et al*. 2023a; Park *et al*. 2023b; Park *et al*. 2023c). However, *IME4*, but not *MUM2* and *SLZ1*, is highly induced by pheromone in a Kar4-dependent manner (fold-change of ∼180 and ∼8 in wild-type and *kar4Δ* cells, respectively; **Supplementary Files S1 and S2**), prompting us to investigate whether the MTC could play a role in the mating transcriptional response. RNA samples from G1-synchronized wild-type and *ime4Δ* cells, with or without pheromone treatment, were collected and processed as described above (**Supplementary Files S1 and S2**). A similarity matrix revealed that samples grouped most similarly based on treatment and then genotype (**Supplementary Figure S5a**). The pheromone-induced changes in gene expression were highly similar between the wild-type and *ime4Δ* samples, much more than between wild-type and *kar4Δ* cells (compare **Supplementary Figure S5b** and **Figure 1c**), with over 80% of upregulated genes shared between wild-type and *ime4Δ* (**Supplementary Figure S5c**). Interestingly, in response to pheromone, more genes were downregulated in the *ime4Δ* strain compared to the wild-type strain (**Supplementary Figure S5d**), a trend also observed with the *kar4Δ* strain (**Figure 1b**). Importantly, however, there were almost no substantial gene expression differences between wild-type and *ime4*Δ cells at ≥ 2-fold (for protein-coding genes and excluding *IME4* itself) before or after pheromone treatment. Exceptions were *AFB1* and *BSC1*, which were upregulated slightly more than 2-fold higher in response to pheromone in the *ime4*Δ strain, and *YGL088W*, which was downregulated slightly more than 2-fold more in the *ime4*Δ strain. Nonetheless, our results indicate that the effect of Kar4 on gene expression cannot be explained by its MTC-associated activity.

### Ste12 and Kar4 are predicted to form a continuous β-sheet

Although Kar4 behaves like a transcription factor, it lacks a clear DNA-binding domain (Kurihara *et al*. 1996; Lahav *et al*. 2007), and it has been hypothesized to function via an interaction with Ste12 (Kurihara *et al*. 1996). Further, Kar4 and Ste12 have been shown to interact both genetically (Lahav *et al*. 2007; Park *et al*. 2023c) and physically (Aymoz *et al*. 2018). However, there is a complete absence of structural data on Ste12 or Kar4 that can be used to hypothesize how Kar4 might modulate Ste12 function. To inform a mechanistic model on the physical relationship between Ste12 and Kar4, we used AlphaFold3 to model their interaction in the presence of DNA containing two Ste12-binding sites (also known as pheromone response elements, or PREs) (**Figure 3, Supplementary Figure S6, and Supplementary File S5**). Because much of Ste12 is a disordered region, we included only the DNA-binding domain (residues 21-220), which has been previously shown to be sufficient to bind PREs (Yuan and fields 1991). Related, Kar4 is expressed as two isoforms, and here we modeled only the Kar4-short isoform (residues 31-335), as only this isoform is transcriptionally induced by pheromone and would therefore be the primary isoform interacting with Ste12 (Gammie *et al*. 1999). Intriguingly, not only were these domains of Ste12 and Kar4 predicted with high confidence to form a dimer, but they were also predicted with high confidence to form an interchain β-sheet structure that stretches across the interface of the two proteins (**Figure 3a and b, and Supplementary Figure S6a and b**). Of note, the model shows only non-specific DNA binding, as expected from the current limitations of AlphaFold3. Importantly, function-specific Kar4 mutations have been previously found that disrupt its interaction with Ste12 and mating function, without affecting its meiotic function (Park *et al*. 2023c). These mutations were nearly all localized to one side of the β-sheet in Kar4, suggesting that they destabilize the dimerization interface (**Figure 3a and c**). In contrast, mutations that specifically disrupted Kar4’s meiotic function localized to peripheral surfaces unrelated to the β-sheet (**Figure 3b and c**). These results suggest that Kar4 is likely to affect transcription via a direct physical interaction with Ste12, which may then modulate Ste12 DNA-binding affinity, and/or its interaction with other proteins.

**Figure 3.**
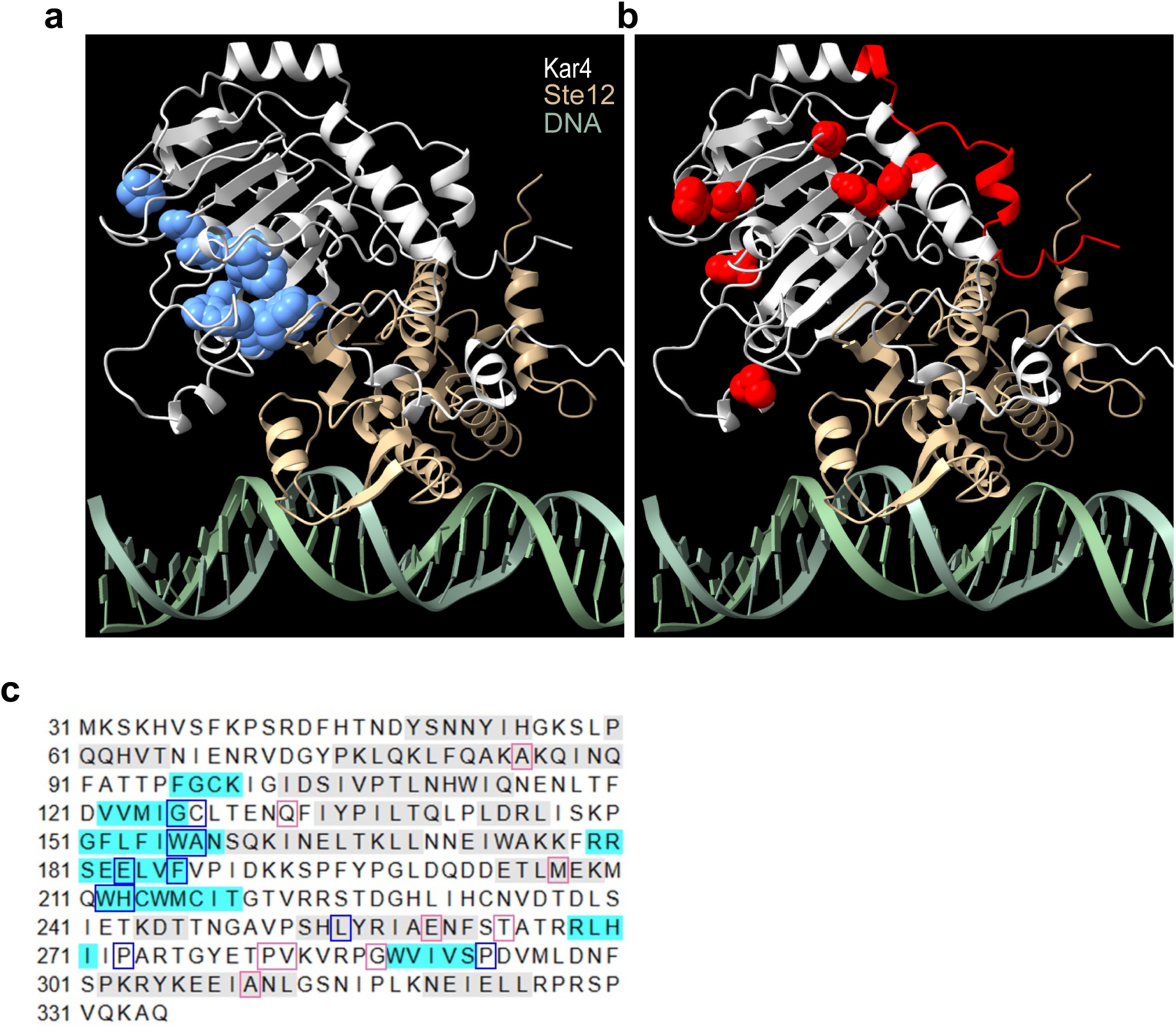
Ste12 and Kar4 are predicted to form a continuous β-sheet. a) Predicted Kar4 (residues 31-335; silver) and Ste12 (residues 21-220; gold) dimer structure by AlphaFold3 in the presence of DNA (green). Mating-defective alleles are shown in blue. b) As in panel a, but meiosis-defective alleles are shown in red. c) Modeled Kar4 protein sequence, with predicted **α**-helices highlighted in grey and predicted **β**-strands in blue. Mating-defective alleles are outlined in blue and meiosis-defective alleles in red.

### Kar4 is present at most of the chromosomal loci bound by Ste12

To further understand the role of Kar4 in Ste12 function, we directly examined whether Kar4 affects the chromosomal sites to which Ste12 binds, both in the presence and absence of pheromone. To this end, we analyzed a dataset created in parallel assessing the chromatin binding of Ste12-TAP and Kar4-TAP during either asynchronous log-phase growth or following 90 minutes of pheromone treatment. The analysis was done using wild-type (for Ste12-TAP and Kar4-TAP) or *kar4*Δ (for Ste12-TAP only) strains (**Supplementary File S6**; note that final sample depth can vary considerably between samples and can affect the final number of peaks detected). DNA association was assessed using ChIP-exo, where the protein of interest is crosslinked to the chromatin, and the chromatin is sheared and subjected to immunoprecipitation (ChIP). This is followed by a 5’ to 3’ exonuclease digestion of the DNA up to the position where a crosslinked protein (e.g., Ste12-TAP) blocks further digestion, enabling higher base-pair resolution of protein-DNA binding over traditional ChIP-Seq (Rhee and pugh 2011; Rossi *et al*. 2018). Of note, because the RNA-seq and ChIP-exo datasets were independently generated, the experimental designs used with the two approaches differ in two major ways: (a) the ChIP-exo and RNA seq experiments used different yeast strain backgrounds, and (b) The ChIP-exo experiment used a 90-minute pheromone treatment of asynchronous log-phase cells (similar to Roberts et al. (2000) and Lahav et al. (2007)), while the RNA-seq results were generated using a 60-minute pheromone treatment of cells pre-synchronized in G1. However, as most of the strongly upregulated genes in our RNA-seq dataset matched the response seen in previous asynchronous experiments (Roberts *et al*. 2000) (**Supplementary Figure S3**), general conclusions regarding Kar4 and Ste12 should be transferable.

Around half of Ste12-TAP and Kar4-TAP binding sites observed during the pheromone response in wild-type cells were in promoter regions (within 500 bp of a feature start site; 333 out of 601 sites for Ste12-TAP and 596 out of 1169 sites for Kar4-TAP) (**Supplementary File S7**, see “closestdist” column). We then associated each site with the nearest gene(s) or long-terminal repeat(s) (LTR) (see Methods) (**Supplementary File S7**, see “likelycat” column). Example ChIP-exo data are shown for promoters of two pheromone-induced genes in **Figures 4a and b** (all samples, including *kar4*Δ and Kar4-TAP, shown in **Figures S7a and b)**. The rank order within biological replicates was highly reproducible (**Supplementary Figure S8a**), typically having rank-based correlation values > 0.8 (**Supplementary Figure S8b**). Because LTRs are essentially copies of each other (with minor sequence variations), we exclude these from further analyses unless otherwise stated. After removing LTRs, Ste12-TAP was associated with 207 genes before pheromone treatment and 399 genes after pheromone treatment (**Figure 4c**) (note that some genes (∼25%) are associated with more than one Ste12 binding site). Of note, not all genes that were bound by Ste12 were transcriptionally induced by pheromone under our experimental conditions. Conversely, not all genes that were induced transcriptionally by pheromone were bound by Ste12. This is common in ChIP studies, and here we consider it likely to be because Ste12 also binds DNA with non-pheromone-induced cofactors (in the former case) and because some genes are induced indirectly by Ste12 (in the latter case), although differences in experimental conditions may contribute as well (see Discussion for more explanations).

**Figure 4.**
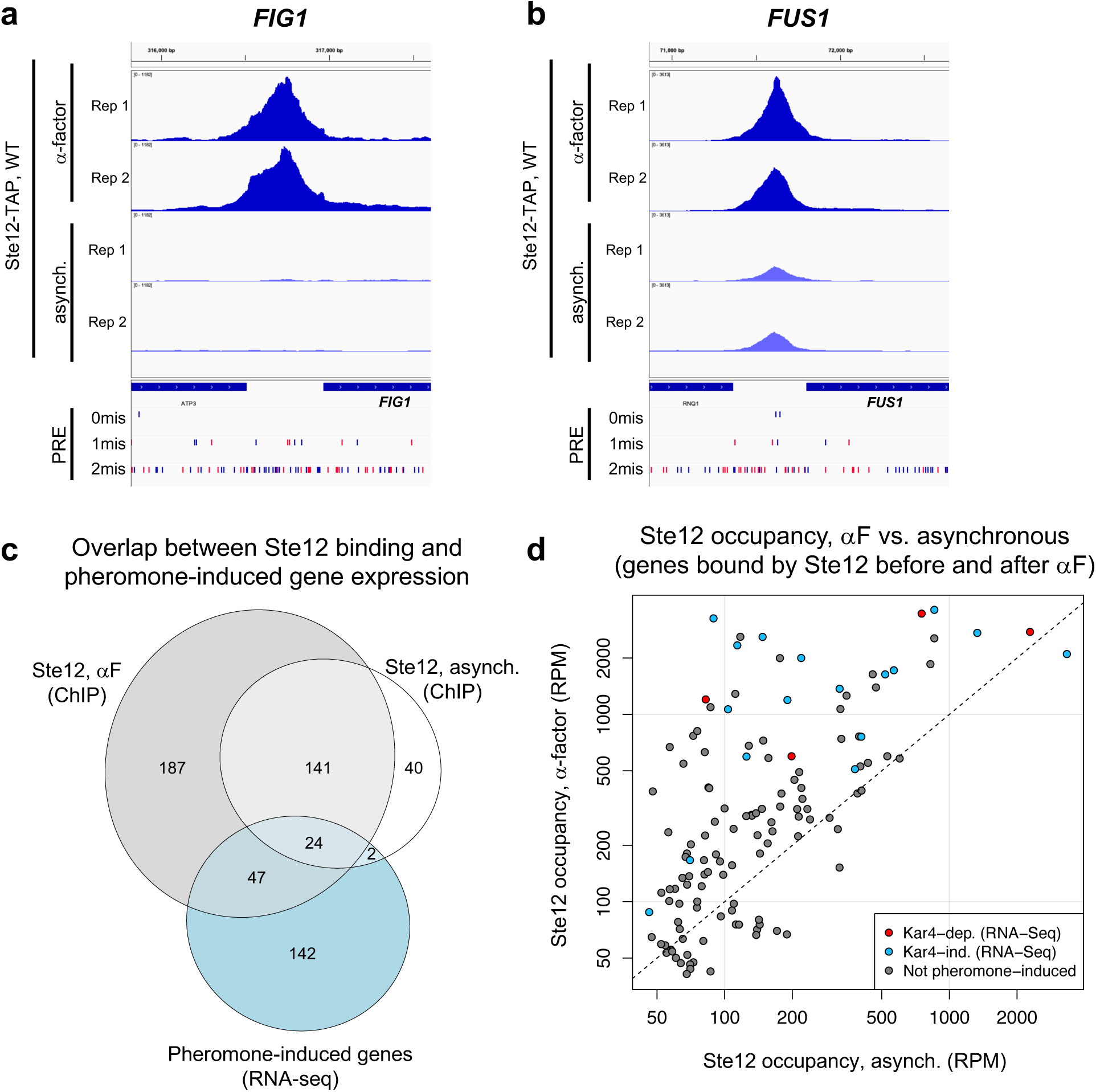
Ste12’s DNA association increases in response to pheromone. a) ChIP-exo data for Ste12-TAP in wild-type cells at the *FIG1* locus, before (bottom two tracks) or after (top two tracks) pheromone treatment. Samples are depth-normalized and thus peak height values are directly comparable between tracks. Ste12 binding is only evident after pheromone treatment. PREs (TGAAACA) containing 0, 1, or 2 mismatches are displayed in the bottom 3 tracks, with matches on the forward strand in blue and reverse strand in red. Data for *kar4*Δ and Kar4-TAP are shown in **Supplemental Figure S7a**. b) As in panel a, but for the *FUS1* locus, which is bound by Ste12-TAP both before and after pheromone treatment. All conditions are shown in **Supplemental Figure S7b**. b) Venn diagram indicating the overlap between all genes associated with Ste12 binding before (white) or after (grey) pheromone, as well as genes induced by pheromone in our RNA-seq dataset (blue). d) Normalized peak heights in reads per million (RPM) for genes bound by Ste12 both before and after pheromone. Red and blue points are pheromone-induced and Kar4-dependent or Kar4-independent by RNA-seq, respectively. Grey points are not pheromone-induced transcriptionally in our dataset. Dashed line indicates 1:1 line for equal values. Note that axes are spaced logarithmically; absolute unit values are displayed.

Kar4 has previously been shown to physically interact with Ste12 by co-IP (Aymoz *et al*. 2018) and one-hybrid (Lahav *et al*. 2007), and to bind to many of same sites as Ste12 by ChIP (Rossi *et al*. 2021) during log-phase growth, without pheromone treatment. Our ChIP-exo analysis revealed that 86% (after pheromone) and 46% (before pheromone) of all Ste12-bound genes are also associated with Kar4 (**Supplementary Figure S9a**). This was somewhat unexpected given that in the presence of pheromone, the induced expression of less than 25% of genes is affected transcriptionally when deleting *KAR4* (**Supplementary Figure S4a and b**). Our data uncovered more genes that were associated with Kar4 binding (775 in pheromone) than Ste12 binding (399 in pheromone; with 344 genes in common) (**Supplementary Figure S9a**). However, we note that this is not necessarily due to Kar4 binding DNA independently of Ste12, but likely reflects technical differences such as better read depth for Kar4 (**see Supplementary File S6**) and possibly a better signal/noise ratio resulting from a better immunoprecipitation efficiency for Kar4-TAP. Consistent with this result being a technical difference, a direct comparison between Ste12 and Kar4 occupancy during pheromone treatment revealed only 32 genes associated with significantly higher Kar4 occupancy than Ste12, of which only 20 were not associated with Ste12 binding at all (**Supplementary File S8**, compare to list in **Supplementary File S3** or **S7**). Moreover, most of the sites bound by Kar4 but not Ste12 contain a Ste12-binding DNA motif found at other Ste12 binding sites (see below). This suggests that nearly all of the binding sites uniquely identified for Kar4 were low-occupancy or were noisy peaks for Ste12, with some binding also observed for Ste12 even though the binding was not significant when measured for Ste12 alone. Therefore, Kar4 likely binds DNA through Ste12, as also concluded by Aymoz et al. (2018).

### Ste12’s and Kar4’s DNA association increases in response to pheromone

Qualitatively, many Ste12 binding sites exhibited increased Ste12 occupancy after pheromone treatment (for examples, see **Figure 4a and b**; see the *TEC1* locus in **Supplementary Figure S7c** for a counter-example). For 60% of Ste12-associated genes (234/399), Ste12 was not detected before pheromone treatment (**Figure 4a and c**). For 40% of Ste12-associated genes (165/399), Ste12 was detected both before and after pheromone treatment (**Figure 4b and c**). In these genes, Ste12 occupancy (peak height) increased significantly in 35% (57/165) after pheromone treatment (**Figure 4d** and **Supplementary File S8**, compare to genes bound before and after pheromone in **Supplementary File S7**). Similar to Ste12, we found that for the 223 genes associated with Kar4 binding both before and after pheromone treatment, 44% (98/223) exhibited a statistically significant increase in Kar4 occupancy (**Supplementary Figure S9b** and **Supplementary File S8**). Interestingly, the pheromone-induced increase in occupancy was not as great for Kar4 as it was for Ste12; comparing genes with significantly increased occupancy for both Ste12 and Kar4, the increase in binding was higher for Ste12 by 2.5-fold on average (observed in 88%, 139/158 of genes) (**Supplementary Figure S9c**). This could reflect the stoichiometry of Ste12 and Kar4 in the DNA-bound complex, or it could represent a mixture of Ste12-bound complexes with and without Kar4 in different cells (see Discussion). Nonetheless, these results demonstrate an increased DNA association for both Ste12 and Kar4 following pheromone treatment, even for sites where binding is observed prior to pheromone addition.

We next compared the ChIP-exo and RNA-seq datasets. Of the 207 genes associated with Ste12 binding before pheromone treatment, 26 were identified by our RNA-seq analysis as having a 2-fold or greater pheromone-induced increase in gene expression (**Figure 4c**). After pheromone treatment, 71 of the 399 genes associated with Ste12 binding exhibited a ≥ 2-fold increase in pheromone-induced gene expression (**Figure 4c**). For the large majority of genes that were associated with Ste12 binding and induced ≥2-fold transcriptionally by pheromone, Ste12 occupancy increased following pheromone treatment (23 out of 26 genes with Ste12 bound prior to pheromone treatment (**Figure 4d**); or 63 out of 71 of genes with Ste12 bound after). However, the correlation between the fold-change in Ste12 occupancy and the fold-change in transcript levels was modest (R = 0.44; **Supplementary Figure S10**). Nevertheless, these results demonstrate that nearly all transcriptionally pheromone-induced genes associated with Ste12 binding exhibit increased Ste12 occupancy following pheromone treatment.

### Kar4 promotes Ste12’s DNA association at most transcriptionally pheromone-induced genes, even those that are transcriptionally Kar4-independent

To address the functional role of Kar4 in Ste12’s DNA association, we examined Ste12 binding in a *kar4Δ* strain. Based on our RNA-seq analysis, we anticipated that Ste12 binding would be largely unchanged in *kar4*Δ cells, except at a small subset of Kar4-dependent gene promoters such as *CIK1* and *KAR3*, and at novel loci not bound in wild-type, such as at promoters of genes expressed only in *kar4Δ* (namely the *kar4*Δ-only category). As expected, Ste12 binding was evident at both the *CIK1* and *KAR3* promoters after pheromone treatment in wild-type but not *kar4*Δ cells (**Figure 5a and Supplementary Figure S11**). However, while the number of genes associated with Ste12 binding after pheromone treatment in wild-type (399) was similar to the number of genes in *kar4*Δ cells (436), the overlap was only 261 genes (**Supplementary Figure S12**). This observation implies that, as for RNA-seq, there are Kar4-independent, Kar4-dependent, and *kar4*Δ-only Ste12 binding sites. Comparing the ChIP-exo and RNA-seq results in wild-type cells, out of the 399 gene promoters bound by Ste12 after pheromone treatment, 52 genes were transcriptionally induced in a Kar4-independent manner and 18 genes were induced in a Kar4-dependent manner (**Figure 5b**).

**Figure 5.**
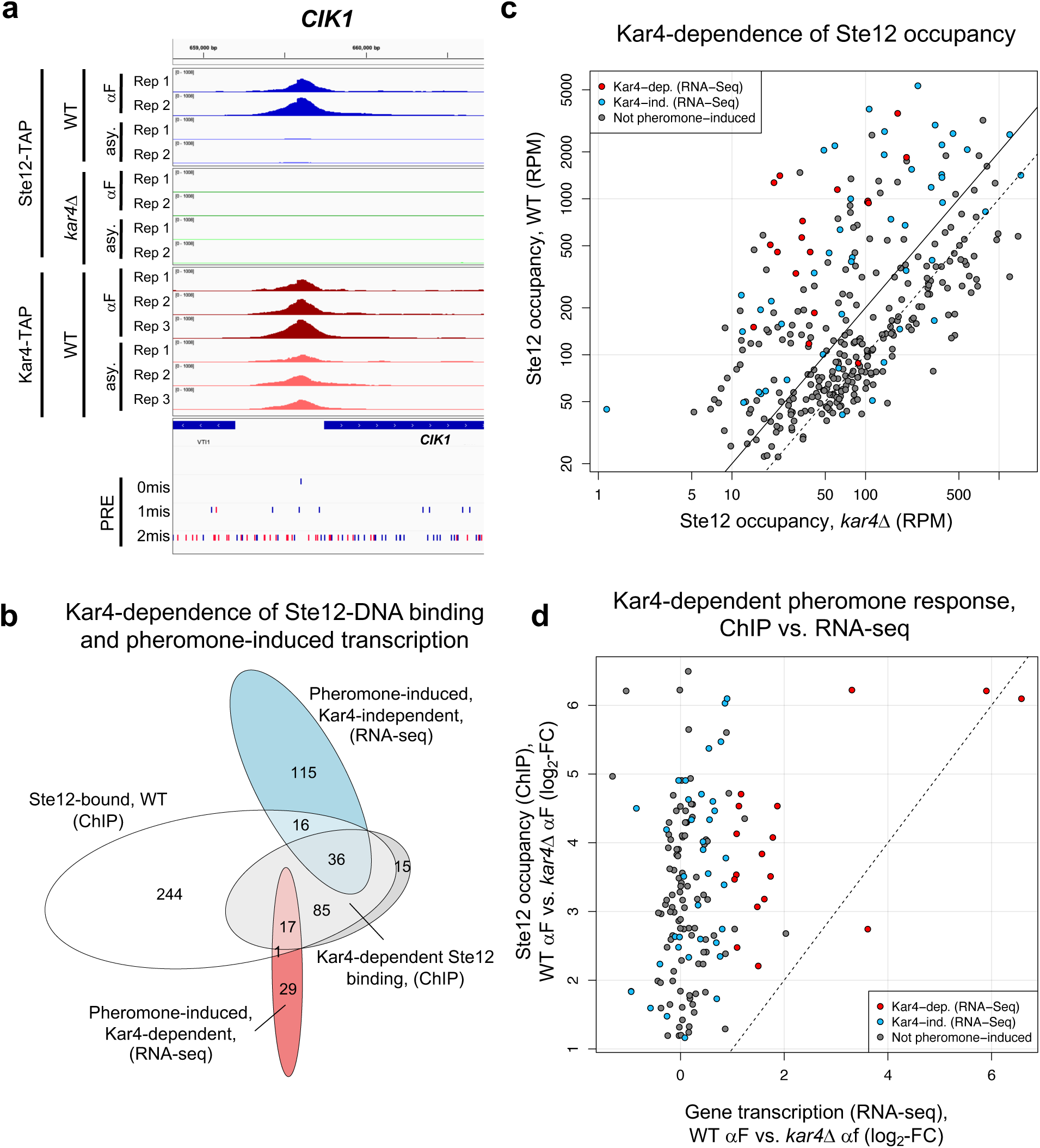
Kar4 promotes Ste12’s DNA association at most transcriptionally pheromone-induced genes, even those that are transcriptionally Kar4-independent. a) Example Ste12-TAP (top 8 tracks) and Kar4-TAP (bottom 6 tracks) binding at the *CIK1* locus by ChIP-exo, in wild-type and *kar4Δ* cells (only of Ste12-TAP in the latter), in asynchronous cells (“asy”) and after pheromone treatment (“αF”). PREs are as in Figure 4a. b) Venn diagram indicating the overlap between genes associated with Ste12 binding (ChIP data) after pheromone (white), genes associated with significant Kar4-dependent Ste12 binding after pheromone (grey), and genes induced by pheromone (RNA-seq data) in a Kar4-independent (blue) or Kar4-dependent (red) manner. Note that the Kar4-dependent gene set (grey) is not strictly a subset of all Ste12 binding sites (white) because the significance tests are performed independently (see Methods). c) Normalized Ste12 occupancy as in Figure 4d, comparing occupancy in wild-type and *kar4*Δ cells for all genes associated with significant Ste12 binding during pheromone treatment in wild-type. Dashed line indicates 1:1 line. Solid line indicates 2-fold change in WT Ste12 occupancy. Red and blue points are pheromone-induced and Kar4-dependent or Kar4-independent by RNA-seq, respectively. Grey points are not pheromone-induced transcriptionally in our dataset. d) Quantification of Kar4-dependence for ChIP-exo vs. RNA-seq. Points are colored as indicated (and the same as in panel c), only genes associated with significant Kar4-dependence by ChIP are shown.

To determine how Kar4 affects Ste12 binding, we examined the difference in Ste12 occupancy between wild-type and *kar4*Δ cells (**Supplementary File S8**). The effect of Kar4 was greater on the binding of Ste12 at genes that were transcriptionally induced by pheromone. Kar4-dependence (defined as significantly higher occupancy at the same site in wild-type vs. *kar4*Δ cells after pheromone treatment) was observed for 38% of all Ste12-bound genes (153/399), rising to 76% of transcriptionally pheromone-induced genes (54/71; this includes the promoter of *KAR4*) **(Figure 5c**, red and blue circles). Of the 17 pheromone-induced genes that were Kar4-independent for Ste12-binding (**Figure 5b**; 16 + 1), 16/17 were also Kar4-independent for transcription as measured by RNA-seq (**Figure 5b**). The role of Kar4 in promoting Ste12 binding was not restricted to the mating response, as 19 genes were also significantly Kar4-dependent for Ste12 binding during asynchronous growth (**Supplementary File S8**), although this is only a small number compared to the total number of Ste12-associated genes prior to pheromone (207). Taken together, these results demonstrate that Kar4 promotes Ste12 binding to the majority of Ste12-binding sites associated with genes that are transcriptionally induced by pheromone.

We were surprised to find that most genes that were transcriptionally induced by pheromone were also Kar4-dependent for Ste12 binding, as only a minority of these genes required Kar4 for normal pheromone-induced gene expression (18 out of 70, **Figure 5b**). To examine this discrepancy, we compared the fold-change in Ste12 occupancy against the fold-change in transcript abundance for wild-type vs. *kar4*Δ cells for all genes exhibiting Kar4-dependent Ste12 binding (**Figure 5d**). We focused on this set of genes rather than all Ste12-bound genes because most genes that were transcriptionally affected by pheromone were contained within it. Overall, the effect of Kar4 on the fold-change of Ste12 occupancy tended to be much larger than its effect on the fold-change of gene expression, with the two being weakly but significantly correlated (R = 0.29; p-val = 3 x 10^−4^). Restricting the analysis to genes which were pheromone-induced at least 2-fold in wild-type by RNA-seq (blue and red points in **Figure 5d**) increased the correlation to 0.46 (p-val = 5 x 10^−4^), with little difference between genes that were Kar4-dependent or Kar4-independent transcriptionally. Therefore, while there is a correspondence between the Kar4-dependence measured by ChIP-exo and RNA-seq, the correlation is weak and the effect of Kar4 on Ste12 binding is greater than anticipated based on the RNA-seq data alone.

### Ste12 binds a PRE di-motif with 4 nucleotide spacing and head-to-tail orientation in at least 44% of sites associated with pheromone-induced transcription

Ste12 is known to bind a pheromone response element (PRE), optimally 5’-TGAAACA-3’, both in vitro and in vivo (Kronstad *et al*. 1987; Hagen *et al*. 1991; Yuan and fields 1991). Further, Ste12 has been shown to exhibit cooperative binding between a pair of PREs, suggesting that DNA binding requires two adjacent Ste12 proteins, widely assumed to dimerize (Yuan and fields 1991). However, attempts to identify a canonical PRE-containing motif in pheromone-induced gene promoters has revealed a multitude of spacing and orientation configurations (possible configurations shown in **Figure 6a** and **Supplementary Figure S13)**, with apparent functional redundancy in vivo between multiple PREs, including suboptimal PREs, within promoters (Chou *et al*. 2006; Su *et al*. 2010; Aymoz *et al*. 2018). It should be noted that PREs with 1, 2, or 3 mismatches are common across the genome, with promoters often exhibiting multiple candidate PREs (see, for example, the density of PREs with 2 mismatches in **Figure 4a and b**, and **Figure 5a**). Given the higher DNA binding resolution of ChIP-exo, we endeavored to identify a favored PRE di-motif configuration for Ste12-binding in vivo, and the extent to which binding might be affected by Kar4.

**Figure 6.**
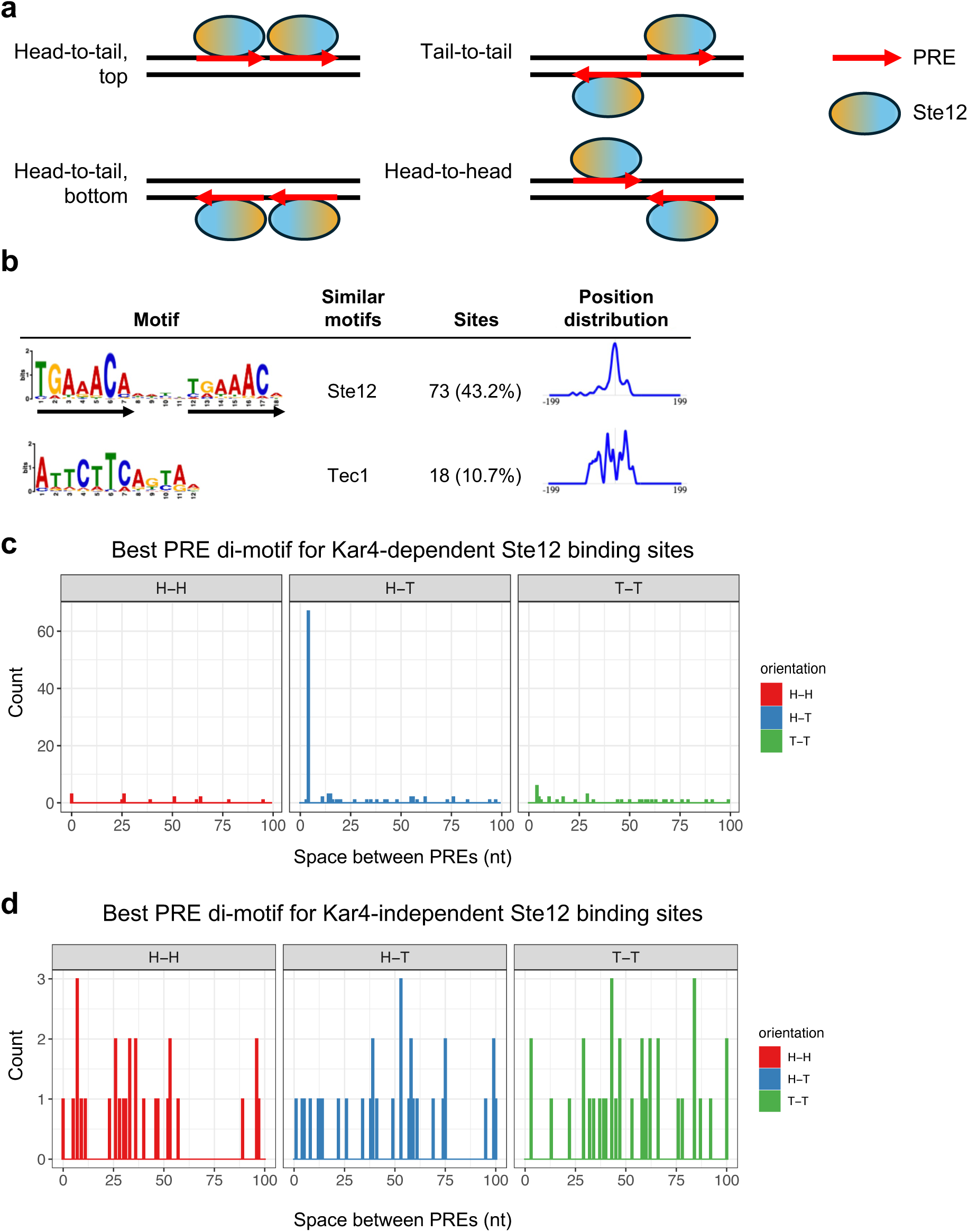
Ste12 binds a PRE di-motif with 4 nucleotide spacing and head-to-tail orientation in at least 44% of sites associated with pheromone-induced transcription. a) Possible PRE di-motif configurations. Arrows are in the 5’ to 3’ direction of the PRE (5’-TGAAACA-3’). Head and Tail indicate the head and tail of the arrow, or the 3’ and 5’ ends of the PRE, respectively. Top and bottom indicate the DNA strand as it appears in the reference genome. The orange and blue ends of the schematic Ste12 are always oriented in the 5’ to 3’ direction. b) Top two hits from a de novo motif analysis (STREME) for Kar4-dependent Ste12 binding regions (LTR-associated sites excluded). Two more hits with similar enrichment to Tec1 not shown for simplicity (potential Azf1 and Mig1-associated motifs). Similar known motif column is from Tomtom. Arrows indicate PRE locations. Position distribution is relative to center of queried sequenced. c) Histogram indicating the number of the best PRE di-motif match with the indicated orientation and spacing (up to 100 nucleotides), using all Kar4-dependent, ChIP-exo non-LTR Ste12-binding sites (n = 151). d) As in c, but for Kar4-independent Ste12-binding sites (n = 94).

We initially performed *de novo* DNA motif discovery on the 121 bp genomic DNA region centered on each Ste12 binding site from ChIP-exo, allowing for a motif size of 6 to 24 nucleotides (nt). As Ste12 is known to bind cooperatively at some sites with other transcription factors such as Tec1 or Mcm1 (Oehlen *et al*. 1996; Wong sak hoi and dumas 2010; Cullen and sprague 2012), we initially restricted the analysis to 169 non-LTR-associated sites exhibiting Kar4-dependent Ste12 binding in the presence of pheromone. The most enriched motif was a PRE di-motif with 4 nt spacing and a head-to-tail (H-T) orientation (hereafter referred to as an H-T 4 motif), identified in at least 43% (73/169) of sequences (**Figure 6b** and **Supplementary File S9**). We also identified a slight enrichment for Tec1-binding sites (11%; 18/169), despite only analyzing Kar4-dependent binding sites.

Although Ste12 was expected to bind to a pair of PREs, we were surprised that almost half of all binding sites use the same orientation and spacing. Previous studies suggested that pheromone-induced promoters have a wide variety of PRE orientations and spacings (Su *et al*. 2010), with a preference for a tail-to-tail orientation with 3 nt spacing in vitro, and possibly in vivo (Dorrity *et al*. 2018; Pinheiro *et al*. 2025). Therefore, we reasoned that better di-motifs may exist in most promoters, but because they lack a conserved orientation or spacing they do not show up as enriched by common motif discovery algorithms (MEME or STREME). To test this possibility, we scanned a larger genomic DNA region (241 bp centered on the binding site, 151 regions overall) for all possible PRE di-motifs, ranging from 0 to 100 nt spacing and any orientation (H-H, H-T top and bottom, and T-T). We then assigned each region a “best” motif by ranking all motifs by q-value, which is largely determined by the number of mismatches, independent of position (from FIMO, see Methods). Using this independent method, we again identified the H-T 4 motif as the best match in 44% (67/151) of Ste12-binding regions (**Figure 6c**). We further identified an H-T 4 motif in the top 5 best possible PRE di-motifs (as ranked by FIMO q-value) in 68% of Kar4-dependent Ste12-binding regions, suggesting that Kar4 may preferentially direct Ste12 to H-T 4 motifs (see Discussion). The H-T 4 motifs had no bias with respect to direction of the gene start site (**Supplementary Figure S14**), implying that Ste12 binding to the H-T 4 motif functions independently of the orientation, and thus has the potential to activate transcription bidirectionally. The remaining Kar4-dependent Ste12-binding sites had more variable best-match PRE di-motifs with respect to orientation and spacing, with the most common motifs being T-T 4 (6 sites), H-T 14 and H-T 15 (3 sites each), and H-H 0, H-H 26, and T-T 29 (3 sites each) (**Figure 6c**).

To explore the functional importance of the H-T 4 motif for transcription, we used a program (Transfactivity) that quantifies how well a motif can predict a set of gene expression fold-changes, given their promoter sequences (Bussemaker *et al*. 2001; Roven and bussemaker 2003; Lee and bussemaker 2010). Unlike motif enrichment algorithms, which find motifs enriched in a set of DNA sequences, this program uses given motifs to predict gene expression fold-changes, given a gene expression dataset, and is therefore an orthogonal method for assessing motif importance. We used Transfactivity on our RNA-seq dataset together with a collection of known budding yeast transcription factor DNA motifs (see Methods). Motifs associated with Ste12 and Crz1 were the strongest predictors of pheromone-induced gene expression (**Figure 7a**; see Appendix for more details and discussion of Crz1). We additionally ran a second analysis comparing the predicted activity of the H-T 4 motif derived from MEME (**Figure 6b**) to that of the standard PRE mono-motif (TGAAACA) already associated with Ste12 binding. For both motifs, transcriptional activity was significantly reduced in the *ste12Δ* strain, both with and without pheromone (**Figure 7b and c**), as expected for promoter regions that are regulated by Ste12. Likewise, for both motifs transcriptional activity was significantly increased with the presence of pheromone. However, only the H-T 4 motif showed a dependence on Kar4, exhibiting a ∼50% reduction in activity in *kar4*Δ cells compared to wild-type in the presence of pheromone (**Figure 7c**). Intriguingly, activity of the H-T 4 motif also declined in *kar4*Δ cells prior to pheromone treatment, suggesting that this motif contributes to some Ste12 basal activity in a Kar4-dependent manner. As Ste12 is not thought to activate transcription via PRE mono-motifs, it is likely that the H-T 4 motif is more physiologically relevant. Together, these results suggest that Kar4 promotes Ste12 binding to the H-T 4 motif, including in most of the transcriptionally pheromone-induced gene promoters.

**Figure 7.**
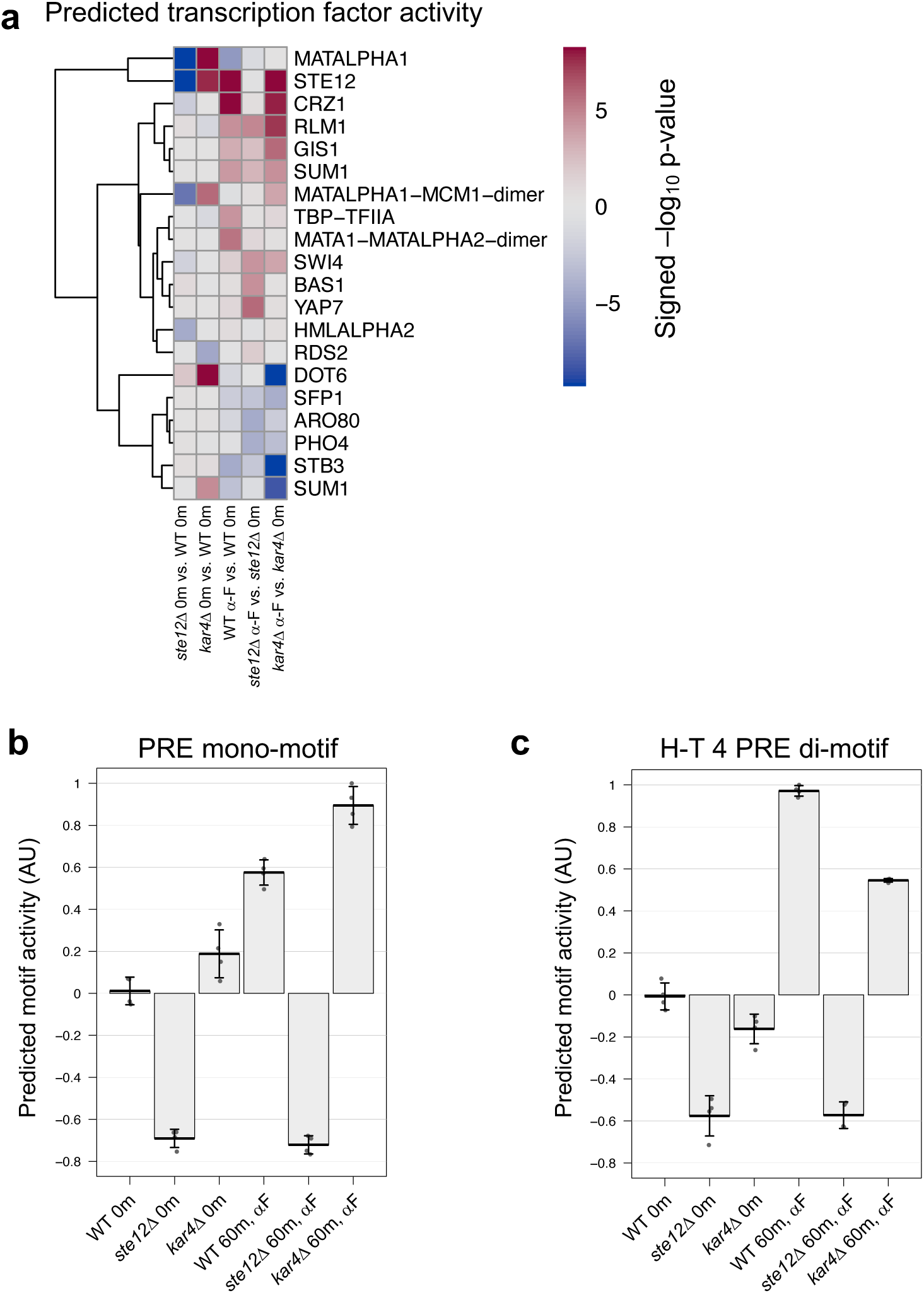
Kar4 function is linked to H-T-4-motif-associated transcriptional activity. a) Predicted transcription factor activities (-log_10_ p-value, with red indicating the transcription factor motif is associated with upregulated genes, and blue indicating an association with downregulated genes) based on Transfactivity (see Methods). Input gene expression fold-changes were derived from the indicated contrasts from DESeq2 for all genes (**Supplementary File S2**). “Signed” indicates that the direction of the fitted score value (negative or positive) was assigned to the p-values. Only transcription factors with at least 1 p-value under 1 x 10^−4^ are shown. b) Predicted motif activity based on Transfactivity in indicated samples for the PRE mono-motif. The full log_2_ fold-change gene expression matrix for all samples used in Figure 2 was used as input. Units are derived from fitted score values (coefficients) and rescaled such that the largest absolute value is 1. Error bars indicate standard deviation. n = 4 samples per group. c) Predicted H-T 4 motif activity in indicated samples, as in panel b.

### Kar4-independent Ste12 binding primarily occurs via cofactor interactions and is not enriched for any PRE di-motif configuration

Many Ste12 binding sites identified after pheromone treatment were Kar4-independent. A priori, these sites could include Ste12 binding to a PRE di-motif that did not require Kar4’s involvement, Ste12 binding mediated through another cofactor (e.g. Mcm1, Tec1, etc.) such that the binding site would include only a single PRE, or a spurious Ste12-DNA association due the pheromone-induced activation of Ste12. To better characterize Kar4-independent Ste12 DNA binding, we looked at the transcriptional activation of these genes in response to pheromone, the extent and induction of Ste12 binding, the nature of the PREs and di-motifs associated with these binding sites, and whether there were sites for other Ste12 cofactors.

In total we identified 291 Kar4-independent Ste12-binding sites after pheromone treatment. Many peaks had low Ste12 occupancy (potentially false positives) and we therefore restricted the analysis to 111 high-confidence peaks (see Methods). These 111 peaks were associated with 117 genes, of which most did not change expression after pheromone treatment (**Supplementary Figure S15a**). Specifically, only 17 out of 117 genes exhibited increased expression upon pheromone treatment. Further, most of these 17 genes contained both Kar4-independent and Kar4-dependent Ste12 binding sites in their promoters, suggesting that even fewer than 17 of the Kar4-independent Ste12-binding sites are functionally important for pheromone-induced transcription. We considered that the low overlap may be due to experimental differences between our RNA-Seq and ChIP-exo datasets, and we therefore also compared our data to Roberts et al. (2000), who measured changes in gene expression after pheromone treatment in asynchronous cells (similarly to our ChIP-exo experimental design). Despite being more comparable experimentally, we still only observed a small overlap with transcriptionally pheromone-induced genes (21/117), with an overlap of 10 genes between all three datasets (**Supplementary Figure S15a**). Therefore, it is likely that very few, if any, of these Ste12 binding sites lead to increased transcription after pheromone treatment.

Given that most of these sites were not associated with increased transcription after pheromone treatment, we examined the levels of Ste12 occupancy before and after pheromone treatment. Surprisingly, the majority of these sites nevertheless exhibited increased Ste12 occupancy after pheromone treatment, although to a lesser degree than for Kar4-dependent sites (**Supplementary Figure S15b**).

Next, we asked whether the Kar4-independent Ste12-binding sites share a common PRE configuration or another non-PRE DNA motif. Direct di-motif scanning revealed no PRE di-motif bias in the Kar4-independent Ste12-binding sites (**Figure 6d**), including in sites associated with transcriptionally pheromone-induced genes (**Supplementary Figure S15c**). Moreover, even when considering the top 5 best possible PRE di-motifs as above, only 6% of sites contained an H-T 4 motif (compared to 68% for Kar4-dependent Ste12 binding sites). However, perfect PREs were fairly common, found in 22% of the best di-motifs in Kar4-independent Ste12-binding regions (21/94; 241 bp width) compared to 36% of the best di-motifs in (54/151) Kar4-dependent regions, consistent with these being genuine Ste12-binding sites. De novo motif analysis revealed a strong enrichment for the Mcm1 motif (∼30% of sites), a known Ste12-interacting cofactor, as well as weak enrichment for two motifs resembling the PRE (aGAAACg (17% sites) and gGAAAtn (25% sites); **Supplementary Figure S15d**). Additionally, although it was not discovered by the de novo analysis, some of these sites likely arise from Ste12-Tec1 binding at PRE-TCS (Tec1-binding site) di-motifs, as the strongest Kar4-independent Ste12 binding peak is at a known PRE-TCS di-motif in the *TEC1* promoter itself (Madhani and fink 1997). Therefore, most Kar4-independent Ste12-binding sites likely arise from combinatorial Ste12 binding with other DNA-binding proteins for processes unrelated to the pheromone response.

### Genes that are Kar4-dependent for transcription are enriched for suboptimal Ste12-bound PREs

One of our initial goals was to determine the underlying difference between Kar4-independent and Kar4-dependent pheromone-induced transcription. Note that these two classes are different from Kar4-dependent vs. Kar4-independent Ste12 binding, which was discussed in the previous section (**Figure 5b**). By ChIP-exo, the Kar4-dependent and Kar4-independent transcriptional gene classes are remarkably similar; promoters of both gene classes are bound by Ste12, show increased Ste12 occupancy after pheromone treatment (**Figure 4d**), and are mostly Kar4-dependent for Ste12 binding (**Figure 5c**). In addition, the most common Ste12 binding motif in both groups is the H-T 4 motif, found as the best PRE di-motif near a Ste12 binding site in 30% of genes whose expression is Kar4-independent (**Supplementary Figure S16a**) and 52% of Kar4-dependent ones (**Supplementary Figure S16b**; **Supplementary File S9**).

Further analyses, however, revealed a subtle difference in the H-T 4 motifs in the two groups. There is a strong inverse correlation between the number of PRE mismatches and the level of Ste12 occupancy at sites where Ste12 is bound in a Kar4-dependent manner (**Figure 8a**). Overall, 51% of the Ste12-bound di-motifs at transcriptionally Kar4-independent genes contained at least one perfect PRE sequence, whereas only 17% of the di-motifs at Kar4-dependent genes contained a perfect PRE (**Figure 8b**). Conversely, Kar4-dependent genes were enriched for sites with mismatches in both PREs (**Figure 8b**). Given that the affinity of Ste12 to a PRE decreases as the number of mismatches increases (Su *et al*. 2010), PREs found in transcriptionally Kar4-dependent gene promoters likely have a lower affinity for Ste12, and thus require Kar4 for efficient binding.

**Figure 8.**
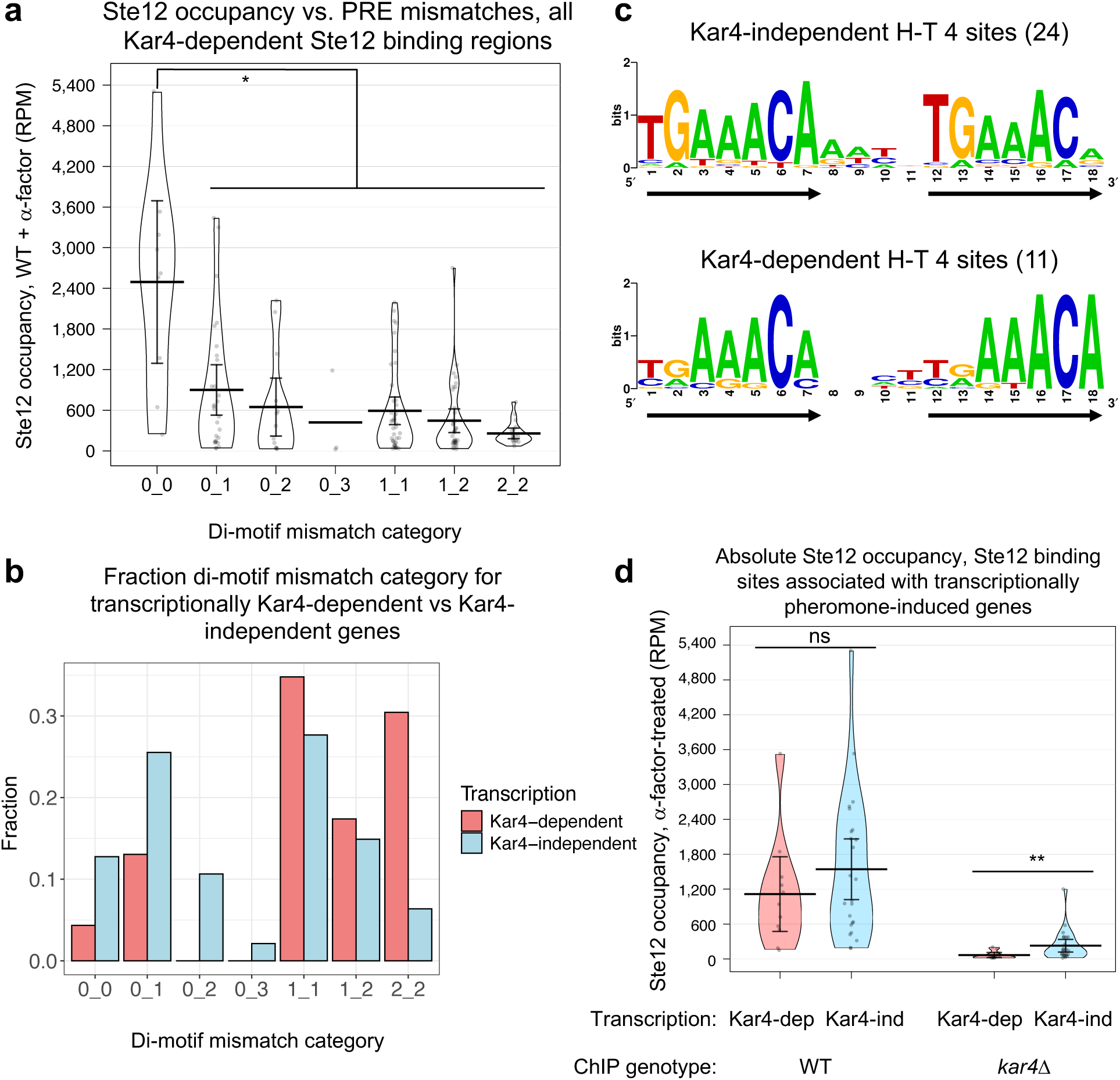
Genes that are Kar4-dependent for transcription are enriched for suboptimal Ste12-bound PREs. a) Ste12 occupancy (RPM) distribution as a function of di-motif mismatch categories. All Kar4-dependent Ste12-binding regions (n = 151, as in Figure 6c), including both H-T 4 and non H-T 4 di-motifs, were used. Di-motifs were categorized by the number of mismatches in each PRE (0_0 being no mismatches in either, 0_1 being a perfect PRE and a 1-mismatch PRE, 1_2 being a 1-mismatch PRE and a 2-mismatch PRE, etc.). Error bars indicate 95% confidence interval. All categories (except 0_3) are significantly different from 0_0 (Mann-Whitney U test, BH-corrected p-value < 0.05). All other comparisons are not significantly different. One data point is not shown (2_3 category, value of 30 RPM). b) Best-match PRE di-motifs (from **Figures S16a and b,** including both H-T 4 and non H-T 4 di-motifs,) were categorized by the number of mismatches in each PRE. The fraction of sites for each di-motif mismatch category was then calculated independently for sites associated with genes whose pheromone-induced expression was Kar4-dependent or Kar4-independent. c) Consensus H-T 4 motifs for manually-curated Ste12 binding sites associated with transcriptionally Kar4-independent (24 motifs) or Kar4-dependent (11 motifs) genes. Motifs were generated using WebLogo (Crooks *et al*. 2004). d) Ste12 occupancy (reads per million) for the indicated genotypes after pheromone treatment for the H-T 4 motifs associated with Kar4-dependent or Kar4-independent gene transcription (n = 35 per genotype, 24 Kar4-independent and 11 Kar4-dependent). Error bars indicate 95% confidence interval. For WT Kar4-dep vs. Kar4-ind, p = 0.29. For *kar4*Δ Kar4-dep vs. Kar4-ind, p = 0.0077. Statistical analyses were done using Mann-Whitney U test. See **Supplementary Figure S18a** for a y-axis-zoomed version.

Because the number of PRE mismatches, and hence Ste12’s binding affinity, appeared to be the primary feature that distinguished between genes that were pheromone-induced transcriptionally in a Kar4-dependent vs independent manner (**Figure 8b**), we further investigated differences in the mutational spectrum of the H-T 4 motifs found in each class. To do so, we first generated a list of high-confidence H-T 4 motifs with increased stringency. Using the Ste12 binding sites from **Figure 8b** as a starting point (23 sites associated with transcriptionally Kar4-dependent genes and 47 sites with Kar4-independent genes), we removed any sites associated with peaks which were not deemed high-confidence by visual inspection. This led to the removal of 16 “satellite peaks” associated with the shoulders of much stronger Ste12 binding sites, often inside transcribed regions, as well as 3 false-positive peaks associated with reads within the deleted *KAR4* gene (these sites were associated with the upstream gene *PEX34*, which is transcriptionally pheromone-induced and Kar4-independent). This left us with 14 and 37 sites associated with genes that were induced by pheromone in a Kar4-dependent or Kar4-independent manner, respectively (**Supplementary File S10**). Each of these sites unambiguously corresponded to a single high-confidence Ste12 binding peak, one per gene promoter. The mutational spectrum for these final 51 sites was qualitatively similar to the original result, except the 2_2 mutational category was less prevalent and the 1_2 mutational category was dominated by Kar4-dependent genes (**Supplementary Figure S17a**; compare to **Figure 8b**).

Within the final 51 sites, we identified any H-T 4 motifs. Some were already selected as the best possible PRE pair (9 and 13 H-T 4 motifs for transcriptionally Kar4-dependent and independent genes, respectively). We considered that some of the remaining sites may also harbor H-T 4 motifs which do not form the best possible PRE pair, but are nevertheless functionally important for Ste12 binding. For example, the *KAR3* promoter harbors a T-T 44 motif with 1 mismatch in each PRE near the Ste12 binding site, but also harbors an H-T 4 motif at the same region with 1 mismatch in the first PRE and 2 mismatches in the second PRE (**Supplementary Figure S17b**). In this instance, prior literature showed that the H-T 4 motif is sufficient for mediating Kar4-dependent transcription (Lahav *et al*. 2007), suggesting that the T-T 44 motif is fortuitous and non-functional (see Discussion). This was also generally evident from our results here, as the H-T 4 motif is by far the most prevalent motif for Kar4-dependent Ste12 binding sites, and, in many cases, the best-match motif is also a suboptimal H-T 4 motif, with 3 or even 4 mismatches total, implying such sites are functional. We therefore ultimately included H-T 4 motifs (one per site) that were near the Ste12 binding peak center (∼100 bp), with no more than 2 mismatches in either PRE. This increased the number of H-T 4 motifs for Kar4-dependent and independent genes to 11 and 24, respectively.

Consensus motifs for these two sets of H-T 4 motifs confirmed that the PREs associated with transcriptionally Kar4-independent genes are closer to the perfect PRE compared to the transcriptionally Kar4-dependent genes (**Figure 8c**). Specifically, in H-T 4 motifs associated with transcriptionally Kar4-independent genes, the first 2 nucleotides of the two PREs in the di-motif are conserved, and the same as the consensus PRE (**Figure 8c**, top row). In H-T 4 motifs associated with transcriptionally Kar4-dependent genes, the first two nucleotides are not conserved (**Figure 8c**, bottom row). Finally, the final nucleotide (position 18) was highly conserved only in the H-T 4 motifs associated with transcriptionally Kar4-dependent genes.

These results support the hypothesis that Kar4 facilitates Ste12 binding to suboptimal PREs. Specifically, when Kar4 is absent, such as in the *kar4Δ* strain, the expression of genes with suboptimal (low affinity) H-T 4 PREs will be affected to a greater extent than genes that have H-T 4 motifs with perfect or near-perfect PREs (high affinity). To examine whether this relationship correlates with a difference in Ste12 binding, we examined the effect of Kar4 on the amount of Ste12 associated with these 35 H-T 4 sites (11 and 24 associated with transcriptionally Kar4-dependent and Kar4-independent genes, respectively). In wild-type cells, Ste12 occupancy was not significantly different between the two classes of Ste12 binding sites (**Figure 8d**, left, p = 0.29). However, in *kar4*Δ cells, Ste12 occupancy was substantially lower for the H-T 4 motifs associated with Kar4-dependent transcription (mean occupancy, 66 RPM) than for those associated with Kar4-independent transcription (mean occupancy 227 RPM, p = 0.0077; **Figure 8d**, right, and see a zoomed-in version at **Supplementary Figure S18a**). Interestingly, although absolute Ste12 occupancy was different for the two types of promoters in *kar4*Δ cells, the change in Ste12 occupancy between wild-type and *kar4*Δ cells, as measured by the absolute change (**Supplementary Figure S18b**) or the fold-change (**Supplementary Figure S18c**), was not significantly different for the two types of promoters. These results suggest that, in either genotype, the absolute Ste12 occupancy is more relevant to transcription than fold-changes in occupancy between conditions. Further, it suggests that, in *kar4*Δ cells, Ste12 occupancy at the transcriptionally Kar4-dependent gene promoters falls below a minimum threshold needed to activate wild-type levels of transcription (see Discussion).

### *kar4*Δ-only genes are primarily Ste12 targets whose expression is not induced during the pheromone response in wild-type cells, and are enriched for T-T 3 PRE di-motifs

Cluster III (**Figure 2)** from our RNA-seq data consists of a set of genes that were upregulated only in the absence of Kar4 (designated *kar4Δ*-only genes). To identify the entire set of *kar4*Δ-only genes, we searched for genes up or down-regulated in response to pheromone at least 2-fold more in *kar4*Δ mutants than in wild-type. This revealed 84 upregulated and 13 downregulated genes (**Supplementary Figure S4c**). Among these genes was *STE12*, which in wild-type is modestly upregulated ∼1.6-fold in response to pheromone, but in *kar4*Δ is upregulated ∼3.5-fold, possibly implying negative feedback by Kar4 on *STE12* transcription in wild-type cells (also see next section for Ste12 binding within the *STE12* promoter). GO analysis of this group revealed only a weak enrichment for cell-wall-related proteins (9 genes, p-adj = 0.003). Of note, although we previously found that many GO terms were unique to the *kar4*Δ pheromone response (**Supplementary File S4**), these GO terms were not significantly enriched in the 84 *kar4*Δ-only genes. These 84 genes only include genes that are different from wild type by at least 2-fold (up- or down-regulated). In contrast, the GO analysis of the kar4Δ transcriptome included genes whose expression changed by at least 2 fold after pheromone treatment compared to the untreated condition, irrespective of how expression changed in wild type cells.

We next searched for a corresponding set of *kar4*Δ-only genes that are bound by Ste12 in *kar4Δ* cells but not wild-type, using the ChIP-exo data sets. We identified 57 sites with significantly increased Ste12 occupancy in *kar4*Δ, corresponding to 54 non-LTR genes (**Supplementary File S8**). There were no significant *kar4*Δ-only Ste12 binding sites prior to pheromone treatment. Top *kar4Δ*-only sites for Ste12 binding after pheromone treatment included the *CLA4* and *AXL1* promoters, and the *LRO1* gene body (**Figure 9a and Supplementary Figure S19**). We next asked whether these loci share a common DNA motif and identified a close match to a PRE di-motif with T-T orientation and 3 nt spacing (T-T 3 motif) in 40% of sites **(Figure 9b**). Analysis of all possible PRE di-motifs as above found the T-T 3 nt motif as the most enriched at 13% (**Figure 9c**). 27% of the best di-motifs contained a perfect PRE (TGAAACA).

**Figure 9.**
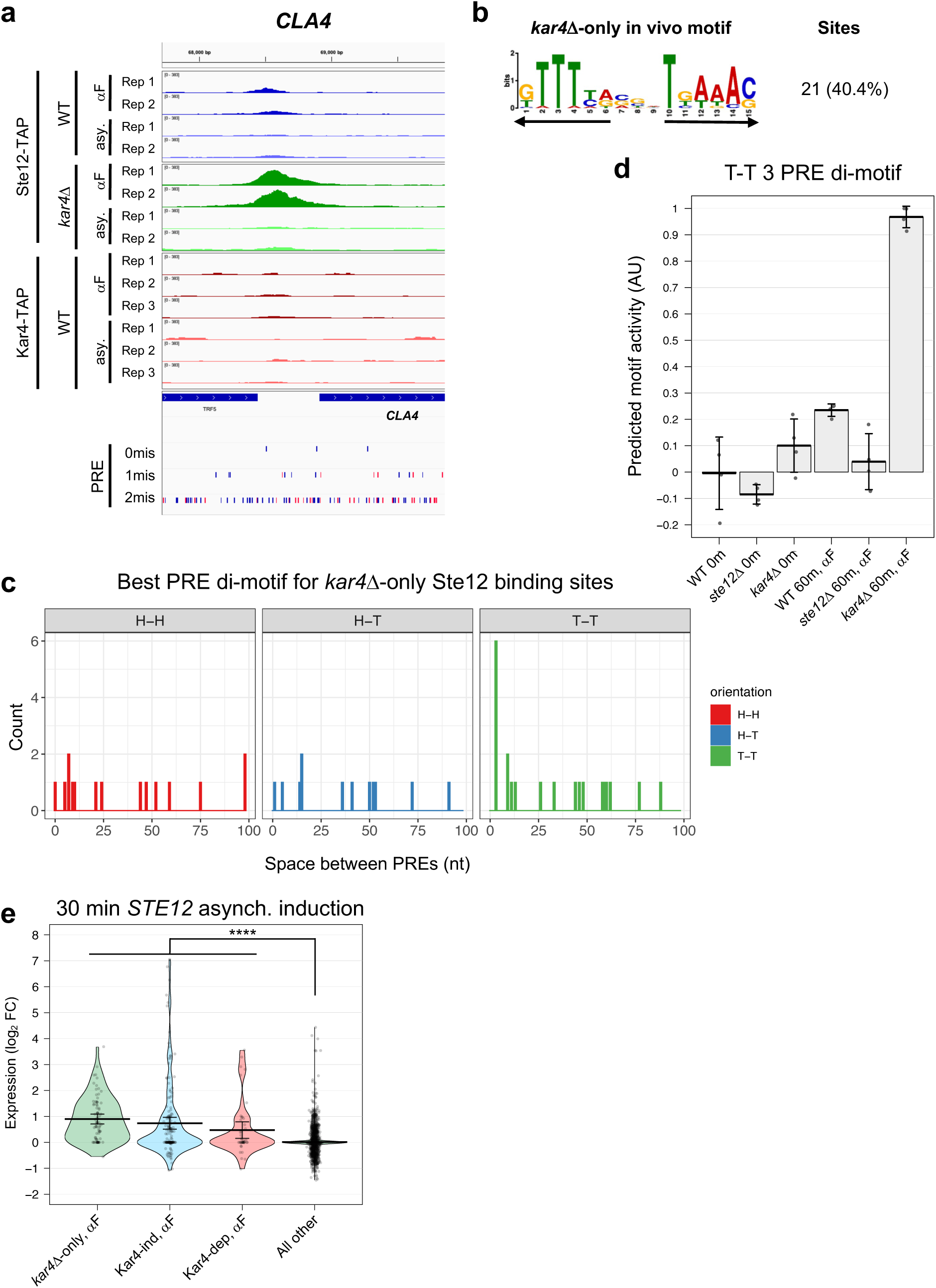
*kar4*Δ-only genes are associated with T-T 3 motif activity and are largely direct Ste12 targets. a) Example of *kar4*Δ-only Ste12 binding for the *CLA4* locus, as described in Figure 5a. b) de novo motif analysis (MEME) as in Figure 6b, but for sites enriched in *kar4*Δ cells relative to wild-type after pheromone treatment (namely *kar4Δ*-only sites). Arrows indicate PRE locations and orientation. c) Histogram, as in Figure 6c, indicating the best-match PRE di-motif for the *kar4*Δ-only Ste12-binding regions (n = 48). d) Predicted T-T 3 motif (from panel b) activity in indicated samples, as in Figure 7b. e) Gene expression log_2_ fold-change distribution for indicated gene sets 30 minutes after Ste12 over-expression, data from Hackett et al., 2020. Note that although *STE12* is a *kar4*Δ-only gene, it was excluded from the plot as its ZEV-induced expression (value of 6.1) would skew the distribution and is non-physiological. Error bars indicate 95% confidence-intervals. Asterisks indicate p-value < 1 x 10^−5^ for the 3 indicated gene sets against “All other” (Mann-Whitney U test). n-values for each group are given in **Table 1**.

**Table 1.** Gene set properties related to Kar4 based on RNA-seq data.

| Set | Size | Perfect PRE* |  | STE12-induced, 30 min** |  | STE12-induced, 90 min** |  |
| --- | --- | --- | --- | --- | --- | --- | --- |
|  |  | % | p-val | % | p-val | % | p-val |
| 1 Ste12-dep, Kar4-ind, α-F | 167 | 30.5 | 3.1E-11 | 22.2 | 4.3E-29 | 41.3 | 1.8E-41 |
| 2 Ste12-dep, Kar4-dep, α-F | 47 | 17.0 | 0.178 | 17.0 | 4.1E-06 | 55.3 | 4.1E-20 |
| 3 *kar4*Δ-only, α-F | 84 | 38.1 | 3.3E-10 | 41.7 | 1.9E-38 | 61.9 | 5.7E-43 |
| 4 All other | 6316 | 11.1 | 1 | 1.3 | 1 | 4.6 | 1 |

\* TGAAACA in 500 bp promoter sequence
\*\* Hackett et al., 2020

Despite identifying *kar4Δ*-only genes by both RNA-seq (84 genes) and ChIP-exo (sites associated with 54 genes), only 7 genes were found in common between the two datasets. Some of this could be due to the thresholding, as genes like *CLA4* and *AXL1* were significantly upregulated in *kar4*Δ cells by RNA-seq but missed the 2-fold cutoff. For the majority of *kar4*Δ-only genes, however, either Ste12 binds at a low level but below the limit of detection, or Ste12 does not bind at all and the genes are regulated by secondary transcription factors downstream of Ste12. These possibilities are addressed below. Nonetheless, to determine whether the *kar4Δ*-only genes in the two datasets reflect the same phenomenon, we performed an independent motif analysis on the entire promoter regions of the 84 *kar4*Δ-only genes as defined by RNA-seq. 38% of the *kar4*Δ-only gene promoters contained a perfect PRE, a value significantly higher than background rate of 11% for all gene promoters (**Table 1**). This value is similar to the percent of perfect PREs in genes induced by pheromone in a Ste12-dependent, Kar4-independent manner (30.5% of PREs associated with these gene promoters), and significantly higher than the percent of perfect PREs in genes induced by pheromone in a Ste12 and Kar4-dependent manner (17% of PREs). De novo motif discovery using pheromone-induced Kar4-independent genes as background identified a T-T 3 PRE di-motif as the most enriched among the *kar4Δ*-only genes from the RNA-seq dataset (21% promoters; **Supplementary Figure S20a)**, similar to our results using the *kar4*Δ-only Ste12 binding sites from ChIP-exo (**Figure 9b**). We also scanned the entire promoter regions of these *kar4*Δ-only genes for the best possible PRE di-motifs. Surprisingly, although we never observed the H-T 4 motif as the best match, the H-T 14 motif was the top match, at 8% (7/84), followed by the T-T 3 motif at 6% (5/84) (**Supplementary Figure S20b)**. These data suggest that, despite Ste12 binding not being detectable at the promoters of most of these genes by the ChIP-exo analysis, a large fraction of these promoters nevertheless contain functional Ste12 binding sites, raising the possibility that they are directly induced by Ste12 via non-H-T-4 motifs.

Although the T-T 3 motif was only modestly enriched in the *kar4Δ*-only gene promoters (or in the *kar4Δ*-only Ste12 binding sites by ChIP), we wondered whether transcriptional activity associated with this motif increased in *kar4Δ* cells. To this end, we predicted transcriptional activity associated with the motif from MEME (**Figure 9b**) using Transfactivity, as was done for the H-T 4 motif in **Figure 7c**. We found that while activity associated with the T-T 3 motif slightly increased during the wild-type pheromone response, this activity was dramatically increased in *kar4Δ* cells (**Figure 9d**). Therefore, these results demonstrate that transcriptional activity associated with the T-T 3 motif is increased in cells lacking Kar4. This is consistent with prior work suggesting that the T-T 3 motif is the preferred Ste12 DNA binding motif in vitro, where Kar4 was absent (Dorrity *et al*. 2018).

Finally, we reasoned that if the *kar4*Δ-only genes are induced directly by Ste12 in response to pheromone, we should be able to see induction under conditions of Ste12 overexpression during asynchronous culture growth, without pheromone, when Kar4 levels are low (Kurihara *et al*. 1996). To this end, we analyzed the transcriptional induction of the *kar4*Δ-only genes in a previously-published over-expression timecourse of *STE12* induction using cells dividing asynchronously (Hackett *et al*. 2020). Under these conditions, many Ste12 targets (including *KAR4*) were induced in the absence of pheromone. *STE12* expression was observed within 5 minutes of induction, and the very first wave of Ste12-dependent gene expression was observed after 15-30 minutes. After ∼30 minutes, secondary waves of induction occur by additional transcription factors other than Ste12 (Hackett *et al*. 2020). Consistent with *kar4*Δ-only genes being direct Ste12 targets, a large fraction (42%) of these genes was strongly induced by 30 minutes (**Figure 9e** and **Supplementary Figure S4c**), compared to 17% of Kar4-independent and 22% of Kar4-dependent pheromone-induced genes, and 1.3% of all other background genes (**Table 1**). By 90 minutes, at which point secondary transcription had occurred, strong transcriptional induction was observed for 41% of Kar4-independent genes, 55% of Kar4-dependent genes, and 62% of *kar4*Δ-only genes, compared to 4.6% of background genes. Further, when quantifying the induction of genes with either H-T 4 or T-T 3 motifs in their promoters by either Ste12 over-expression or pheromone (our RNA-seq dataset), Ste12 over-expression led to the induced expression of a larger number of genes containing T-T 3 motifs, much more than in our wild-type pheromone response and similar to the *kar4Δ* pheromone response (**Supplementary Figure S21**). Taken together, these data indicate that during pheromone treatment in *kar4*Δ cells half or more of the *kar4*Δ-only genes are directly induced by Ste12, often via binding to a T-T 3 motif. Our data further suggest that during pheromone exposure in wild-type cells, Kar4 prevents this induction of gene expression and instead directs Ste12 to promoters containing the H-T 4 motif (see Discussion).

### Interactions between H-T 4 and T-T 3 motifs during the pheromone response

After recognizing the importance of H-T 4 and T-T 3 motifs, we identified several promoters of key pheromone-regulated genes that contain both motifs. For example, the *STE12* promoter contains two distinct Ste12-binding peaks: an upstream peak which is largely Kar4-independent and pheromone-independent for binding, and a downstream peak which is strongly Kar4-dependent and pheromone-dependent for binding (**Figure 10**). Consistent with expectations based on our findings above, the upstream peak contains the best T-T 3 match in the genome (also present upstream of *GPA1*). In contrast, the Kar4-dependent downstream peak, which is occupied by Ste12 and Kar4 only in the presence of pheromone, contains an H-T 4 motif with a total of 3 mismatches (**Figure 10**, PRE tracks). As another example, the *FUS3* promoter contains a singular Ste12 binding peak which is partially but not entirely Kar4-dependent. This site contains both an H-T 4 and a T-T 3 motif which both share the same central perfect PRE sequence (**Supplementary Figure S22**). Thus, *FUS3* transcription, which is Kar4-independent, may arise from either motif in different scenarios. We have additionally identified strong T-T 3 motifs in at least the *FUS1*, *FUS2*, *PRM4*, *PRM6*, and *AGA1* promoters (typically in addition to an H-T 4 motif, and only considering Ste12-bound and transcriptionally pheromone-induced genes). Some of these genes have also been previously identified as having a T-T 3 motif in their promoter region (Dorrity *et al*. 2018). As many genes containing the T-T 3 motif in their promoters appear transcriptionally upregulated both during the *kar4*Δ pheromone response or when Ste12 is over-expressed in asynchronous cells, promoters containing both T-T 3 and H-T 4 motifs may be particularly sensitive to the Ste12:Kar4 ratio and may have a more complex regulatory pattern during the pheromone response. Overall, given that Kar4 affects the DNA sequence motifs to which Ste12 binds, it can be considered a Ste12 regulator.

**Figure 10.**
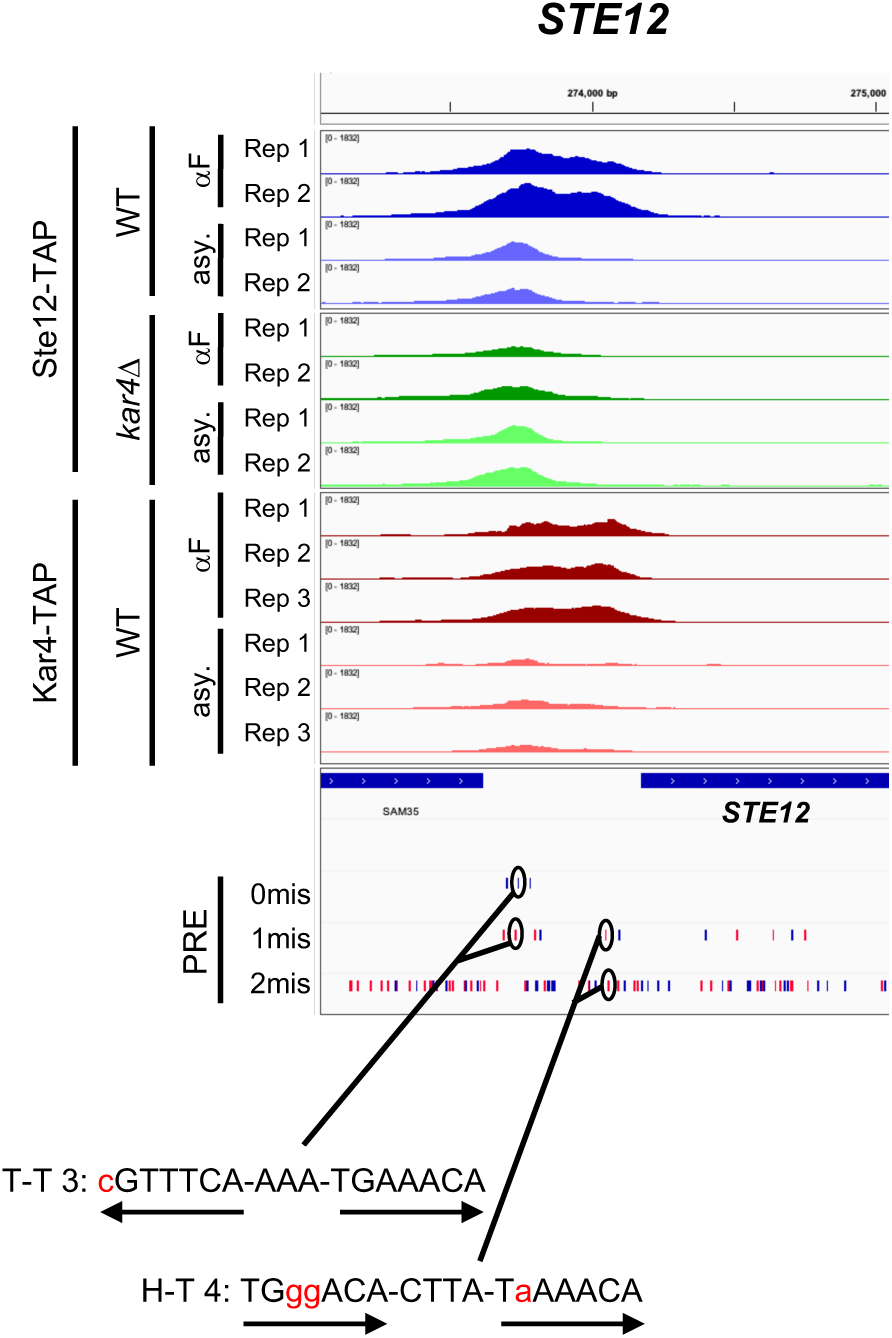
Example of a promoter containing both an H-T 4 and a T-T 3 motif. Shown is the *STE12* promoter. ChIP-exo data are shown as in **Supplementary Figure S7**. Positions of motifs are circled. Note that the H-T 4 motif is on the reverse strand, but shown as the reverse complement for clarity. Arrows denote location and orientation of PREs. See also **Supplementary Figure S22**.

## Discussion

Early work on the role of Kar4 during mating demonstrated that in the absence of Kar4, multiple genes are inadequately expressed, leading to nuclear congression defects (Kurihara *et al*. 1994; Kurihara *et al*. 1996; Lahav *et al*. 2007). Initially, Lahav et al. (2007) suggested that Kar4 might be a transcription factor that binds DNA directly in combination with Ste12, as is observed for Ste12-Tec1 binding. In contrast, subsequent work suggested that Kar4 is a Ste12 stabilization factor, prolonging Ste12 binding at suboptimal PREs (Aymoz *et al*. 2018). Additionally, although it was known that Ste12 can bind cooperatively to a pair of PREs, the location and configuration (i.e., orientation and spacing) of PREs bound by Ste12 at promoters in vivo remained unknown. Here we re-examined the role of Kar4 in Ste12 function by employing two approaches, RNA-seq on pre-synchronized cells, and ChIP-exo, with and without pheromone treatment. The effect of Kar4 was determined using cells in which *KAR4* was deleted. While the main focus of the study was the role of Kar4, our analyses provide new insights into the pheromone response, presented in the Appendix, as well as the discovery of a prevalent Ste12-binding PRE di-motif associated with pheromone-induced expression. As will be discussed below, the nature of this di-motif can also explain why genes that are pheromone-induced in a Kar4-dependent manner are expressed later than Kar4-independent genes.

Our mapping of Ste12-bound and Kar4-bound DNA by ChIP-exo revealed that Kar4 promotes the binding of Ste12 to the majority of Ste12 DNA-binding sites associated with pheromone-induced gene transcription. The majority of Ste12-bound pheromone-induced genes, most of which are Kar4-dependent for Ste12-binding, harbor a head-to-tail PRE di-motif with four nucleotide spacing (H-T 4 motif). In contrast, Kar4-independent Ste12-binding sites, most of which are not associated with pheromone-induced transcription, do not contain an H-T 4 motif. Moreover, we found that genes that are Kar4-dependent for transcription have particularly suboptimal H-T 4 motifs, often with neither PRE in the di-motif being a perfect match to the consensus sequence, while genes that are Kar4-independent for transcription have PREs that are closer to the consensus sequence. Lastly, in the absence of Kar4 (e.g., *kar4*Δ cells), Ste12 showed increased binding to non-H-T-4 motifs, particularly those with a T-T 3 configuration.

On the basis of the above evidence, we propose that rather than Kar4 being either a transcription factor or a general Ste12 stabilization factor, Kar4 functions as a Ste12 regulator, promoting Ste12 binding toward H-T 4 motifs, possibly by preventing other DNA-binding orientations (**Figure 11a**). By promoting binding to H-T 4 motifs, promoters with these motifs can contain highly suboptimal PREs and still be transcribed, but only in the presence of Kar4, explaining why some genes are Kar4-dependent for transcription (**Figure 11a, middle row**). In contrast, Kar4-independent promoters have near-optimal PREs, whether in the H-T 4 configuration or not, and thus can be strongly bound by Ste12 and can activate transcription even without Kar4 (**Figure 11a, top row**). Lastly, *kar4Δ*-only promoters contain PREs in non-H-T 4 configurations, often T-T 3, which in our model are better substrates for Ste12 binding in the absence than in the presence of Kar4. At these sites, while there may be a small amount of Ste12 binding during the pheromone response in wild-type cells (**Figure 11a, bottom left**), the amount of binding is not sufficient to activate transcription. In the absence of Kar4, the entire Ste12 pool, which can now bind DNA without Kar4, is available for binding non H-T 4 sites, such as T-T 3, leading to the increased expression of genes that fall under the *kar4Δ* only category (**Figure 11a, bottom right**).

**Figure 11.**
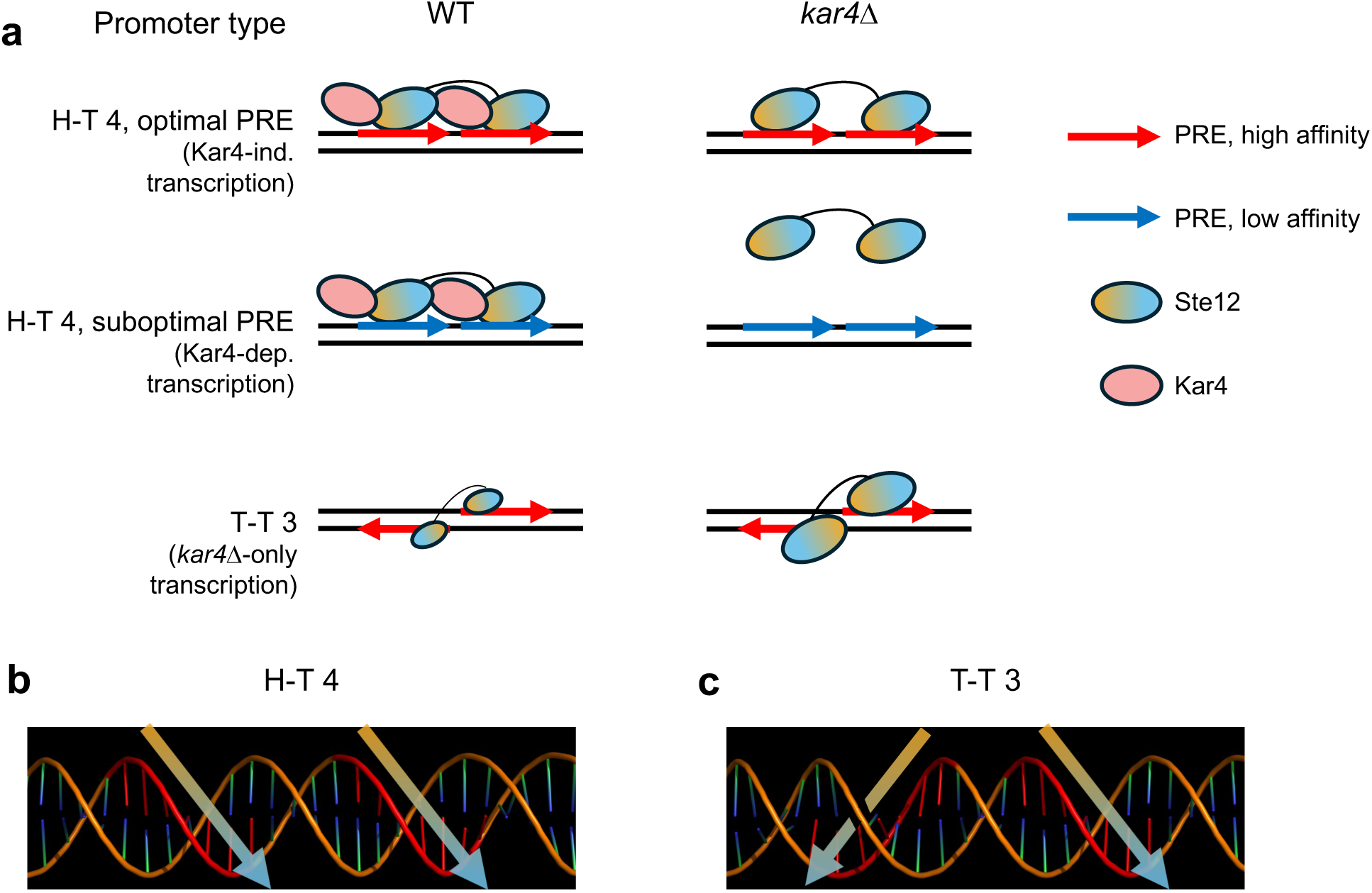
Model: Kar4 promotes Ste12’s binding to H-T 4 motifs. a) Model for the three classes of promoters of pheromone-induced genes related to Kar4, derived from our RNA-seq datasets. At Kar4-independent promoters (top row), the presence of high-affinity PREs (few mismatches) permits Ste12 binding with or without Kar4. At Kar4-dependent promoters (middle row), PREs are low affinity and require Kar4 for efficient Ste12 binding. In both classes, H-T 4 motifs are the most common, but other configurations may be functional as well (see Discussion). At *kar4*Δ-only promoters, only Ste12 complexes lacking Kar4 bind these sites efficiently, with T-T 3 being the most common although other configurations are likely included as well. Ste12 is drawn as smaller in wild-type to indicate a small pool of free Ste12 co-existing with the larger Kar4-bound Ste12 pool. Solid lines indicate an unknown dimer interface linking proteins together. Colors on Ste12 indicate direction, with orange at its N-terminus and blue at its C-terminus. b and c) Position of the PREs (red) on DNA (orange) in either an H-T 4 (b) or T-T 3 (c) configuration. Large arrows indicate direction of PRE (and a presumed direction Ste12 would be oriented).

Our model, whereby Kar4 is needed for binding to imperfect PRE di-motifs, is consistent with a prediction made by Su et al. (2010), who noted that many pheromone-induced gene promoters lack a perfect PRE and speculated that another factor may be required in conjunction with Ste12 to induce transcription in response to pheromone. Our model is also supported by in vivo mutational analysis of the *FIG1* promoter. This promoter region contains a suboptimal H-T 4 motif, TGAcACA-TACA-TGAAACc, as well as other PREs upstream and downstream. Optimizing the first PRE to TGA**A**ACA changed transcription from Kar4-dependent to Kar4-independent (Aymoz *et al*. 2018). Based on our model, this result is consistent with the idea Kar4 becomes dispensable for Ste12-DNA binding only when the sequence of the PRE is optimal. That Kar4 promotes Ste12 binding to H-T 4 motifs in vivo is also supported by the mutational analysis of the *KAR3* promoter, where Kar4-dependent pheromone-induced transcription was abolished when the already imperfect second PRE in the sequence TcAAACA-AAAT-caAAACA was made worse by mutating the second PRE to cagAACA (Lahav *et al*. 2007). Additionally, Lahav et al. (2007) demonstrated that while Ste12 (but not Kar4) can bind this sequence by itself in vitro, Kar4 significantly increases Ste12 binding affinity. Similarly, it was demonstrated that Ste12 overexpression in vivo can bypass the requirement for Kar4 for sufficient *KAR3* expression during mating, while Kar4 overexpression cannot bypass the requirment for Ste12 (Kurihara *et al*. 1996).

Both we and Lahav et al. (2007) observed that certain genes are induced in a pheromone-dependent manner only in the absence of Kar4 (referred to here as “*kar4Δ*-only” genes). This could have been consistent with the possibilty that Kar4 acts at these genes as a transcriptional repressor, but our model raises an alternative explanation. We propose that the expression of *kar4Δ*-only genes is absent in wild-type cells as a consequence of Ste12 being largely in a complex with Kar4, which is then redirected to H-T 4 motifs. In the absence of Kar4, Ste12 affinity in vivo likely reflects its affinity in vitro, favoring the T-T 3 motif (Dorrity *et al*. 2018), but also able to bind a wider variety of PRE orientations and possibly spacings, including H-T 4. Unlike sites linked to Kar4-independent transcription, however, the *kar4*Δ-only sites are poor substrates for the Ste12-Kar4 complex and are preferentially bound by the Ste12 complex lacking Kar4. This is supported by our finding that transcriptional activity associated with the T-T 3 motif increased dramatically during the pheromone response in *kar4*Δ cells compared to wild-type (**Figure 9d**). This is further consistent with experiments over-expressing Ste12 in asynchronous cells (where Kar4 levels are low and Ste12 complexes lacking Kar4 would dominate), where *kar4*Δ-only gene expression is readily observed, despite the cells being *KAR4*+ (**Figure 9e**). There may also be a contribution to kar4Δ-only gene expression from elevated levels of Ste12 itself, which is transcriptionally induced ∼2-fold more by pheromone in *kar4*Δ cells compared to wild-type.

We note, however, that the overlap between *kar4*Δ-only Ste12 binding and *kar4*Δ-only gene expression was particularly small, despite both groups being enriched for T-T 3 motifs. Similar examples of non-detectable Ste12 binding (i.e., false negatives) can also be observed at genes which are associated with strong Ste12 binding in wild-type cells and exhibit strongly pheromone-induced transcription in both wild-type and *kar4*Δ cells, but nevertheless exhibit no detectable Ste12 binding in *kar4*Δ cells. While some of this is likely due to the thresholds used to define each gene class, we note that such false negatives are common in ChIP studies, and are generally thought to be because 1) occupancy is too low to be detected (but still above the threshold needed to initiate transcription); 2) transcription arises from other, non-queried transcription factors; 3) there are differences in experimental design between the RNA-seq and ChIP-exo experiments; 4) a small fraction of cells give rise to most of the transcriptional response; or 5) the timepoint relevant for transcription factor binding was missed (Swift and coruzzi 2017; Mahendrawada *et al*. 2025). This latter possibility is interesting, as it would suggest that Kar4 affects the dynamics of the pheromone response, sustaining Ste12 activity for longer (here, at least 90 minutes after pheromone addition).

Based on our data, it appears that pheromone-mediated transcriptional induction broadly follows from a threshold of Ste12 occupancy, below which no transcription occurs, and above which other processes quickly become rate-limiting for determining mRNA levels. For example, in *kar4*Δ cells, Ste12 occupancy decreases for nearly all Ste12 binding sites associated with pheromone-induced transcription (**Figure 5c**), yet the amount of change in Ste12 occupancy is only weakly correlated with the corresponding change in transcription (**Figure 5d**). A less than 2-fold change in transcription was observed for the majority of genes, the transcriptionally Kar4-independent class, where a significant drop in Ste12 binding does not translate to a corresponding drop in gene expression. On an absolute level, however, Ste12 occupancy in *kar4*Δ cells is markedly lower for most transcriptionally Kar4-dependent genes compared to the Kar4-independent ones (**Figure 8d, right**), suggesting that there exists a minimum Ste12 occupancy threshold needed to activate wild-type levels of transcription in response to pheromone, and that Ste12 occupancy at most Kar4-independent genes is above that threshold even in *kar4*Δ cells. That there is still some overlap in the distributions of Ste12 occupancy in the absence of Kar4 (i.e., Ste12 occupancy is similar for some of the Kar4-independent and Kar4-dependent genes), however, suggests that the occupancy threshold is not constant but may vary between genes.

Our model also provides an explanation as to why, in the presence of pheromone, Kar4-dependent genes tend to be induced later than Kar4-independent genes. Lahav et al. (2007) proposed that Ste12 and Kar4 form a feed-forward loop whereby Ste12 induces the expression of Kar4 and both proteins together then induce the expression of late pheromone-response genes. It was unclear, however, why Kar4 was necessary for the expression of this late genes set. We found that H-T 4 motifs associated with Kar4-dependent genes tend to have more mismatches compared to the canonical PRE sequence, likely reducing the binding affinity of Ste12 to these suboptimal di-motifs in the absence of Kar4. Therefore, the association of suboptimal PREs with Kar4-dependent genes can provide a built-in timer mechanism that forces induced expression by Ste12 to be delayed until Kar4 levels are sufficiently high. A similar model was proposed by Aymoz et al. (2018), although they based it primarily on careful perturbation of two promoters, whereas here we have provided evidence at a genome-wide level.

We found many more GO categories associated with pheromone-induced transcription in *kar4*Δ cells compared to wild-type. Many of these categories were also altered in wild-type but the genes within them did not quite meet the 2-fold cutoff for significant changes in gene expression. This suggests that the global pheromone response was elevated in *kar4*Δ cells, implying that Kar4 imposes negative feedback on Ste12 during mating. This could be due to the increased transcription of *STE12*, which is upregulated by pheromone ∼1.6-fold in wild-type cells but ∼3.5-fold in *kar4*Δ cells. As we also observed that many genes are only expressed in *kar4*Δ cells, however, GO terms linked to these genes likely represent aberrant expression and processes not found during the normal mating response.

An open question in our model is how Kar4 structurally alters Ste12 DNA-binding specificity. We found that Kar4 and Ste12 are predicted by AlphaFold3 to dimerize via an interchain β-sheet. Such interchain β-sheet interactions are commonly found in protein dimers and other quaternary structures (Dou *et al*. 2004; Cheng *et al*. 2013). In our prediction, Kar4 appears to physically extend the region of Ste12 facing outward from the DNA binding domain (**Figure 2**). Notably, while the H-T 4 motif orients Ste12 proteins to face in the same direction on the same side (**Figure 11b**), the T-T 3 motif orients Ste12 such that the two proteins are on either side of the DNA with their tails facing each other (**Figure 11c**). Therefore, a simple model (as drawn in **Figure 11a**) is that when Kar4 is bound, Ste12 is physically occluded from dimerizing in a configuration that would fit the T-T orientation. This is largely speculative, however, and another model is that Ste12 complexes lacking Kar4 are highly flexible (to accommodate H-T, T-T, and H-H orientations), and Kar4 rigidifies the complex, restricting it to only the H-T orientation and possibly only the 4 bp spacing. However, we found that H-T 14 and H-T 15 motifs (which are one helical turn longer than H-T 4 and should be similarly functional to H-T 4 based on Pinheiro et al. (2025)) are also somewhat common at Kar4-dependent Ste12-binding sites (e.g., the *FUS2* and *MSS11* promoters contain an H-T 14 motif with no obvious good H-T 4 motif near the Ste12 binding site), suggesting that Kar4 may not restrict the distance between the PRE motifs. A third model, as has been observed for other transcription factor pairs, is that Ste12 does not dimerize at all, and cooperativity could be mediated by protein-DNA interactions which locally distort the DNA helix, enhancing nearby transcription factor binding without requiring direct protein-protein dimerization (Jolma *et al*. 2015). Given the importance of understanding how a single transcription factor can be modified to regulate different sets of genes under different conditions, discerning how Kar4 structurally affects Ste12’s DNA binding should be of particular interest in future studies. Overall, our study underscores the need to include Kar4 when examining the DNA binding activity of Ste12.

## Materials & Methods

### Strains

*S. cerevisiae* strains used for RNA-sequencing are *MAT***a** and derivatives of LPY1339 (*MAT***a** *ade1 bar1 cln1 cln2 cln3 his2 leu2:LEU2::GAL-CLN3 trp1 ura3*). Notably, all strains used had G1 cyclins knocked out with *CLN3* attached to a Gal-inducible promotor, allowing for pre-synchronization of cells in G1. To LPY1339 we tagged Ncl1 with GFP-NatMX to generate VM05, which served as the wild-type strain used for RNA-seq. Mutant strains were made by transformation of a PCR product from the Yeast Knockout Collect ion (YKO) into the VM05 background switching the gene of interest with KanMX (Giaever and nislow 2014). Strain names were ACY01, ACY02, and ACY13 for *kar4*Δ::KanMX, *ste12*Δ::KanMX, and *ime4*Δ::KanMX, respectively.

Strains used for ChIP-exo analysis were derived from the integrated TAP-tag library based on strain BY4741 (Ghaemmaghami *et al*. 2003) and obtained from Horizon Discovery (Cambridge, UK). TAP tags are linked to the His3MX6 marker. Strain BY4741 is *MAT***a** *his3*-Δ1 *leu2*-Δ0 *met15*-Δ0 *ura3*-Δ0. To construct the *kar4* deletion derivative (strain MY16682), *kar4*::KanMX was PCR-amplified from the yeast deletion collection using flanking primers (Kar4-up: TTTGTCGCTCTGTAGTCTTG and Kar4-down: GGTTTTGAGAAGAGTTTCCAG) and transformed into the Ste12-TAP strain.

### Pre-synchronization of cells and RNA extraction

For RNA-sequencing, cells were grown overnight at 30°C in YPGal (1% yeast extract, 2% peptone, and 2% galactose) to mid-log phase and then centrifuged and resuspended in YPD (1% yeast extract, 2% peptone, and 2% glucose) for 3 hours in order to arrest cells in G1 (**Figure 1a**). After the 3-hour synchronization, half of the culture (t0 – α-factor) was pelleted, the YPD was removed, and the pellet was flash frozen in liquid nitrogen for RNA extraction. To the second half of the culture, α-factor (10mM in 0.1 M sodium acetate pH 5.2, Zymo Research) was added at a 1:500 ratio (20 µM final concentration) for 60 minutes before the same freezing process was done. Total RNA was prepared using SDS, hot acid phenol, and chloroform, as previously described (Young and guydosh 2019).

### Library Preparation and RNA-sequencing

Libraries for RNA-sequencing were prepared by the NHLBI DNA Sequencing and Genomics Core at NIH (Bethesda, MD) using “Illumina Stranded Total RNA Prep, Ligation with Ribo-Zero Plus” kits (with 50-nt sequencing) from total RNA. During the reverse-transcription step of library preparation, removal of cytoplasmic and mitochondrial rRNA was done using the QIAseq FastSelect -rRNA Yeast Kit. Paired end sequencing (2 x 50 bp) was performed on an Novaseq 6000 machine at the NHLBI DNA Sequencing and Genomics Core at NIH (Bethesda, MD).

### Sequencing quality check and initial RNA-sequencing analysis

Fastq files from sequencing were trimmed of remaining sequencing adaptors and low-quality sequences using *bbtools*. The resulting fastq files were aligned to the sacCer3 reference genome using *STAR* (Dobin *et al*. 2013). Duplicate reads were removed and a table of read counts per gene per sample were generated using *featureCounts* (Liao *et al*. 2014). Read counts were used to run differential expression in R. Differential expression analysis was carried out using DESeq2 (v1.42.1) (Love *et al*. 2014). For the main dataset (WT, *kar4*Δ, and *ste12*Δ, compared at t0 and t60 + alpha-factor), we contrasted samples one treatment/genotype grouping at a time (i.e., for comparing *kar4*Δ t60 vs. WT t60, all t0 and all *ste12*Δ samples were excluded to minimize variance, and vice versa), and controlled for batch effect (i.e., the design formula was “∼ batch + treatment_genotype”). Distances between sample transcriptional profiles were estimated with the *dist* function of the R package *stats* (v4.3.1), while heatmaps summarizing transcriptional profile distances among samples were generated with the R package *pheatmap* (v1.0.12) **(Supplementary Figure S2b)**. PCA was performed with the *prcomp* function of the R package *stats* (**Supplementary Figure S2a)**. Log_2_ fold-change of genes was shrunk using *lfcShrink* from DESeq2 with the “ashr” method (Stephens 2017), and are generally used for all figures and analyses unrelated to quality-control plots.

Following differential expression using DEseq2, genes were classified as up or downregulated by using a log_2_ fold-change threshold of 1 or −1, respectively, and an adjusted p-value threshold of less than 0.05. By intersecting genes that were up or downregulated in the wild-type pheromone response (t60 + α-factor vs. t0 no α-factor) and the t60 response (t60 no α-factor vs. t0 no α-factor), we identified 4 pheromone-upregulated and 25 pheromone-downregulated genes which were potentially false positives due to the t60 treatment. These were removed from subsequent analyses related to gene set classifications and GO enrichment (i.e., not ChIP-exo).

### GO analysis

Overrepresentation analysis of Gene Ontology (GO) categories were identified using the function *enrichGO* from the package *clusterprofiler* (v4.10.1) in R (Yu *et al*. 2012). Only categories with an overlap of at least 3 genes were considered.

### Gene set clustering, classification, and analyses

Prior to clustering (**Figure 2)**, raw TPMs were calculated for all samples independently. A pseudocount of 0.5 was added to all TPM values, then log_2_ fold-changes were calculated for each sample against the average TPM of all WT control samples for each experiment. Throughout the paper, unless otherwise noted (quality control plots such as PCA and correlation matrices), non-protein-coding genes (except *RME2*), mitochondrial genes, and Ty-element-related features were removed from the analysis. Hierarchical clustering was performed using Cluster 3.0 (http://bonsai.hgc.jp/~mdehoon/software/cluster/) (De hoon *et al*. 2004) with a Pearson correlation (uncentered) similarity metric and average-linkage clustering. Heatmaps were generated using Java Treeview v1.2.0 (https://jtreeview.sourceforge.net/) (Saldanha 2004).

To define the gene sets used in **Supplementary Figure S4**, DESeq2 was used to define pheromone-regulated genes as those which were significantly (p-adj < 0.01) upregulated (or downregulated) with at least a shrunk fold-change of 2 in WT T60 cells vs. WT T0. These genes were then subsetted on those which were Ste12-dependent by keeping only genes which were significantly downregulated (if upregulated in wild-type) at least 2-fold in the *ste12*Δ T60 vs. WT T60 contrast (or vice versa for downregulated wild-type genes), leading to 215 pheromone-induced, Ste12-dependent genes. We then identified Kar4-dependent genes by keeping only those which were significantly downregulated (if upregulated in wild-type) at least 2-fold in the *kar4*Δ T60 vs. WT T60 contrast (or vice versa for downregulated wild-type genes), leading to 47 Kar4-dependent genes. Note that by this definition, *KAR4* is also Kar4-dependent (it is pheromone-induced in WT but not in *kar4*Δ), but we removed it from our analyses as based our data alone it is unclear how *KAR4* transcription would be affected in the absence of Kar4 function (it could be Kar4-dependent or Kar4-independent). The remaining 167 pheromone-induced genes were classified as Kar4-independent. *kar4*Δ-only genes were defined as those genes that increased (or decreased) expression significantly at least 2-fold in the *kar4*Δ T60 vs. WT T60 contrast, and were also significantly increased (or decreased) at least 2-fold in *kar4*Δ T60 vs. *kar4*Δ T0 samples (thus, their response to pheromone is at least 2-fold stronger than in WT, and they changed at least 2-fold in the *kar4*Δ pheromone response). Note that the final gene sets are not strictly mutually exclusive, as some of the *kar4*Δ-only genes are also classified as Kar4-independent pheromone upregulated. “All other” (**Figure 9e**) was defined as all remaining genes which were not classified in the other three gene sets.

### Motif identification

For *de novo* motif analysis of gene promoters, 500 bp promoter sequence (S288C reference genome, release R64-4-1) was used as input for XSTREME (https://meme-suite.org/meme/tools/xstreme) (Grant and bailey 2021), which simultaneously searches for motifs with STREME and MEME, with default settings and possible motif widths of 6-24 bp. Discovered motifs were automatically submitted to Tomtom (https://meme-suite.org/meme/tools/tomtom) (Gupta *et al*. 2007) to search for known motifs with sequence similarity using default settings and the YEASTRACT database as reference.

### Transcription factor and motif activity inference

Transcription factor activities were inferred using Transfactivity from the REDUCE suite (http://reducesuite.bussemakerlab.org/) (Bussemaker *et al*. 2001; Roven and bussemaker 2003; Lee and bussemaker 2010). Inputs were PSAMs (position-specific affinity matrix) derived from YeTFaSCo expert-curated motifs (http://yetfasco.ccbr.utoronto.ca/1.02/Downloads/Expert_PFMs.zip) (De boer and hughes 2012), and 600 bp promoter sequence (excluding any upstream ORF sequence) for all genes. All values are obtained from a multivariate analysis including all the expert-curated motifs. For **Figure 7a**, we used as input shrunk log_2_ fold-changes obtained from DESeq2 for all genes. The direction of the fitted score value (negative or positive) was then assigned to the p-values. For all remaining figures using Transfactivity, the normalized gene expression matrix for all samples (shown in **Figure 2)** was used as input. Fitted score values for each motif were then rescaled such that the largest absolute value is 1 prior to plotting.

For di-motif analyses, the motifs were converted from a position-specific probability matrix (PSPM) to PSAM format using Convert2PSAM (REDUCE suite). Coefficient values were derived from a multivariate model including the standard expert-curated motifs from YeTFaSCo (including the Ste12 mono-motif, see previous paragraph), as well as the H-T 4 and T-T 3 di-motifs. Tests on reduced models indicated the three motifs (PRE alone, H-T 4, and T-T 3) were largely orthogonal and all significantly contributing.

### Ste12-Kar4 structure prediction

One copy each of Kar4-short (residues 31-335), the Ste12 DNA-binding domain (residues 21-220), and DNA consisting of a H-T 4 motif (AGATGA**TGAAACA**AACA**TGAAACA**TCTGC) and its reverse complement strand were used as input for modeling by AlphaFold3 (https://alphafoldserver.com/) (Abramson *et al*. 2024) using default parameters. The top predicted model was visualized in ChimeraX. Note that multiple DNA sequences were modeled and the predicted DNA-binding interface was not specific to the PRE.

### ChIP-exo sample preparation and data generation

Cells were grown to mid log-phase at 30 °C in YPD pH 3.5 (buffered with 85 mM succinic acid) and split into two portions. For the pheromone-treated samples, alpha factor (ABI Scientific; Sterling, VA) dissolved in methanol was added to the culture to 10 µg/ml (6 µM) final concentration. An equal volume of methanol (to 1% v/v final concentration) was added to control cells. Cells were then grown for an additional 90 minutes, at which point >95% had formed shmoos, and formaldehyde was added to a final concentration of 1% and incubated for 15 minutes at 30 °C with shaking. Crosslinking was stopped by the addition of 2.5 M glycine to a final concentration of 125 mM. Approximately 50 OD of crosslinked cells were harvested by centrifugation, washed twice with ST buffer (10 mM Tris-Cl pH 7.5, 100 mM NaCl) containing a protease inhibiter cocktail at 0.2% (cOmplete mini, EDTA-free, Roche), and cell pellets were flash frozen in liquid nitrogen. Samples were shipped to the Cornell BRC Epigenomics Facility for ChIP-exo processing and sequencing. Following bead beating and sonication, chromatin from TAP-tagged yeast strains was incubated overnight on IgG-conjugated magnetic Sepharose beads. Libraries were prepared from immunoprecipitated DNA as described in the ChIP-exo 5.1 protocol (Rossi *et al*. 2018) pooled at equimolar concentration, and size selected by agarose gel excision (200-500bp). Sequencing was performed in-house on an Element Biosciences AVITI sequencer (2 x 75bp high output kit) to an average of 10M paired-end reads per sample, with data management, quality control, and processing through the PEGR automated workflow (Shao *et al*. 2022).

### ChIP-exo data processing and peak calling

Reads were aligned using bwa-mem (Li 2013) against the sacCer3 genome using default options. Aligned and sorted BAM files then had duplicates and non-uniquely mapped reads removed using Picard Tools MarkDuplicates (https://github.com/broadinstitute/picard) using default options (except VALIDATION_STRINGENCY=’LENIENT’) and samtools (v1.19.2 and htslib v1.19.1) (Bonfield *et al*. 2021; Danecek *et al*. 2021). BAM files were then further cleaned using samtools view -f 2 (read mapped in proper pair) and -q 30 (minimum mapping quality 30) options. Note that the mapping quality restriction additionally removes any reads mapping to repetitive or duplicated regions, such as rDNA and Ty elements.

Peaks were identified using ChExMix (v0.52), a peak-calling software optimized specifically for ChIP-exo datasets (Yamada *et al*. 2020). To identify all peaks for a given condition (**Supplementary File S7**), all replicates were specified as independent signal experiments with no control, using the –design option. To generate specific contrasts (e.g., Kar4-dependent sites, **Supplementary File S8**), the –design option was used to specify the relevant signal experiments (i.e., Ste12-TAP WT with pheromone) against their corresponding control experiments (i.e., Ste12-TAP *kar4*Δ with pheromone). In all cases, we specified the following non-default options: –back using the yeast.back file provided by ChExMix (http://lugh.bmb.psu.edu/software/chexmix/backgrounds/yeast.back), –exclude against a file containing the coordinates for all yeast telomeres, the rDNA locus, and the mitochondrial genome, –mememaxw 24 for wider motif width discovery, and –round 5 for more iterations.

### ChIP-exo downstream data analysis

Analyses downstream of ChExMix were performed using custom R scripts. First, significant peaks identified by ChExMix were associated with nearby genes and LTRs using the sacCer3 R64-4-1 (20230830) GFF annotation file (http://sgd-archive.yeastgenome.org/sequence/S288C_reference/genome_releases/S288C_reference_genome_R64-4-1_20230830.tgz). Additionally, peaks that were either closest to an LTR start site (compared to gene start sites) or inside of an LTR feature were flagged as LTR-associated for downstream filtering. To perform gene-based analyses, when multiple peaks were associated with a single gene we used the statistics (occupancy, fold-change, Q-value, etc.) from the peak with the lowest Q value (most significant). Peaks could be associated with multiple features (genes or LTRs) if there were multiple feature start sites within 500 bp of the peak in either direction (bidirectional promoters, e.g. *ERG24*/*PRM1*). If a gene had any peaks associated with both it and an LTR, the gene was flagged as LTR-associated for downstream filtering. All figures use non-LTR-associated gene statistics, unless otherwise stated. For **Figure 5c**, in order to fairly compare WT and *kar4*Δ occupancy statistics at the same peaks, we started by using ChExMix to contrast Ste12-TAP WT vs. *kar4*Δ samples (this gives Kar4-dependent fold-changes used in **Figure 5d**), then, instead of only considering significantly different peaks, we selected statistics for all genes that exhibited significant Ste12 binding from a separate analysis, even if they were not significantly different between WT and *kar4*Δ samples. This resulted in a very slight difference (394 instead of 399) in the number of Ste12-bound genes analyzed due to very slight shifts in peak locations, causing gene associations to differ in a few cases.

### ChIP-exo data visualization

To visually compare ChIP-exo data, final cleaned BAM files were depth-normalized using bamCoverage (v3.5.5, (Ramirez *et al*. 2016)) ‘--binSize 1 --normalizeUsing CPM - e’. The resulting bigWig files were visualized using IGV (v2.11.9, (Robinson *et al*. 2011; Thorvaldsdottir *et al*. 2013)).

### Motif analysis from ChIP-exo data

For *de novo* motif analysis, 120 bp genomic DNA surrounding non-LTR sites (60 bp up and down from each site) was extracted and submitted to XSTREME (which runs both STREME and MEME; https://meme-suite.org/meme/tools/xstreme) (Grant and bailey 2021) using default options except motif width was set to 6-24 nt. Shuffled input sequence was used for background. If sites were within 60 bp of each other their sequences were merged into one long sequence (60 bp up from the first site to 60 bp down from the final site, with all intervening sequence included) to prevent false positive enrichments from duplicated sequence input.

For analyses related to **Figure 6d** and **Supplementary Figure S15**, Kar4-independent sites were defined as sites significantly bound by Ste12 during pheromone treatment in wild-type (sites underlying **Figure 4c**, grey circle), and not within 100 bp of any significantly Kar4-dependent Ste12 binding sites. This yielded 291 Kar4-independent non-LTR sites, but many of these sites had weak Ste12 binding or were likely false positive genomic regions. For motif analysis, we opted to exclude these sites in favor of the strongest Kar4-independent Ste12 binding sites by restricting the list to high-confidence sites (log_2_Q < −30), leaving 111 Kar4-independent Ste12-binding sites before merging nearby sites. For comparing Ste12 occupancy associated with Kar4-independent and Kar4-dependent Ste12-binding peaks in **Figure S15b**, first we reduced each mapped read pair down to a “tag” position using the final cleaned BAM file using bamCoverage (v3.5.5) --binSize 1 --scaleFactor 1.0 --minMappingQuality ‘1’ --Offset 1 with --samFlagInclude 64 --samFlagExclude 16 for the top strand and --samFlagInclude 80 for the bottom strand. Then, tags were manually counted over Ste12-binding regions using multiBigWigSummary (v3.5.5 (Ramirez *et al*. 2016)) and multiplying the output by the region length. Counts were then summed across strands and replicates and normalized to RPM using read-depth scaling factors previously calculated by ChExMix.

For PRE di-motif discovery, sites were generated as above except using 240 bp (120 bp up and 120 bp down) genomic DNA and submitted to FIMO (https://meme-suite.org/meme/tools/fimo) (Grant *et al*. 2011). Sequences were scanned using all PRE (TGAAACA) di-motifs with spacing (N) from 0-100 nt and H-H, H-T, and T-T orientations. Note that T-H was not input separately because it is identical to H-T motifs on the bottom strand, and both strands were searched by default. Allowed matching p-value was set to 0.01 for match calling. Each site was then assigned a best match by using the lowest q-value from all motif matches.

### Set comparisons

Venn diagrams comparing gene sets were generated using the eulerr package in R.

### DNA motif models

To visualize the H-T 4 and T-T 3 motifs on DNA (**Figure 11b and c**), a double-stranded DNA helix in the B conformation was generated using PyMOL (3.1.3) with the command ‘fnab GAGTAAAAGAATTTTGTGTTTCAGGGTGAAACAGATCTGAAACACAAGAGCTTATGCATT.’

### Statistical analysis

All statistical analyses were performed in R (v 4.3.1). Specific tests and corrections used are stated in main text or figure legends. Generally, distributions were compared using a pairwise Wilcoxon rank sum test (also known as a Mann-Whitney U test) and p-values were corrected for multiple comparisons using the Benjamini-Hochberg method. If data were normally distributed then a t-test was used. For gene set comparisons (e.g., in **Table 1**), a hypergeometric test was used.

## Data availability

Raw RNA-seq read files and gene-count matrix can be found at GEO accession ID GSE306669. Raw ChIP-exo read files and final depth-normalized read alignments for use with IGV can be found at GEO accession ID GSE306667. Scripts to reproduce specific figures or analyses will be made available upon request.

## Acknowledgments

We thank the Cornell University BRC Epigenomics Facility (<u>RRID:SCR_021287</u>) for performing ChIP-exo data collection and initial data processing. We thank Lorraine Pillus (UCSD) for kindly providing the *cln* mutant strain (LPY1339) used to generate strains for G1 arrests in RNA-seq experiments. We also thank David Young (NIH, NIDDK) and Ellen Morgan (NIH, NIDDK) for assistance and advice with the RNA-seq experiments. We thank Audrey Gasch (University of Wisconsin-Madison) and Jeremy Thorner (UC Berkeley) for advice and suggestions, and Patrick Klees (NIH, NIDDK) for comments on the manuscript.

This research was supported in part by the Intramural Research Program of the National Institute of Diabetes and Digestive and Kidney Diseases (NIDDK) within the National Institutes of Health (NIH). The contributions of the NIH author(s) are considered Works of the United States Government. The findings and conclusions presented in this paper are those of the author(s) and do not necessarily reflect the views of the NIH or the U.S. Department of Health and Human Services.

## Supplementary File list

File S1: Raw log_2_ fold-changes for RNA-seq samples for wild-type, kar4Δ, ste12Δ, and ime4Δ in the presence and absence of pheromone

File S2: Differential expression analyses for RNA-seq derived from datasets in File S1

File S3: Main gene sets used in this study

File S4: GO term analysis related to our RNA-seq dataset

File S5: AlphaFold3 Ste12-Kar4 model output (.cif file)

File S6: ChIP-exo sample metadata and summary statistics

File S7: ChIP-exo peak annotations for each genotype and treatment combination

File S8: ChIP-exo differential contrast results

File S9: Best PRE di-motif annotations for various subsets of Ste12-binding regions

File S10: Curated motif annotations related to **Figure S17a**

