## Supplemental Figures and legends for "Kar4 acts as a Ste12 regulator in *Saccharomyces cerevisiae*, promoting Ste12 binding to a specific DNA motif genome-wide"

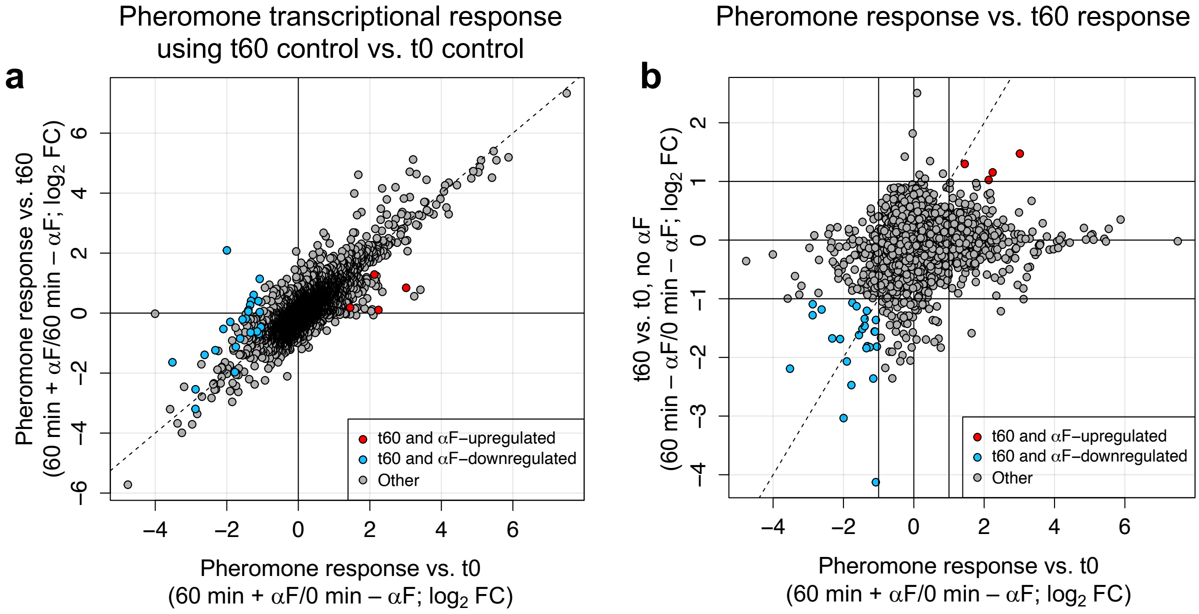

Pheromone transcriptional response
using t60 control vs. t0 control
a
Pheromone response vs. t60
(60 min + αF/60 min – αF; log2 FC)
6
4
2
0
−2
−4
−6
−4
−2
0
2
4
6
t60 and αF–upregulated
t60 and αF–downregulated
Other
Pheromone response vs. t0
(60 min + αF/0 min – αF; log2 FC)
Pheromone response vs. t60 response
b
t60 vs. t0, no αF
(60 min – αF/0 min – αF; log2 FC)
2
1
0
−1
−2
−3
−4
−4
−2
0
2
4
6
t60 and αF–upregulated
t60 and αF–downregulated
Other
Pheromone response vs. t0
(60 min + αF/0 min – αF; log2 FC)


**Figure S1**

**Supplementary Figure S1**. a) Gene expression changes in the t60 + $\alpha$-factor condition relative to either t0 no $\alpha$-factor (x-axis) or t60 no $\alpha$-factor (y-axis). Note that each t60 + $\alpha$-factor group is from its own experiment to remove batch effects. Genes that were removed as being potentially t60 artifacts (i.e., genes which increased expression due to extended time in G1 arrest rather than pheromone treatment) are shown in red and blue. Dashed line indicates 1:1 line. b) As in panel a, but showing the t60 no $\alpha$-factor vs. t0 no $\alpha$-factor comparison, which was used to define the excluded genes. Dashed line indicates 1:1 line, and solid lines indicate 2-fold change. Note that as these were performed in separate experiments, there is also a small batch effect between the samples being compared on the y-axis. Genes which were up or downregulated $\geq$ 2-fold on both axes were flagged (blue and red points) and excluded in future analyses.

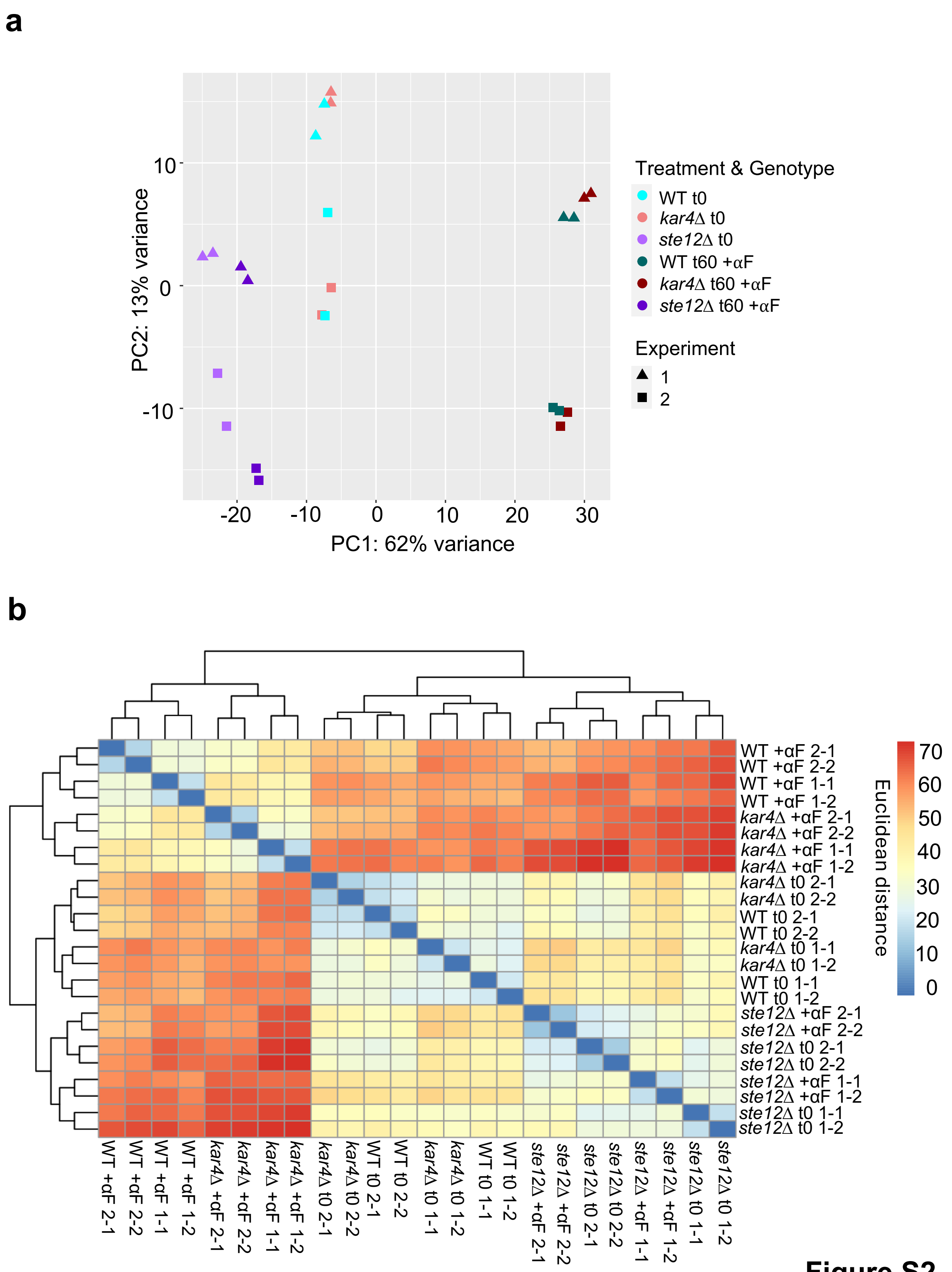
a
PC2: 13% variance
PC1: 62% variance
Treatment & Genotype
WT t0
kar4Δ t0
ste12Δ t0
WT t60 +αF
kar4Δ t60 +αF
ste12Δ t60 +αF
Experiment
1
2
b
Euclidean distance
WT +αF 2-1
WT +αF 2-2
WT +αF 1-1
WT +αF 1-2
kar4Δ +αF 2-1
kar4Δ +αF 2-2
kar4Δ +αF 1-1
kar4Δ +αF 1-2
kar4Δ t0 2-1
kar4Δ t0 2-2
WT t0 2-1
WT t0 2-2
kar4Δ t0 1-1
kar4Δ t0 1-2
WT t0 1-1
WT t0 1-2
ste12Δ +αF 2-1
ste12Δ +αF 2-2
ste12Δ t0 2-1
ste12Δ t0 2-2
ste12Δ +αF 1-1
ste12Δ +αF 1-2
ste12Δ t0 1-1
ste12Δ t0 1-2

**Figure S2**

**Supplementary Figure S2**. a) Principal component analysis (PCA) plot of all RNA-seq gene expression data for the indicated genotypes and treatments. Experiment indicates different batches of data collection. Variance explained by each principal component is shown on each axis. b) Clustered similarity matrix of all RNA-seq gene expression data. Numbers next to the genotype and condition indicate batch (1- or 2-) and replicate (-1 or -2).

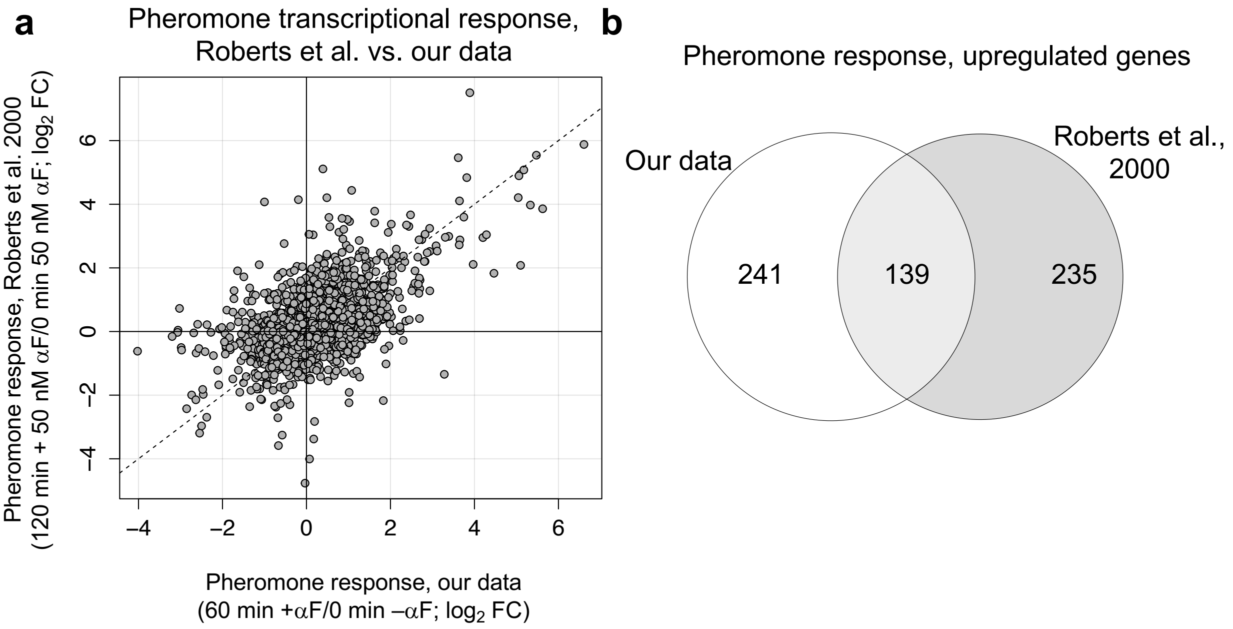
a
Pheromone transcriptional response,
Roberts et al. vs. our data
Pheromone response, Roberts et al. 2000
(120 min + 50 nM αF/0 min 50 nM αF; log2 FC)
Pheromone response, our data
(60 min +αF/0 min –αF; log2 FC)
–4
–2
0
2
4
6
b
Pheromone response, upregulated genes
Our data
Roberts et al.,
2000
241
139
235

**Figure S3**

**Supplementary Figure S3**. Scatterplot of the fold-change in gene expression for all genes in response to pheromone in wild-type cells in our data vs. Roberts et al. (2000). Fold-changes in our data were calculated using DESeq2. b) Venn diagram of the overlap between genes significantly upregulated at least 2-fold in our data vs. Roberts et al. (2000). Genes were required to have adjusted p-value < 0.05 for our data and unadjusted p-value < 0.01 for Roberts.

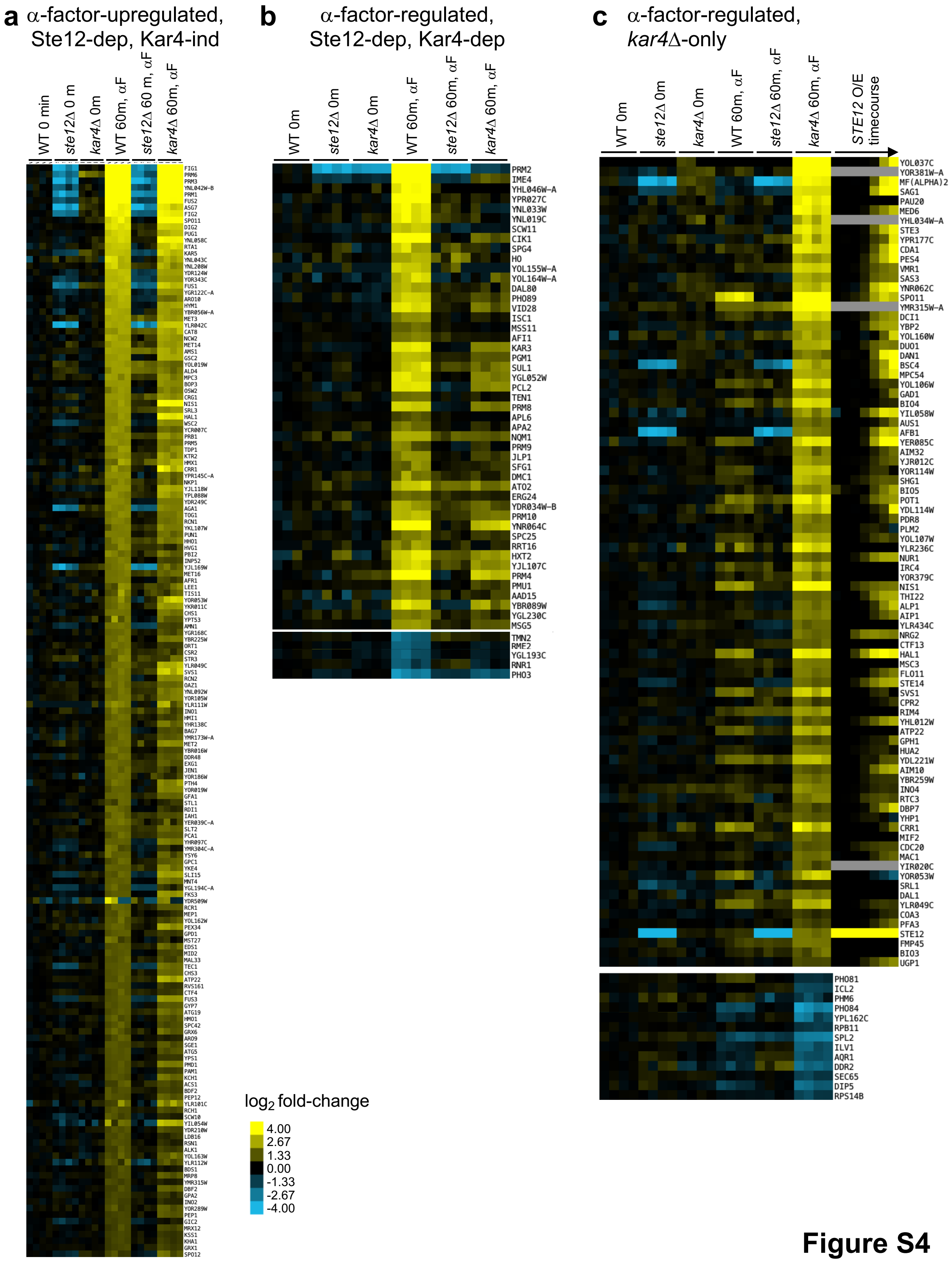

a α-factor-upregulated, Ste12-dep, Kar4-ind
WT 0 min
ste12Δ 0 m
kar4Δ 0m
WT 60m, αF
ste12Δ 60 m, αF
kar4Δ 60m, αF
b α-factor-regulated, Ste12-dep, Kar4-dep
WT 0m
ste12Δ 0m
kar4Δ 0m
WT 60m, αF
ste12Δ 60m, αF
kar4Δ 60m, αF
PRM2
IME4
YHL046W-A
YPR027C
YNL033W
YNL019C
SCW11
CIK1
SPG4
HO
YOL155W-A
YOL164W-A
DAL80
PHO89
VID28
ISC1
MSS11
AFI1
KAR3
PGM1
SUL1
YGL052W
PCL2
TEN1
PRM8
APL6
APA2
NQM1
PRM9
JLP1
SFG1
DMC1
ATO2
ERG24
YDR034W-B
PRM10
YNR064C
SPC25
RRT16
HXT2
YJL107C
PRM4
PMU1
AAD15
YBR089W
YGL230C
MSG5
TMN2
RMF2
YGL193C
RNR1
PHO3
c α-factor-regulated, kar4Δ-only
WT 0m
ste12Δ 0m
kar4Δ 0m
WT 60m, αF
ste12Δ 60m, αF
kar4Δ 60m, αF
STE12 O/E timecourse
YOL037C
YOR381W-A
MF(ALPHA)2
SAG1
PAU20
MED6
YHL034W-A
STE3
YPR177C
CDA1
PES4
VMR1
SAS3
YNR062C
SPO11
YMR315W-A
DCI1
YBP2
YOL160W
DUO1
DAN1
BSC4
MPC54
YOL106W
GAD1
BIO4
YIL058W
AUS1
AFB1
YER085C
AIM32
YJR012C
YOR114W
SHG1
BIO5
POT1
YDL114W
PDR8
PLM2
YOL107W
YLR236C
NUR1
IRC4
YOR379C
NIS1
THI22
ALP1
AIP1
YLR434C
NRG2
CTF13
HAL1
MSC3
FLO11
STE14
SVS1
CPR2
RIM4
YHL012W
ATP22
GPH1
HUA2
YDL221W
AIM10
YBR259W
INO4
RTC3
DBP7
YHP1
CRR1
MIF2
CDC20
MAC1
YIR020C
YOR053W
SRL1
DAL1
YLR049C
COA3
PFA3
STE12
FMP45
BIO3
UGP1
PHO81
ICL2
PHM6
PHO84
YPL162C
RPB11
SPL2
ILV1
AQR1
DDR2
SEC65
DIP5
RPS14B
log2 fold-change
4.00
2.67
1.33
0.00
-1.33
-2.67
-4.00


**Figure S4**

**Supplementary Figure S4.** Gene sets from RNA-seq experiments with wild-type, *ste12*Δ and *kar4*Δ cells, in the absence (0 min) or presence (60 min, αF) of pheromone, exhibiting a) Kar4-independent (n = 167 up), b) Kar4-dependent (n = 47 up, n = 5 down), or c) *kar4*Δ-only (n = 84 up, n = 13 down) response to pheromone (see methods for how each group was defined). Values are raw $\log_2$ fold-changes, as in **Figure 2**. In panel c, the gene expression heatmap for *kar4*Δ-only upregulated genes is shown with a *STE12* over-expression timecourse, based on data from Hackett et al., 2020. Timecourse timepoints are (from left to right) 5, 10, 15, 20, 30, 45, and 90 minutes. This dataset is discussed later in the study.

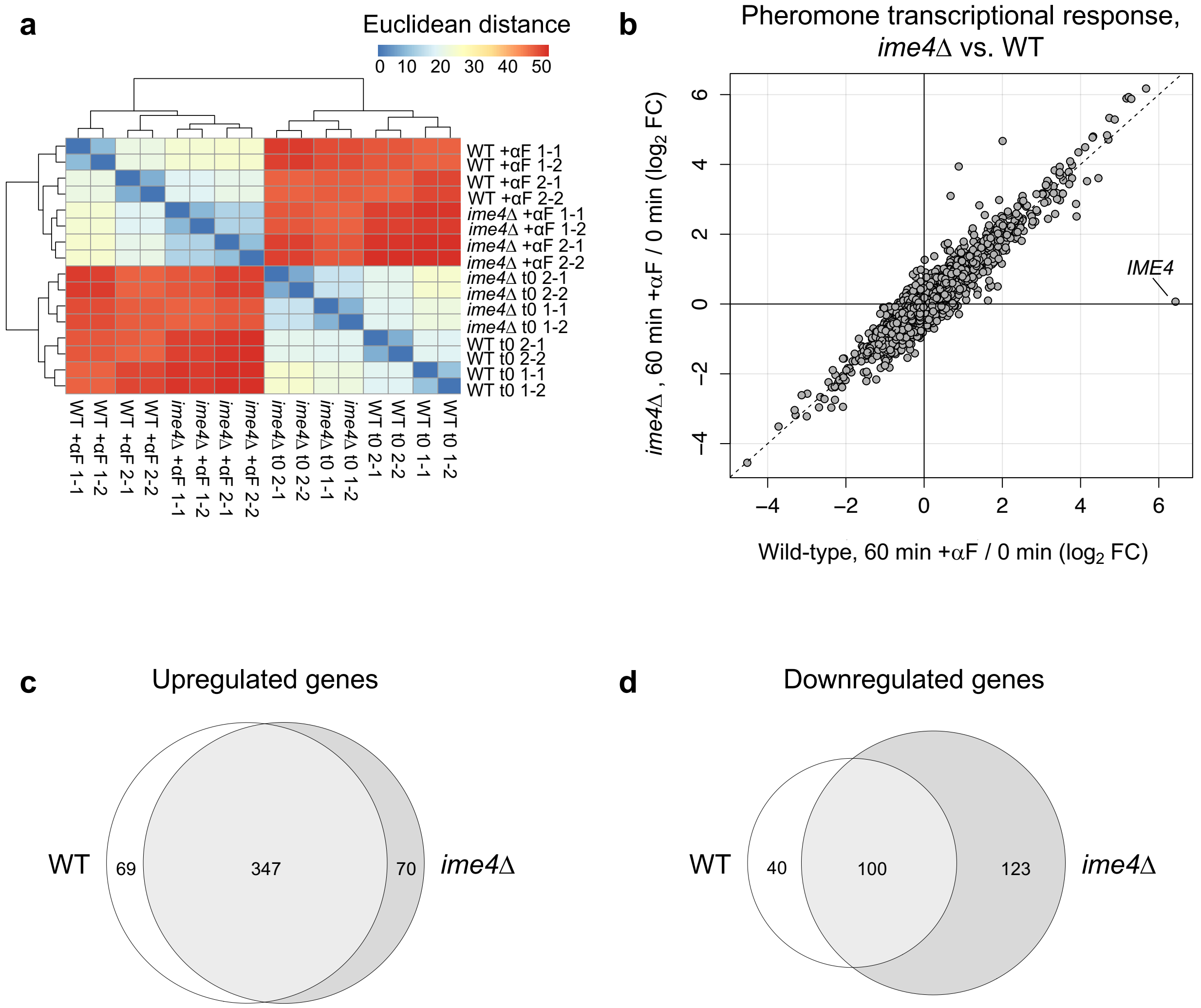
a
Euclidean distance
0 10 20 30 40 50
WT +αF 1-1
WT +αF 1-2
WT +αF 2-1
WT +αF 2-2
ime4Δ +αF 1-1
ime4Δ +αF 1-2
ime4Δ +αF 2-1
ime4Δ +αF 2-2
ime4Δ t0 2-1
ime4Δ t0 2-2
ime4Δ t0 1-1
ime4Δ t0 1-2
WT t0 2-1
WT t0 2-2
WT t0 1-1
WT t0 1-2
b
Pheromone transcriptional response,
ime4Δ vs. WT
ime4Δ , 60 min +αF / 0 min (log2 FC)
Wild-type, 60 min +αF / 0 min (log2 FC)
IME4
−4
−2
0
2
4
6
c
Upregulated genes
WT
69
347
70
ime4Δ
d
Downregulated genes
WT
40
100
123
ime4Δ

**Figure S5**

**Supplementary Figure S5.** Analysis of pheromone response in *ime4*Δ cells. a) Similarity matrix (as in **Supplementary Figure S2b)** comparing samples related to *ime4*Δ. b) Scatterplot comparing $\log_2$ fold-changes after pheromone treatment for wild-type and *ime4*Δ cells, as **Figure 1c**. c and d) Overlap between significantly up- and down-regulated genes, as in **Figure 1b**.

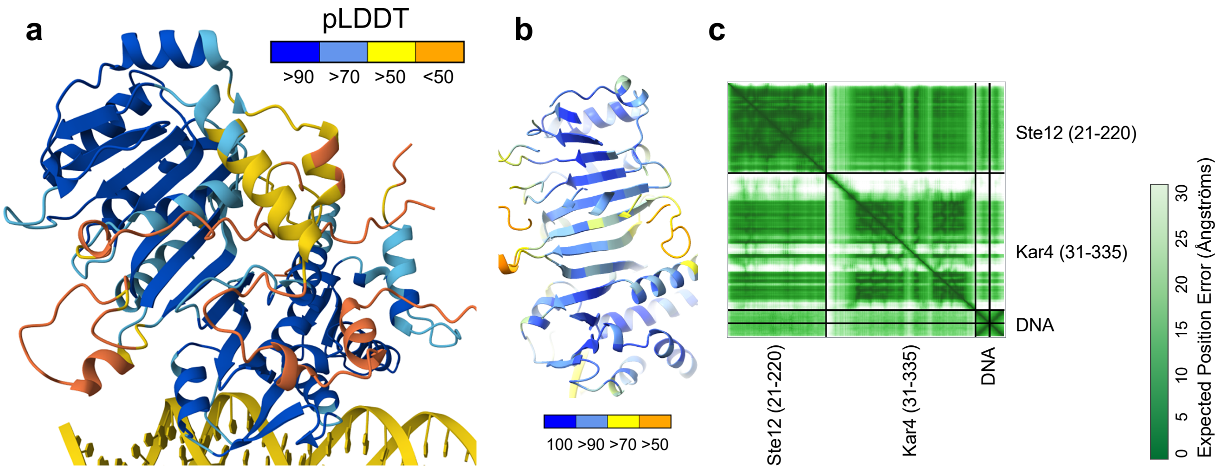
a
pLDDT
>90 >70 >50 <50
b
100 >90 >70 >50
c
Ste12 (21-220)
Kar4 (31-335)
DNA
Ste12 (21-220)
Kar4 (31-335)
DNA
0 5 10 15 20 25 30
Expected Position Error (Ångströms)

**Figure S6**

**Supplementary Figure S6.** a) Predicted AlphaFold3 Ste12-Kar4-DNA structure (as in **Figure 3**), colored by confidence level (pLDDT). b) Zoom-in of predicted extended β-sheet structure. c) Predicted Aligned Error (PAE) matrix for the model, indicating the accuracy of inter-residue positional prediction.

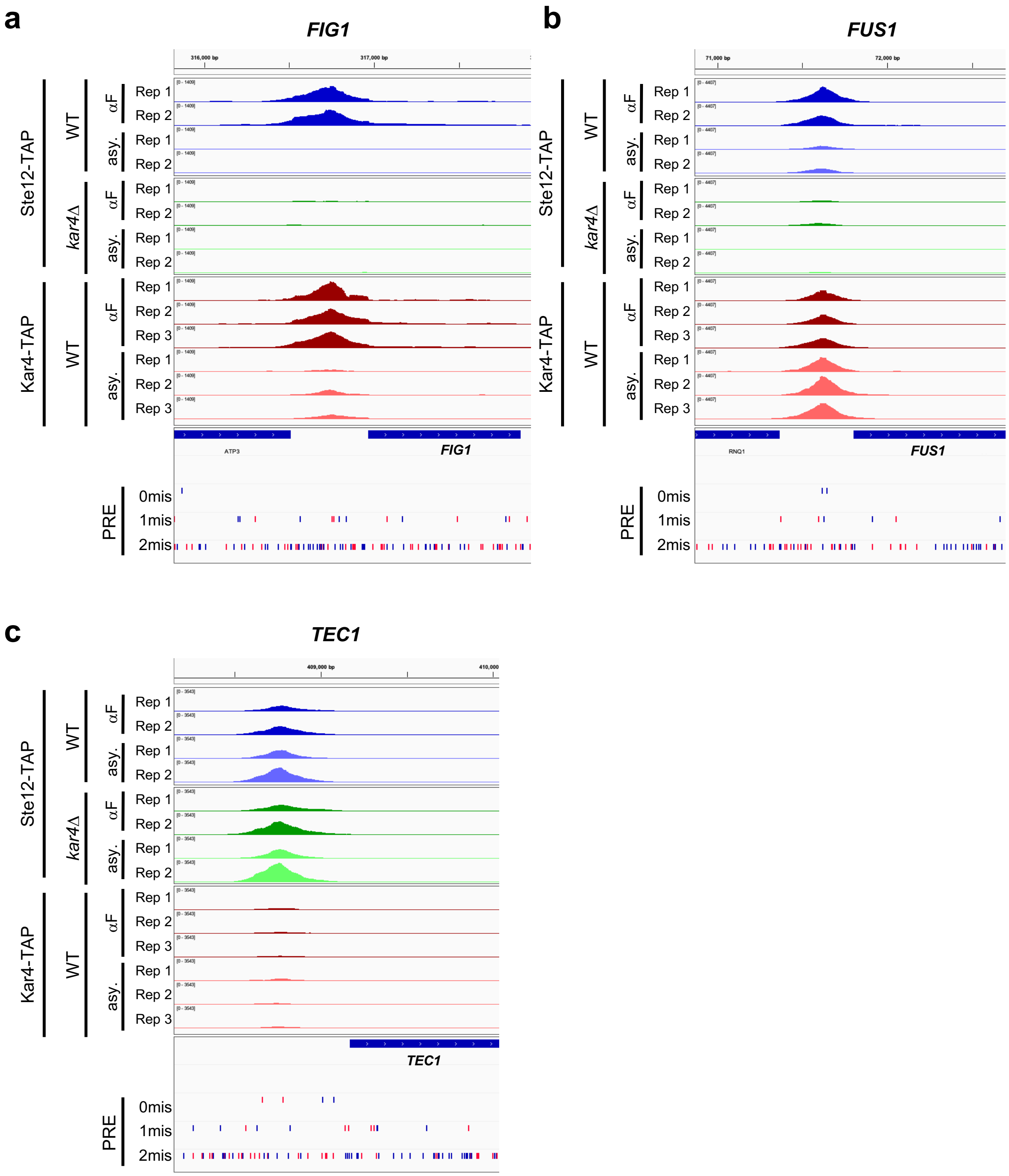

a
FIG1
316,000 bp
317,000 bp
Ste12-TAP
Kar4-TAP
WT
kar4Δ
αF
asy.
Rep 1
Rep 2
Rep 3
ATP3
FIG1
PRE
0mis
1mis
2mis
b
FUS1
71,000 bp
72,000 bp
RNQ1
FUS1
c
TEC1
409,000 bp
410,000
TEC1


Figure S7

**Supplementary Figure S7.** Example loci (IGV browser) as in **Figure 4a** showing ChIP-exo results for the indicated samples. Samples are depth-normalized and thus peak height values are directly comparable between tracks. PREs containing 0, 1, or 2 mismatches are displayed in the bottom 3 tracks, with matches on the forward strand in blue and reverse strand in red.

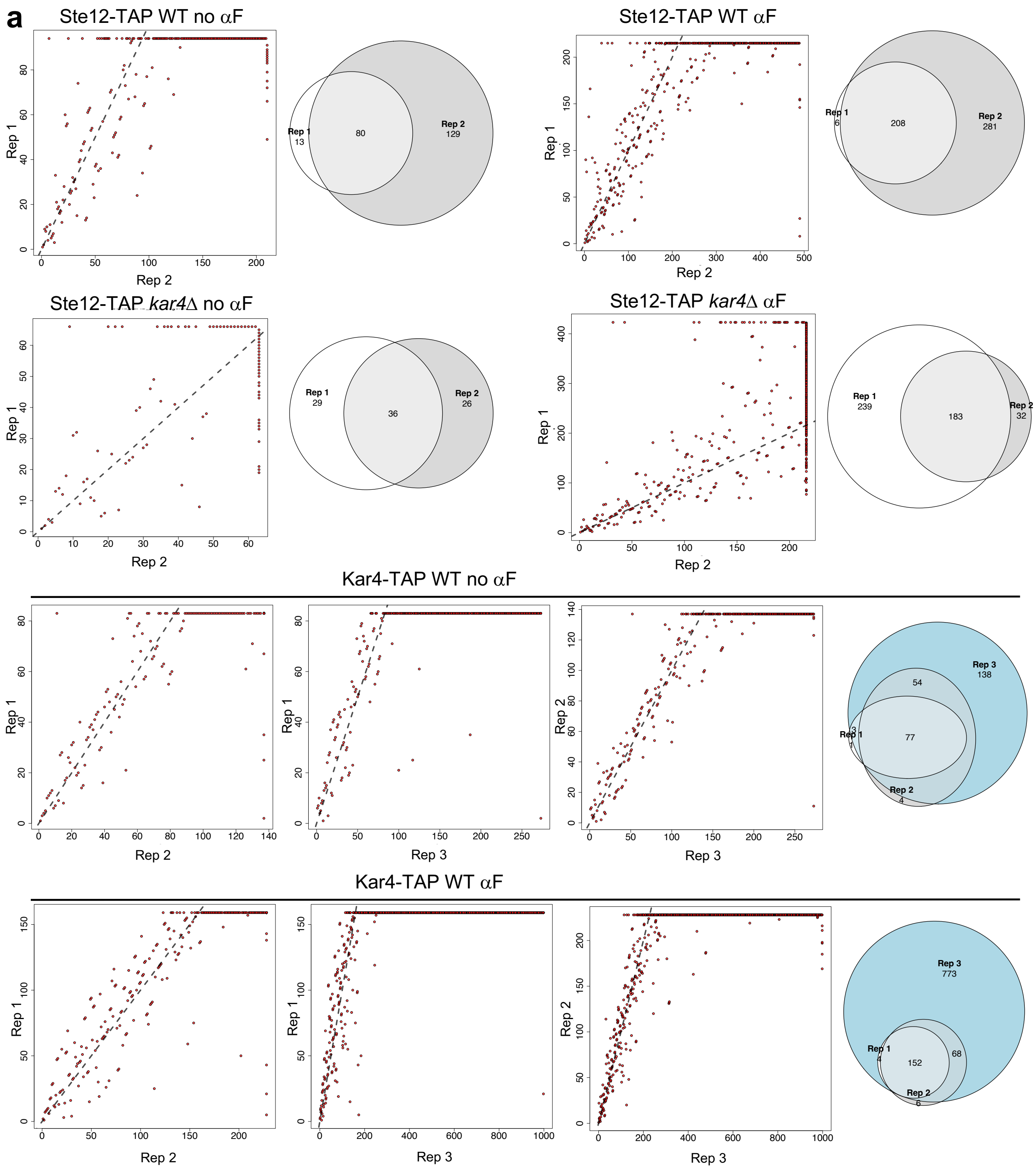


**b** Rank-correlation

| Group | Rep1 vs. Rep2 | Rep1 vs. Rep3 | Rep2 vs. Rep3 |
|---|---|---|---|
| Kar4 WT αF | 0.841 | 0.718 | 0.906 |
| Kar4 WT no αF | 0.880 | 0.724 | 0.937 |
| Ste12 *kar4*Δ αF | 0.854 | | |
| Ste12 *kar4*Δ no αF | 0.616 | | |
| Ste12 WT αF | 0.863 | | |
| Ste12 WT no αF | 0.833 | | |

**Figure S8**

**Supplementary Figure S8**. ChIP-exo replicate quality control. a) Replicate comparisons for the indicated groups. Genes were ranked by significance (q-value) of their associated peaks (tightly correlated with occupancy), with rank 1 being the most significant (or highest-occupancy). Genes found only in one replicate are given the same lowest value in the dataset from which they are missing, and are thus displayed on the edge of the plot. Gene set overlaps are shown in the Venn diagrams for each group. Note that genes were used instead of specific sites as exact peak locations between replicates varied, and one-to-one associations between peak locations were not always possible without first collapsing to nearby associated genes (see Methods). However, as some peaks were associated with multiple genes, these sites appear as diagonal streaks in the scatterplots (ranks are tied but for visualization displayed sequentially). Rarely, a peak at nearly the same location was associated with different genes in different replicates, leading to the appearance of extreme outliers. b) Heatmap of rank-based (Spearman) correlation values for each replicate comparison. Values were calculated using only the genes found in both replicates (no missing data), and when a single peak was associated with multiple genes it was only included once (no double-counting).

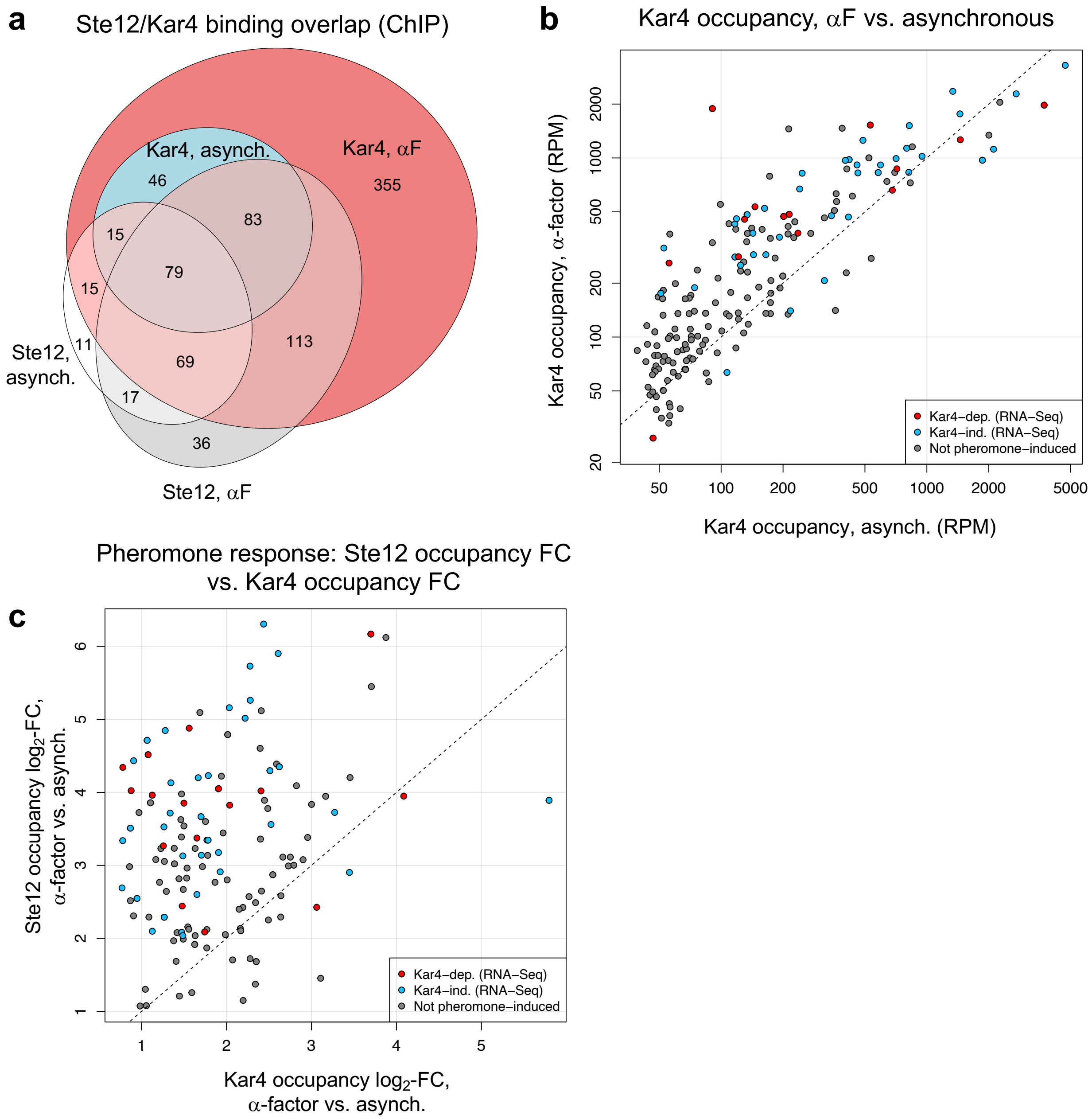

a
Ste12/Kar4 binding overlap (ChIP)
Kar4, asynch.
Kar4, αF
46
355
83
15
79
15
11
113
69
17
36
Ste12, asynch.
Ste12, αF
b
Kar4 occupancy, αF vs. asynchronous
Kar4 occupancy, α-factor (RPM)
Kar4 occupancy, asynch. (RPM)
Kar4−dep. (RNA−Seq)
Kar4−ind. (RNA−Seq)
Not pheromone−induced
c
Pheromone response: Ste12 occupancy FC vs. Kar4 occupancy FC
Ste12 occupancy log2-FC, α-factor vs. asynch.
Kar4 occupancy log2-FC, α-factor vs. asynch.
Kar4−dep. (RNA−Seq)
Kar4−ind. (RNA−Seq)
Not pheromone−induced


**Figure S9**

**Supplementary Figure S9.** a) Venn diagram indicating the overlap between all genes associated with Ste12 binding before (white) or after (grey) pheromone, and genes associated with Kar4 binding before (blue) or after (red) pheromone. Note that due to limitations of using Venn diagrams, three small intersections are not drawn: three genes found only in Kar4 asynchronous samples, two genes found in common only between Kar4 asynchronous and Ste12 pheromone-treated samples, and one gene found in common only between Kar4 asynchronous and Ste12 mitotic samples. b) Normalized peak height scatterplot as in **Figure 4d**, but for Kar4-TAP binding before and after pheromone. c) Fold-change in occupancy after pheromone treatment for genes with significantly increased occupancy for both Ste12 and Kar4. Red and blue points are pheromone-induced by RNA-seq and grey points are not. Dashed line indicates 1:1 line for equal values.

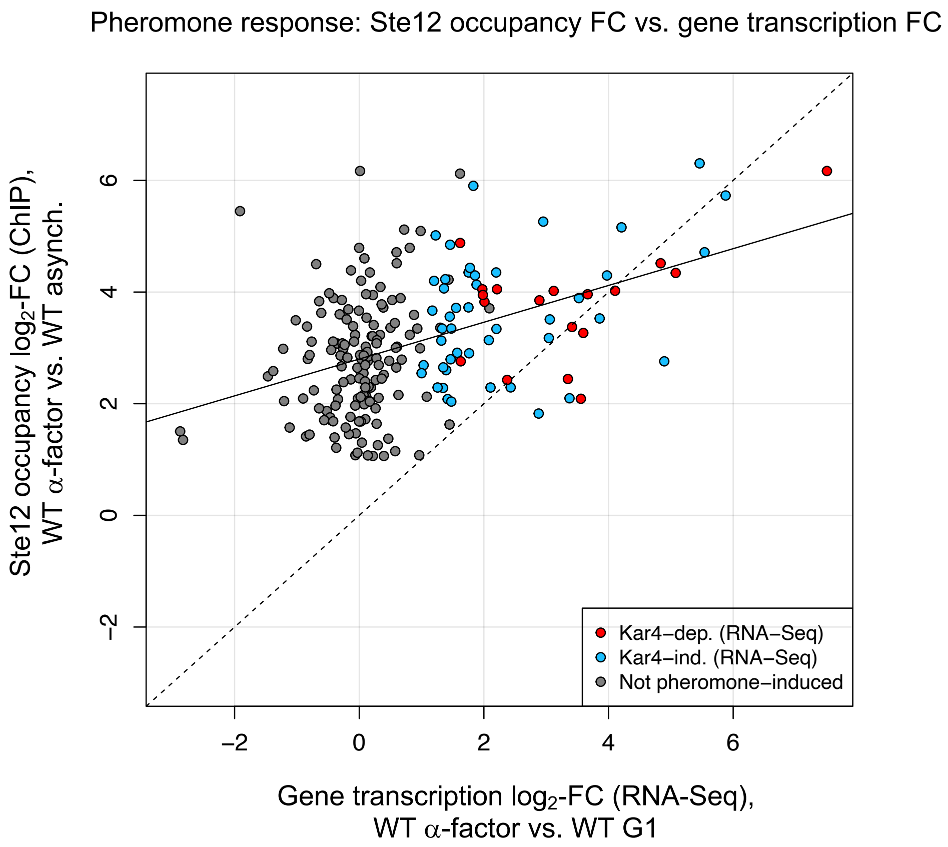
Pheromone response: Ste12 occupancy FC vs. gene transcription FC
Ste12 occupancy log2-FC (ChIP),
WT α-factor vs. WT asynch.
Gene transcription log2-FC (RNA-Seq),
WT α-factor vs. WT G1
Kar4−dep. (RNA−Seq)
Kar4−ind. (RNA−Seq)
Not pheromone−induced
−2
0
2
4
6

**Figure S10**

**Supplementary Figure S10**. Quantification of Ste12 occupancy change following pheromone treatment from ChIP-exo vs. transcriptional fold-change following pheromone treatment from RNA-seq. Points are colored as indicated; only genes associated with significantly increased Ste12 occupancy following pheromone in wild-type by ChIP are shown. Solid line indicates regression line. Dashed line indicates 1:1 line for equal values.

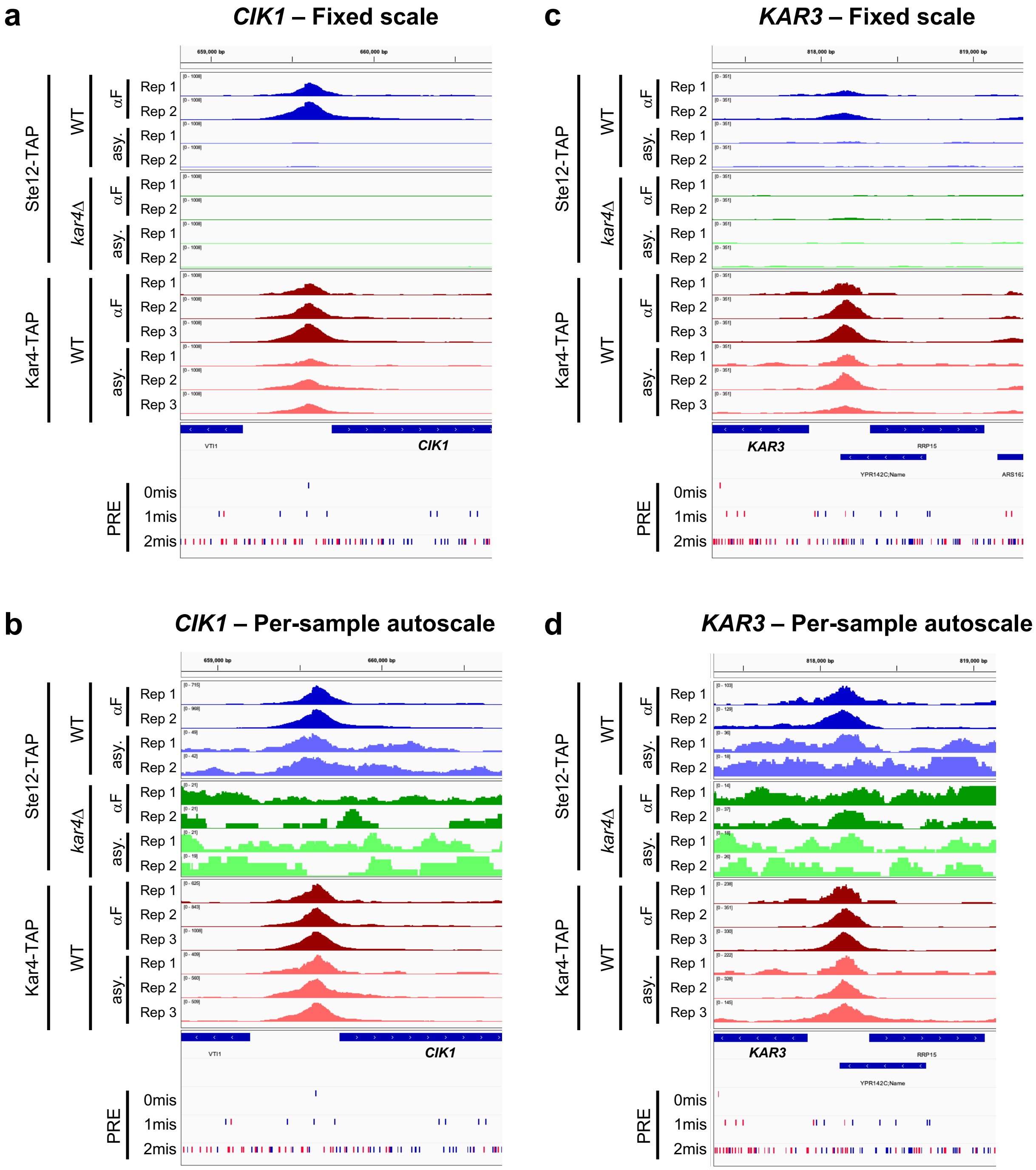
a
CIK1 – Fixed scale
b
CIK1 – Per-sample autoscale
c
KAR3 – Fixed scale
d
KAR3 – Per-sample autoscale
Ste12-TAP
Kar4-TAP
WT
kar4Δ
αF
asy.
Rep 1
Rep 2
Rep 3
VTI1
CIK1
KAR3
RRP15
YPR142C;Name
PRE
0mis
1mis
2mis

**Figure S11**

**Supplementary Figure S11.** Example Kar4-dependent loci, as in **Supplementary Figure S7**. a and b) *CIK1* locus shown with either a) fixed height scaling or b) per-sample autoscaling. c and d) As in a and b but for *KAR3* locus.

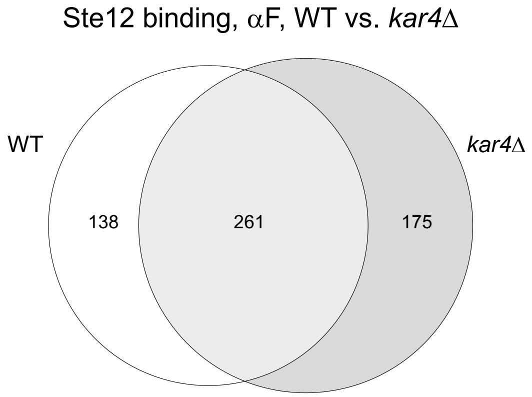


**Figure S12**

**Supplementary Figure S12.** Overlap between genes associated with Ste12 binding after pheromone treatment in wild-type and *kar4*Δ cells.

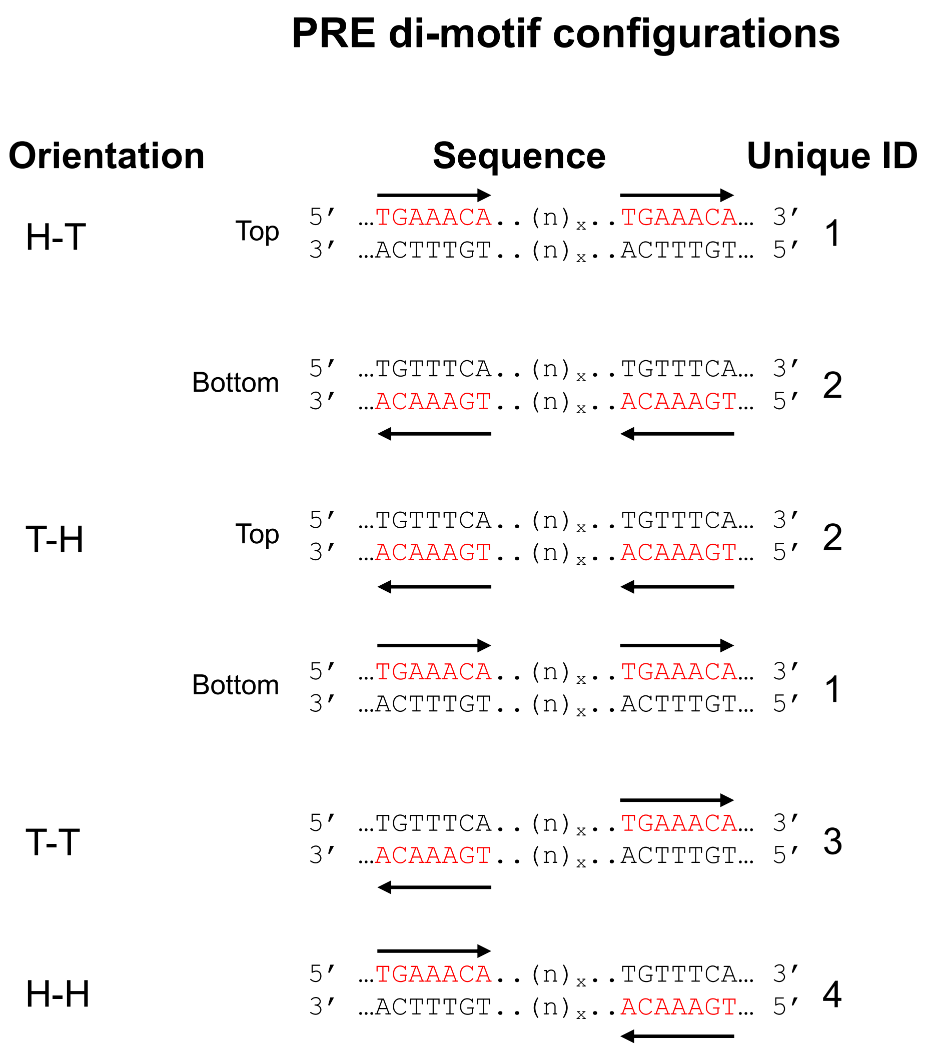
PRE di-motif configurations
Orientation
Sequence
Unique ID
H-T
Top
5′ …TGAAACA..(n)x..TGAAACA… 3′
3′ …ACTTTGT..(n)x..ACTTTGT… 5′
1
Bottom
5′ …TGTTTCA..(n)x..TGTTTCA… 3′
3′ …ACAAAGT..(n)x..ACAAAGT… 5′
2
T-H
Top
5′ …TGTTTCA..(n)x..TGTTTCA… 3′
3′ …ACAAAGT..(n)x..ACAAAGT… 5′
2
Bottom
5′ …TGAAACA..(n)x..TGAAACA… 3′
3′ …ACTTTGT..(n)x..ACTTTGT… 5′
1
T-T
5′ …TGTTTCA..(n)x..TGAAACA… 3′
3′ …ACAAAGT..(n)x..ACTTTGT… 5′
3
H-H
5′ …TGAAACA..(n)x..TGTTTCA… 3′
3′ …ACTTTGT..(n)x..ACAAAGT… 5′
4

**Figure S13**

**Supplementary Figure S13.** All possible PRE di-motif sequences, showing the matching sequence on either the top or bottom strand. Unique ID indicates which sequences are unique. Note that while H-T and T-H are the same as each other when the strand is flipped, T-T and H-H are palindromic and appear identical on both top and bottom strands. Sites are drawn with respect to the reference genome sequence, not relative to any downstream gene orientations.

Best PRE di-motif for Kar4-dependent Ste12 binding sites

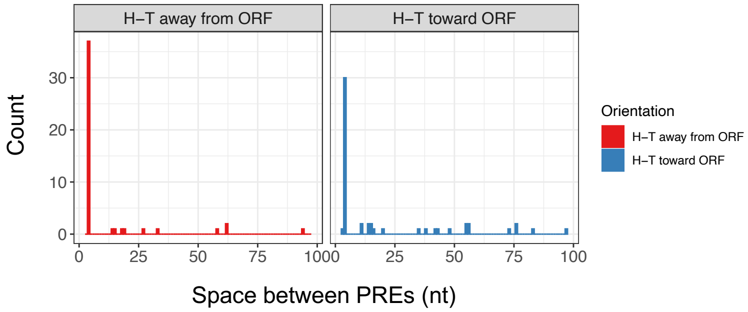


**Figure S14**

**Supplementary Figure S14.** Best PRE di-motif histogram as in **Figure 6c**, except the H-T di-motif category is separated by motifs facing toward or away from the nearby gene start codon.

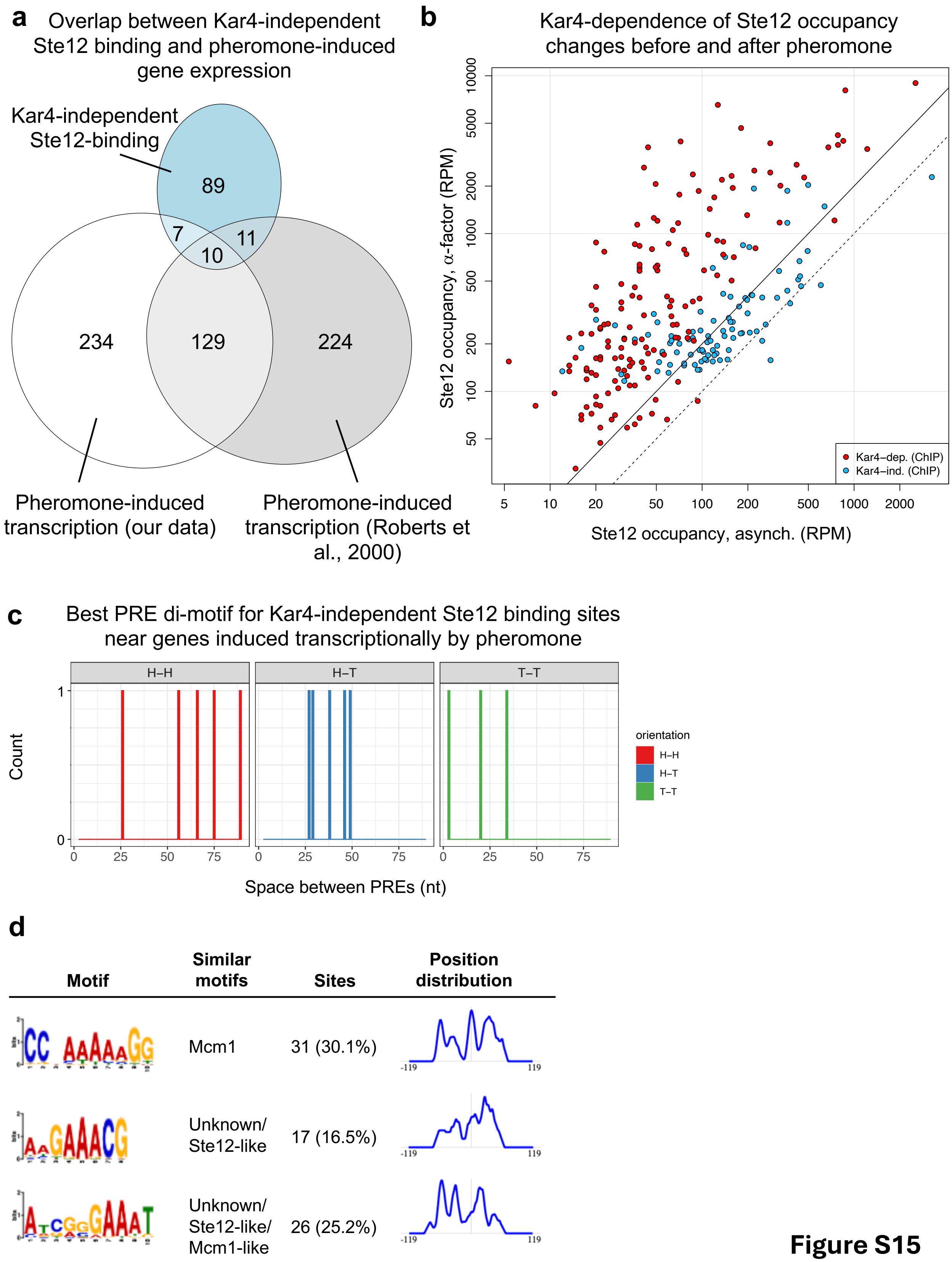

a
Overlap between Kar4-independent Ste12 binding and pheromone-induced gene expression
Kar4-independent Ste12-binding
89
7
11
10
234
129
224
Pheromone-induced transcription (our data)
Pheromone-induced transcription (Roberts et al., 2000)
b
Kar4-dependence of Ste12 occupancy changes before and after pheromone
Ste12 occupancy, α-factor (RPM)
10000
5000
2000
1000
500
200
100
50
Kar4–dep. (ChIP)
Kar4–ind. (ChIP)
5
10
20
50
100
200
500
1000
2000
Ste12 occupancy, asynch. (RPM)
c
Best PRE di-motif for Kar4-independent Ste12 binding sites near genes induced transcriptionally by pheromone
H–H
H–T
T–T
Count
1
0
0
25
50
75
Space between PREs (nt)
orientation
H–H
H–T
T–T
d
Motif
Similar motifs
Sites
Position distribution
Mcm1
31 (30.1%)
Unknown/ Ste12-like
17 (16.5%)
Unknown/ Ste12-like/ Mcm1-like
26 (25.2%)
-119
119


**Figure S15**

**Supplementary Figure S15.** a) Venn diagram comparing overlap of genes associated with Kar4-independent Ste12 binding (blue) to genes transcriptionally induced by pheromone in either our data (white) or Roberts et al. (2000) (grey). b) Normalized peak heights in reads per million (RPM) for Kar4-dependent (red) and Kar4-independent (blue) Ste12-binding regions from **Figures 6c and d**, respectively, before and after pheromone treatment in wild-type cells. Dashed line indicates 1:1 line for equal values. Solid line indicates a 2-fold increase in occupancy. Note that axes are spaced logarithmically; absolute unit values are displayed. c) Best PRE di-motif analysis as in **Figure 6d**, but restricted to Kar4-independent Ste12 binding sites associated with pheromone-induced genes (n = 13). d) De novo motif analysis as in **Figure 6b**, but for Kar4-independent Ste12-binding sites. For the bottom motif, Tomtom identified similarity to an Mcm1-associated motif, although by visual inspection it appears to be more similar to a PRE (Ste12-associated motif).

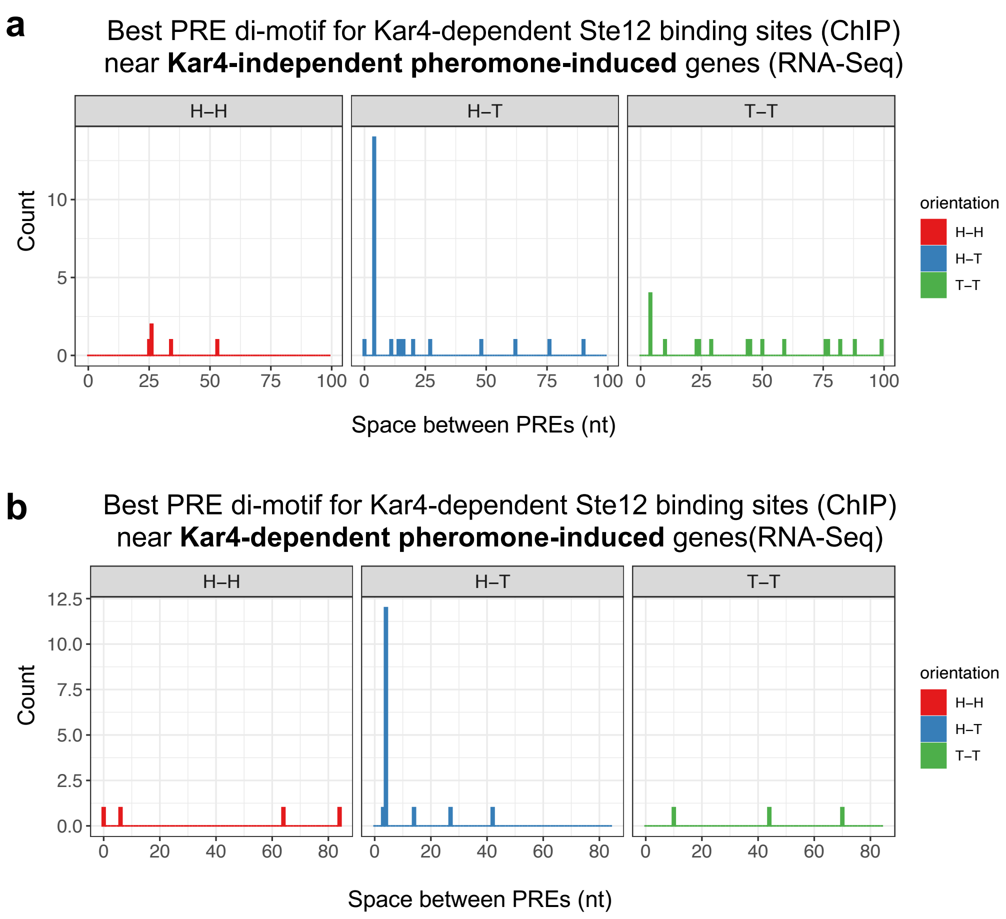
a
Best PRE di-motif for Kar4-dependent Ste12 binding sites (ChIP)
near Kar4-independent pheromone-induced genes (RNA-Seq)
H–H
H–T
T–T
Count
0
5
10
0
25
50
75
100
Space between PREs (nt)
orientation
H–H
H–T
T–T
b
Best PRE di-motif for Kar4-dependent Ste12 binding sites (ChIP)
near Kar4-dependent pheromone-induced genes(RNA-Seq)
H–H
H–T
T–T
Count
0.0
2.5
5.0
7.5
10.0
12.5
0
20
40
60
80
Space between PREs (nt)
orientation
H–H
H–T
T–T

**Figure S16**

**Supplementary Figure S16.** a) Best PRE di-motif analysis as in **Figure 6c**, but restricted to sites associated with genes classified as pheromone-induced in a Kar4-independent fashion by RNA-seq (n = 47). b) As in a, but restricted to sites associated with genes classified as pheromone-induced in a Kar4-dependent fashion by RNA-seq (n = 23).

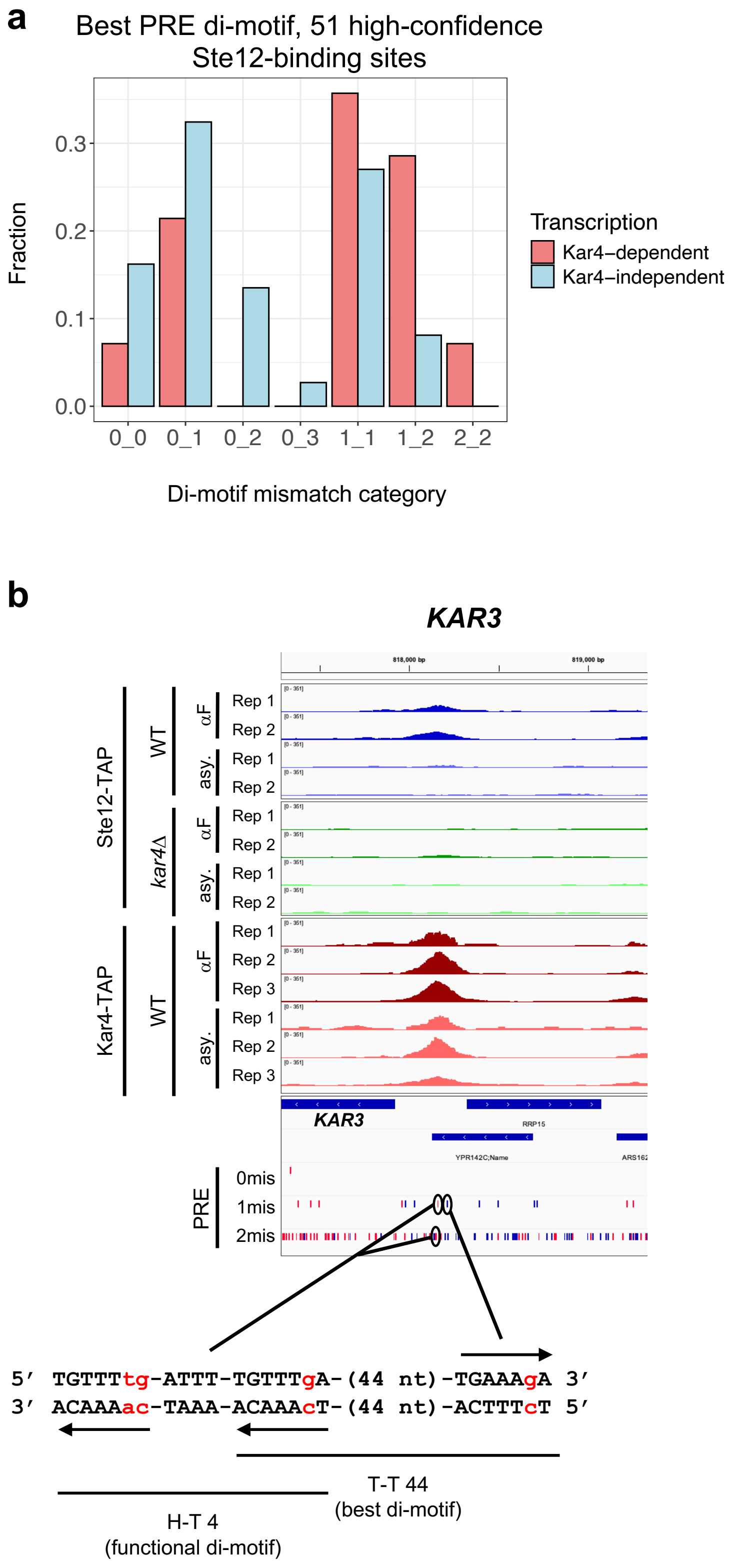
a
Best PRE di-motif, 51 high-confidence
Ste12-binding sites
Fraction
0.3
0.2
0.1
0.0
0_0
0_1
0_2
0_3
1_1
1_2
2_2
Di-motif mismatch category
Transcription
Kar4–dependent
Kar4–independent
b
KAR3
818,000 bp
819,000 bp
Ste12-TAP
WT
kar4Δ
Kar4-TAP
WT
αF
asy.
Rep 1
Rep 2
Rep 3
KAR3
RRP15
YPR142C;Name
PRE
0mis
1mis
2mis
5′ TGTTTtg-ATTT-TGTTTgA-(44 nt)-TGAAAgA 3′
3′ ACAAAac-TAAA-ACAAAcT-(44 nt)-ACTTTcT 5′
T-T 44
(best di-motif)
H-T 4
(functional di-motif)

**Figure S17**

**Supplementary Figure S17**. a) Mismatch histogram as in **Figure 8b** (70 motifs), but using the manually-curated subset of 51 motifs. b) Example di-motifs at *KAR3* promoter. ChIP-exo data are shown as in **Supplementary Figure S7**. Positions of motifs are circled. The center and right PREs form a T-T 44 motif, which was identified as the best di-motif in the region (each PRE has 1 mismatch), but the center and left PREs form an H-T 4 (although the left PRE has 2 mismatches). Arrows denote location and orientation of PREs.

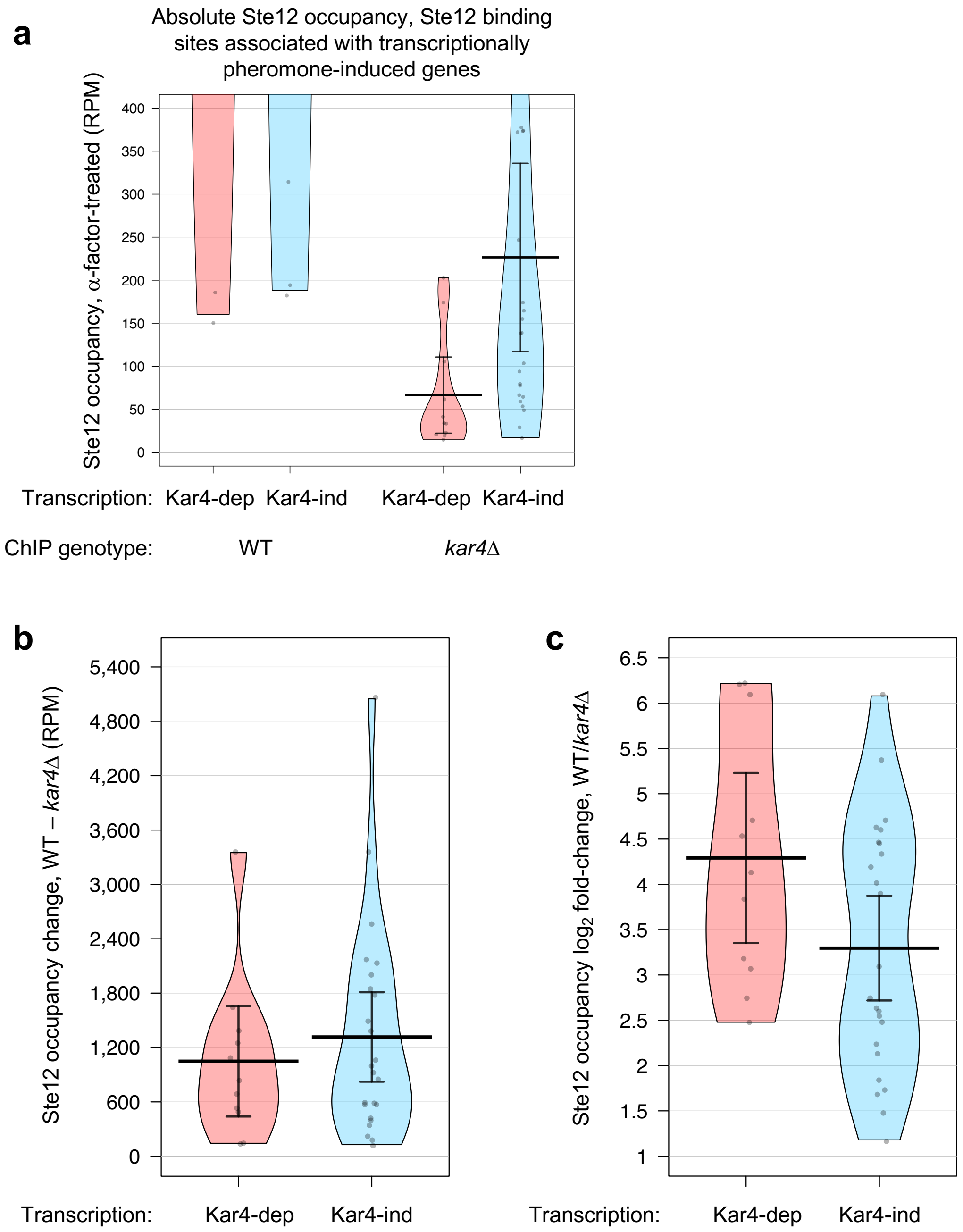

a
Absolute Ste12 occupancy, Ste12 binding sites associated with transcriptionally pheromone-induced genes
Ste12 occupancy, α-factor-treated (RPM)
400
350
300
250
200
150
100
50
0
Transcription: Kar4-dep Kar4-ind Kar4-dep Kar4-ind
ChIP genotype: WT kar4Δ
b
Ste12 occupancy change, WT − kar4Δ (RPM)
5,400
4,800
4,200
3,600
3,000
2,400
1,800
1,200
600
0
Transcription: Kar4-dep Kar4-ind
c
Ste12 occupancy log2 fold-change, WT/kar4Δ
6.5
6
5.5
5
4.5
4
3.5
3
2.5
2
1.5
1
Transcription: Kar4-dep Kar4-ind


**Figure S18**

**Supplementary Figure S18**. a) Same plot as **Figure 8d**, but zoomed-in on the y-axis to better visualize the datapoint values in *kar4*Δ cells. b) Ste12 occupancy metrics for the sites as in **Figure 8d**, but showing the change in occupancy between wild-type and *kar4*Δ for Ste12-binding sites associated with genes that are induced in a Kar4-dependent or independent manner, as indicated. Distributions are not significantly different ($p = 0.57$). c) As in panel a, but showing the $\log_2$ fold-change in Ste12 occupancy between wild-type and *kar4*Δ, as indicated. Distributions are not significantly different ($p = 0.076$). Statistical analyses were done using the Mann-Whitney U test.

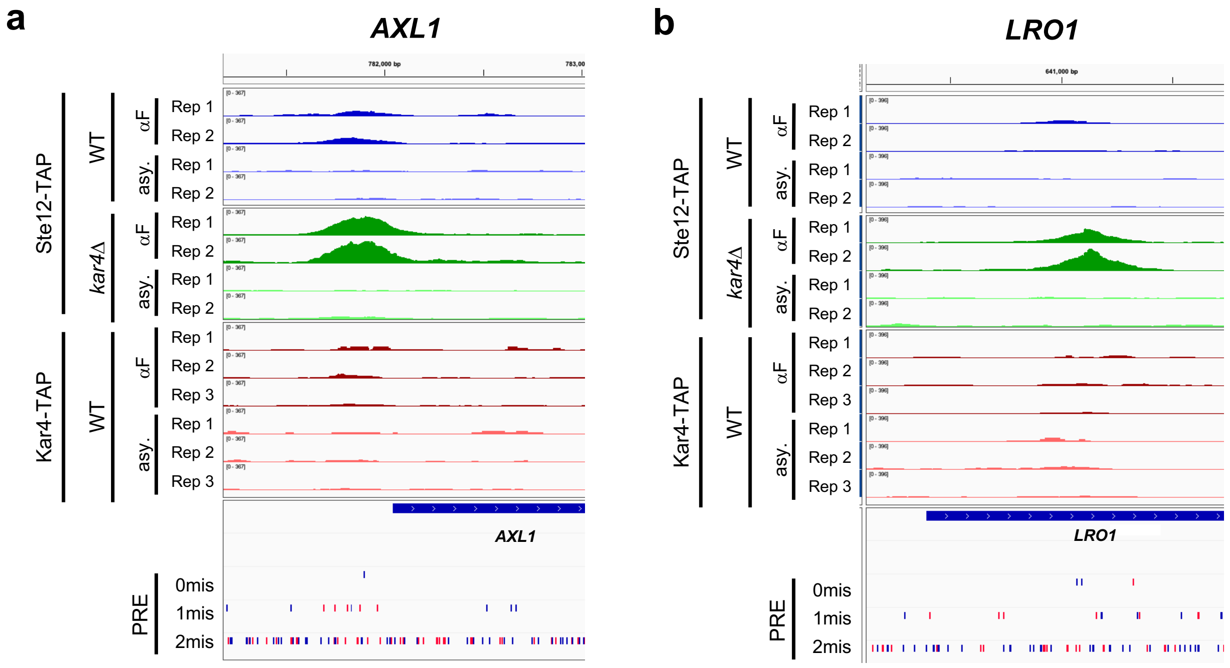
a
AXL1
782,000 bp
Ste12-TAP
WT
kar4Δ
Kar4-TAP
αF
asy.
Rep 1
Rep 2
Rep 3
AXL1
PRE
0mis
1mis
2mis
b
LRO1
641,000 bp
Ste12-TAP
WT
kar4Δ
Kar4-TAP
αF
asy.
Rep 1
Rep 2
Rep 3
LRO1
PRE
0mis
1mis
2mis

Figure S19

**Supplementary Figure S19.** Example loci as shown in **Supplementary Figure S7**, highlighting Ste12-binding peaks enriched in *kar4*Δ cells relative to wild-type after pheromone treatment.

#### *kar4Δ*-only gene promoters

**a** *kar4Δ*-only vs. Kar4-independent pheromone-induced

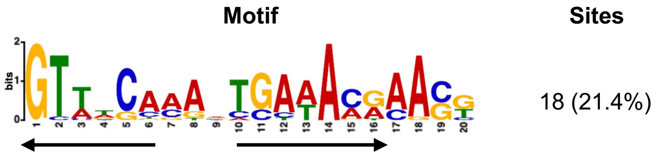


**b** Best PRE di-motif for *kar4Δ*-only gene promoters

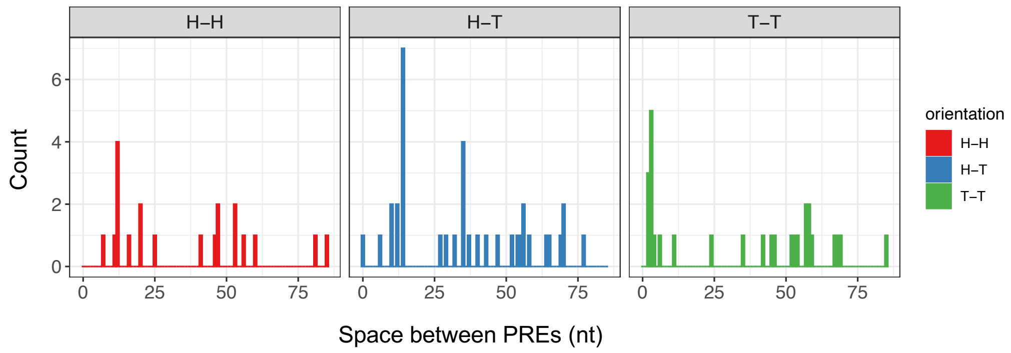


**Figure S20**

**Supplementary Figure S20.** a) de novo motif analysis (MEME) for the *kar4*Δ-only gene promoters vs. Kar4-independent pheromone-induced gene promoters, using pheromone-induced Kar4-independent genes as background. Arrows indicate location and orientation of the PREs. b) Histogram for best-match PRE di-motif (as in **Figure 6c**) for the 500 bp promoter regions associated with the *kar4*Δ-only genes from RNA-seq (n = 84).

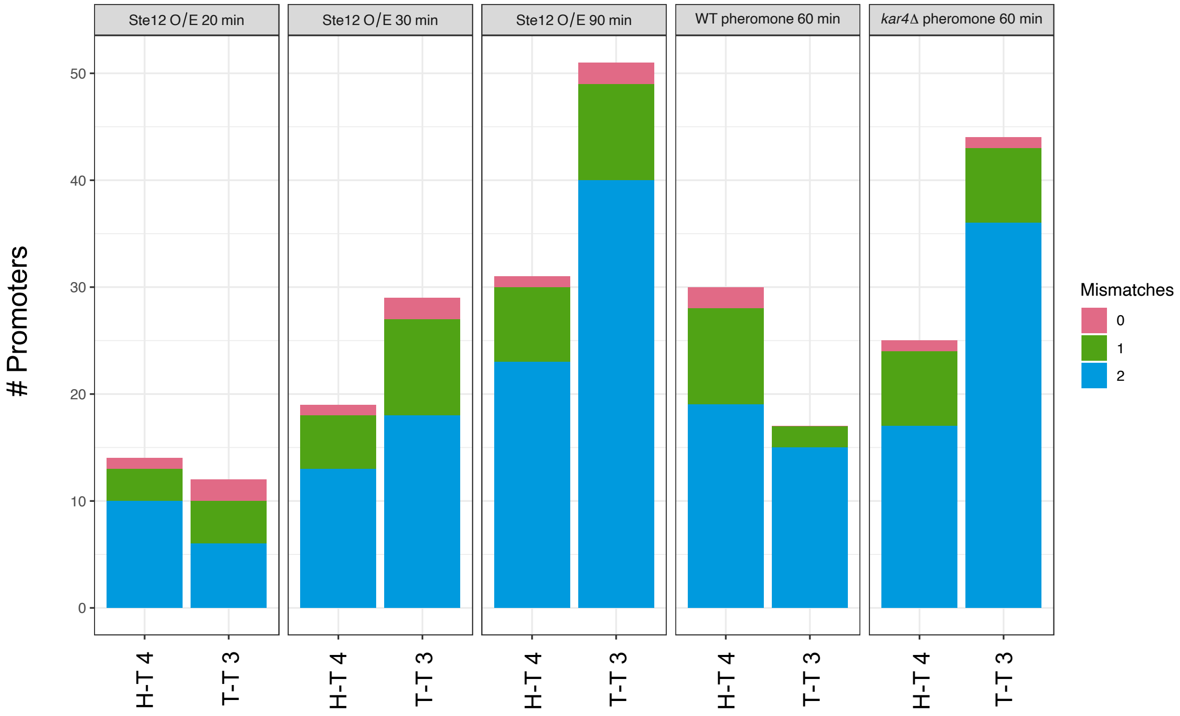

Ste12 O/E 20 min
Ste12 O/E 30 min
Ste12 O/E 90 min
WT pheromone 60 min
kar4Δ pheromone 60 min
### Promoters
0
10
20
30
40
50
H-T 4
T-T 3
Mismatches
0
1
2


**Figure S21**

**Supplementary Figure S21**. Quantification of the number of promoters of genes induced under the indicated condition (over expression (O/E) of Ste12 or pheromone addition) containing H-T 4 or T-T 3 motifs with the indicated number of mismatches (0, 1, or 2 mismatches). H-T 4 was defined using the full PRE (TGAAACA), T-T 3 was defined using the first 6 nt (TGAAAC), as this was the PRE associated with all T-T 3 motifs identified here as well in Dorrity et al. (2018). Each group includes only upregulated genes at ≥ 2 fold-change. Total number of genes in each group was 48 (*STE12* O/E 20 minutes), 134 (*STE12* O/E 30 minutes), 398 (*STE12* O/E 90 minutes), 385 (WT + pheromone 60 minutes), 488 (*kar4*Δ + pheromone 60 minutes).

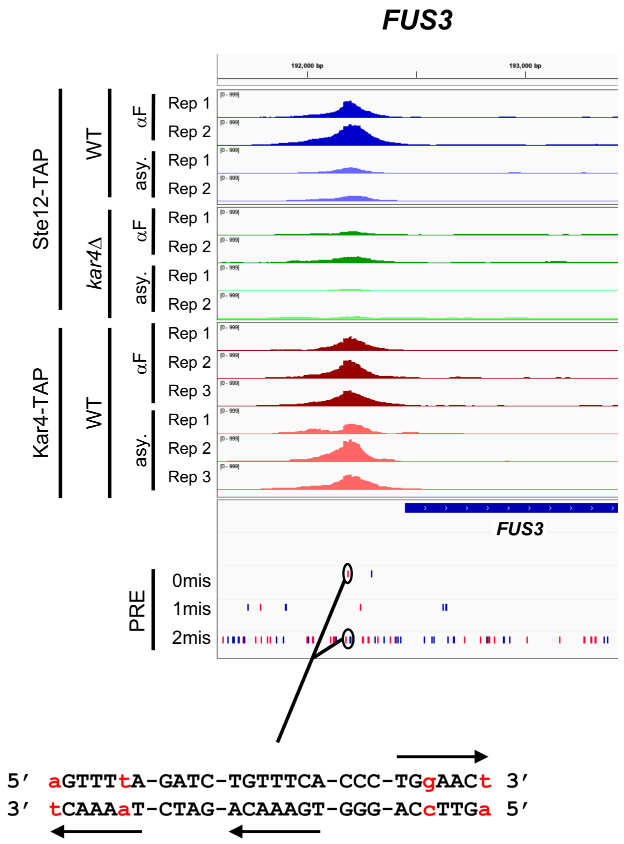

FUS3
192,000 bp
193,000 bp
Ste12-TAP
WT
kar4Δ
Kar4-TAP
αF
asy.
Rep 1
Rep 2
Rep 3
FUS3
PRE
0mis
1mis
2mis
5′ aGTTTtA-GATC-TGTTTCA-CCC-TGgAACt 3′
3′ tCAAAaT-CTAG-ACAAAGT-GGG-ACcTTGa 5′


Figure S22

**Supplementary Figure S22**. As in **Figure 10**, but for the *FUS3* promoter. The center and left PREs form an H-T 4 motif on the bottom strand, while the center and right PREs form a T-T 3 motif.
