## Appendix for "Kar4 acts as a Ste12 regulator in *Saccharomyces cerevisiae*, promoting Ste12 binding to a specific DNA motif genome-wide"

### Appendix: The pheromone response revisited

The main goal of this study was to understand the effect of Kar4 on Ste12 function. However, we additionally analyzed our RNA-seq and ChIP-exo data regarding various aspects of the mating response that are of general interest but unrelated to the Kar4/Ste12 relationship. Our observations include:

1. The calcineurin-responsive transcription factor, Crz1, induced a large proportion of pheromone-induced genes
2. Pheromone treatment can lead to the activation of the environmental stress response
3. A considerable number of genes are downregulated in response to pheromone through a complex regulatory network, independent of cell-cycle considerations
4. A Ste12-independent response to pheromone
5. Pheromone treatment leads to the transcription of intergenic transcripts.

It should be noted that the transcriptional response observed in this experimental system is a consequence of exposure to mating pheromone in the absence a mating partner. It remains to be determined whether, and to what extent, these responses occur under physiological settings. Nonetheless, the data provided here serve as a useful starting point to examine previously unknown and underappreciated aspects of the budding yeast mating response.

### Results & Discussion

#### Crz1 induces a large proportion of pheromone-regulated genes

Although Ste12 is required for nearly the entire transcriptional response during pheromone treatment, it is likely that a significant portion of gene expression changes are not due to direct activation by Ste12 but by downstream, secondary effects. This could be because some Ste12 targets are themselves transcription factors, or because the changes to cell physiology during the mating response trigger various cellular pathways. To gain an unbiased view of additional transcription factors acting during the pheromone response, we inferred relative transcription factor activity using Transfactivity, as described in the main text (**Figure 7a**). As expected, Ste12 was among the most significant explanatory transcription factors in our dataset. Ste12 activity was predicted to be highly increased during mating in wild-type, while in *ste12*Δ mutants the predicted activity was lower than the basal activity in wild-type and did not change in response to pheromone (**Figure 7b**). Surprisingly, however, in *STE12*+ cells two transcription factors stood out as by far the most significantly-altered activity (with similar p-values) in response to pheromone: Ste12 itself, and the calcineurin-responsive transcription factor Crz1 (**Figure 7a**). Calcium uptake has been shown to increase during pheromone treatment, activating calcineurin and leading to several physiological responses including Crz1 activation (OHSUMI AND ANRAKU 1985; IIDA *et al.* 1990; MATHEOS *et al.* 1997). Crz1 activation occurs via calcineurin-mediated dephosphorylation of Crz1, which then translocates to the nucleus and induces gene expression (STATHOPOULOS-GERONTIDES *et al.* 1999). Moreover, *crz1*Δ mutants have a survival defect in the presence of mating pheromone (STATHOPOULOS AND CYERT 1997). Using cells lacking calcineurin, this survival defect has been attributed to a build-up of mitochondrial-respiration-generated reactive oxygen species (ZHANG *et al.* 2006). However, the extent of the contribution of Crz1 to the pheromone response has not, to our knowledge, been previously reported. Here we found that although the level of *CRZ1* mRNA did not increase in response to pheromone (**Supplementary File S2**), Crz1 activity increased, as inferred by activation of genes containing a Crz1 motif in their promoter, in a manner that was Ste12-dependent and Kar4-independent (**Appendix Figure A1**). We further found that Crz1 activity was slightly downregulated in *ste12*Δ cells in the absence of pheromone, implying a constitutive role for Ste12 in regulating Crz1 activity. However, expression of *CRZ1* mRNA was not affected, suggesting that a Ste12 target affects Crz1 activity, rather than regulating Crz1 levels directly.

To estimate the fraction of pheromone-responsive genes attributable to Crz1 signaling, we compared our data to a Crz1 overexpression timecourse dataset from the same study used in the main text for Ste12 overexpression (HACKETT *et al.* 2020) (**Appendix Figure A2**). To identify all gene expression changes attributable to Crz1 activity, we classified genes as Crz1 targets if their mean fold-change in the Crz1-overexpression timecourse was at least 2-fold upregulated at all timepoints starting at 30 minutes of Crz1 induction. Under these criteria, 199 genes were induced by Crz1. This list of genes includes 31% (52/167) of our pheromone-induced Kar4-independent gene list (p-val = $4.05 \times 10^{-40}$) and 21% (10/47) of our pheromone-induced Kar4-dependent gene list (p-val = $1.01 \times 10^{-6}$), compared to only 7% (6/84) of the *kar4*Δ-only genes (p-val = 0.04) and 2% of all background genes. For the 10 Kar4-dependent genes (*SUL1*, *PHO89*, *YDR034W-B*, *YGL052W*, *PRM8*, *JLP1*, *HXT2*, *SPG4*, *YNR064C*, and *PRM4*), we suspect that Crz1 and Ste12 function independently and additively on their promoters (rather than functioning cooperatively), as pheromone-dependent induction of these genes is reduced but not ablated in the *kar4*Δ mutant. Intriguingly, 181 genes were downregulated by Crz1 overexpression, including 18% (22/119) of the genes downregulated during the pheromone response (p-val = $6.86 \times 10^{-13}$), suggesting both upregulated and downregulated genes during the pheromone response can be attributed to Crz1 signaling.

We also wondered whether Ste12 and Crz1 regulated distinct sets of target genes, or whether there was any overlap which might suggest functional redundancy or even cooperativity. Using our ChIP-exo data, we found that Ste12 binding is associated with 10% (20/199) of genes induced by Crz1. Although modest, the overlap was significant with a p-value of 0.018. Because Ste12 activity should be largely inhibited by Dig1 and Dig2 during the Crz1 overexpression timecourse, which was performed in cycling cells in the absence of pheromone, it is likely that these genes are induced redundantly by Crz1, rather than being cooperatively expressed via a Ste12-Crz1 interaction. Intriguingly, several of these 20 genes (*FUS1*, *RVS161*) were identified in a screen for genes required for activation of the low-affinity $Ca^{2+}$ influx system during the mating pheromone response (MULLER *et al.* 2003), suggesting that the genes regulated by both Crz1 and Ste12 may have multiple roles during mating. Overall, however, these results suggest that Ste12 and Crz1 generally do not directly regulate the same genes. This is further consistent with the observation above that *kar4*Δ-only genes, which were demonstrated in the main text to be ~50% or more direct Ste12 targets, were generally not induced by Crz1 (6/84 genes, p-val = 0.04). Therefore, although Ste12 is ultimately required for Crz1 activation in response to pheromone, Crz1 is directly responsible for ~1/3 or more of the genes upregulated in the normal response to pheromone, a value similar to or greater than the number of genes directly induced by Ste12. Given the shared importance of calcium signaling during fertilization and zygote formation in other species (STRICKER 1999), better characterizing the role of Crz1 during yeast mating in future studies may prove broadly informative.

**The environmental stress response is activated during exposure to pheromone**

Even after accounting for direct Ste12 and Crz1 targets, there were still hundreds of genes that are differentially expressed during pheromone treatment. Much of this response has previously been attributed to a difference in cell-cycle stage caused by the experimental design, where the control cells were asynchronous while the pheromone-treated samples were arrested in G1. For example, when asynchronous cells are treated with pheromone, some of the strongest downregulated targets include the histone genes, which are normally expressed only during S-phase (ROBERTS *et al.* 2000). This was explained by the depletion of cells in S-phase in the pheromone-responsive cells (which arrest in G1), rather than an actual consequence of pheromone-regulated transcription (ROBERTS *et al.* 2000). Consistent with this interpretation, in our dataset, where cells were pre-arrested in G1, we saw no significant expression changes in the histone genes (except *HHO1*, which is upregulated ~4-fold (**Supplementary File S2**)). Conversely, the comparison in previous studies between untreated asynchronous cells and pheromone-treated G1 cells also likely masked real but weak pheromone-dependent changes in gene expression. Given that in our experimental system, both control and pheromone treated cells are in G1, we sought an explanation other than cell-cycle changes for how hundreds of genes are changing expression during pheromone treatment.

The environmental stress response (ESR) is a regulon containing hundreds of genes that change synchronously in response to nearly all environmental perturbations. Given the preponderance of ribosome genes, which are part of the ESR, in our set of pheromone-dependent down-regulated genes (especially noticeable in the *kar4*Δ mutants), we reasoned that the ESR might be triggered in response to pheromone as well, perhaps in response to morphological changes such as cell-wall remodeling (GASCH *et al.* 2000). To quantify the strength of the ESR in our dataset, we used a PCA-based method developed by quantifying the universal ESR signal from a compendium of ~1500 deletion mutant strains (KEMMEREN *et al.* 2014; O'DUIBHIR *et al.* 2014). Briefly, ~1500 deletion mutant strains were chosen for transcriptome profiling based on having potential roles in regulating gene expression. Then, it was found that the first principal component of the transcriptome compendium (accounting for 24% of the total variance) corresponded to a slow-growth signature (O'DUIBHIR *et al.* 2014) which matched gene expression signatures found in earlier studies on the environmental stress response (GASCH *et al.* 2000). Therefore, the magnitude of the ESR can be estimated in any gene expression dataset by calculating the similarity between this principal component vector and a given set of gene expression fold-changes (see Methods). Using this method, we found that the ESR was significantly induced in response to pheromone treatment in a manner that was largely dependent on Ste12 (**Appendix Figure A3**). However, as the ESR induction was not entirely Ste12-dependent, this suggests that some of it can be attributed to a Ste12-independent transcriptional response (characterized below). We also considered that ESR induction could arise as a response to the increased time (1 hour) that the pheromone-treated cells were arrested in YPD. However, even when this was controlled for by using a +60 minute no-pheromone sample as the baseline, the ESR was still observed in response to pheromone and at a similar magnitude (**Appendix Figure A3**, rightmost two bars). Therefore, this effect appears to be a genuine aspect of pheromone-induced transcriptional changes using our experimental design and not an experimental artifact. These results fit with our previous observation by GO analysis (main text) that mating is a stressful process. Elucidating the nature of this stress (cell wall remodeling, changes in biomass production rate, etc.) could prove informative for a better understanding of the mating response at a physiological level.

**Genes downregulated during pheromone treatment arise from a complex regulatory landscape**

As noted earlier, previous work has largely attributed genes downregulated during pheromone treatment to cell-cycle effects. Because our experimental design allowed us to exclude cell-cycle effects from our dataset, we sought to identify the transcription factors underlying the observed downregulated gene expression. During our analysis of Crz1 targets, we observed that a significant fraction (~18%) of the downregulated genes also decrease expression during *CRZ1* overexpression (**Appendix Figure A2b**), suggesting that Crz1 could be responsible for both upregulated and downregulated genes during pheromone treatment. Stb3 and Dot6 are repressors of ribosome biogenesis genes, and their repressive activity increased significantly in response to pheromone, albeit more so in the *kar4*Δ strain (**Figure 7a**). This result is consistent with our GO analysis revealing an enrichment of ribosome biogenesis genes in the downregulated gene list (**Supplementary File S4**). Likewise, a positive regulator of ribosomal and ribosome biogenesis genes, Sfp1, was predicted to decrease activity in both wild-type and *kar4*Δ cells (**Figure 7a)**, consistent with a downregulation of ribosomal proteins. These results were further corroborated by the pheromone-dependent general increase in ESR noted above (**Appendix Figure A3**), as downregulation of ribosome biogenesis and ribosomal protein genes is a core feature of the ESR.

In addition to these observations, we searched directly for known transcription factor motifs in the promoters of the downregulated genes. Unlike the upregulated genes, no single motif was evident; however motifs associated with 8 different transcription factors were weakly enriched, including Mot2, Pdr1, and Nrg2 (**Appendix Figure A4**). These results suggest that downregulated genes arise from a multitude of transcription factors, whose activities likely respond secondarily after the initial wave of Ste12 activity during mating.

**A Ste12-independent pheromone response**

An unexpected cluster of genes were upregulated in response to pheromone in all strains tested, including *ste12*Δ (**Figure 2,** cluster IV). This was not an artifact of cells spending an extra 60 minutes in G1, as these genes were similarly upregulated when the 60-minute no-pheromone condition was used as a reference (**Supplementary File S2**). The majority of these genes were weakly upregulated, and only 23 increased expression over 2-fold in all pheromone-treated conditions (**Figure 1b**). Nonetheless, 352 genes did exhibit significant upregulation (p-adj < 0.01) in response to pheromone in all genotypes (**Supplementary File S2**). GO analysis of these genes revealed a strong enrichment for terms related to respiration and mitochondria, oxidoreductase activity, cell wall-related biological processes and for proteins localizing to the cell periphery and vacuole (**Supplementary File S4**), raising the intriguing possibility of a genuine Ste12-independent response to pheromone. This could arise either from Ste12-independent effects downstream of pheromone binding to its normal receptor, or from off-target effects of pheromone binding to other plasma membrane receptors. As Ste2 (the alpha-factor receptor) mRNA expression is only reduced to ~25% its normal basal expression in *ste12*Δ G1-arrested cells (**Supplementary File S2**), this cluster of genes could arise from Ste12-independent effects of the Ste2-mediated pheromone response. For example, both Fus3 (the primary MAP kinase mediating the pheromone response) and Kss1 (another MAP kinase activated by pheromone) phosphorylate numerous target proteins unrelated to Ste12 activation (CHEN AND THORNER 2007). Additionally, there is known cross-talk between the pheromone-signaling cascade and other stress-responsive MAPK pathways (VAN DROGEN *et al.* 2020). In this regard, we note that activity associated with Rlm1, a component of protein kinase C-mediated cell integrity signaling, is predicted to increase in response to pheromone in all genotypes (**Figure 7a**). This would also fit with our observation that the ESR signal increases somewhat in *ste12*Δ cells in response to mating pheromone, as would be expected from activation of the cell integrity pathway (**Appendix Figure A3**). However, previous work, which similarly found an activation of cell integrity signaling (as measured by Mpk1 activity) after pheromone treatment in G1-arrested cells, found that cell integrity pathway activation was strictly dependent on the Ste12-dependent transcriptional response (BUEHRER AND ERREDE 1997). This suggests that the Ste12-independent pheromone response could also be affected by other unknown experimental factors.

**Intergenic RNA transcripts are transcriptionally regulated during the pheromone response, many directly by Ste12**

Our RNA-seq allowed for the analysis of all transcribed RNAs, including unannotated genes and intergenic non-coding RNAs, during the pheromone response. To identify novel transcripts, we looked for transcripts in intergenic regions (which are therefore likely to be non-coding RNAs or short unannotated proteins) and quantified their response to mating pheromone in wild-type, *ste12*Δ, and *kar4*Δ strains. Briefly, intergenic transcripts were defined by predicting all transcripts de novo, then retaining only those that were > 200 bp in length and not overlapping known transcribed elements, then merging overlapping predicted transcripts (see Methods for more details). In total, we identified 684 intergenic transcripts (ITs) (**Appendix File A1**), of which 42 were significantly pheromone-induced (≥ 2-fold) in a Ste12-dependent manner, and 22 were pheromone-repressed (≤ 2-fold). Similarly to our gene analysis, we identified a Kar4-dependent class (10 pheromone-induced ITs, corresponding to 24% (10/42) of total pheromone-induced ITs, compared to 22% of pheromone-induced standard genes which were Kar4-dependent). We also identified a *kar4*Δ-only class (25 pheromone-induced ITs, of which 2 were also upregulated in wild-type cells).

We next determined which of the ITs are associated with DNA-bound Ste12 using our ChIP-exo data. Following pheromone treatment, we identified 70 ITs associated with Ste12 binding in wild-type, of which 25 transcripts were Kar4-dependent for Ste12 binding, and independently we identified 16 transcripts that exhibited higher Ste12 occupancy in a *kar4*Δ cells (*kar4*Δ-only) **(Appendix File A2**). Although about half of the peaks were shared bidirectionally between the intergenic transcript and a strongly pheromone-induced gene (**Appendix Figure A5a and b**), there were many examples where Ste12 binding was far away from any annotated genes (for example, see **Appendix Figure A5c**).

The overlap between the ITs that were associated with Ste12 binding and the ITs that were transcriptionally induced by pheromone was modest and similar to what we observed for protein-coding genes (**Appendix Figure A6a and b**, compare to **Figure 5b**). Of the 11 ITs which exhibited Ste12-binding and transcriptional upregulation during pheromone treatment, most were upregulated ~6-fold or less, but two transcripts were upregulated more than 20-fold: MSTRG.81 and MSTRG.403. MSTRG.81 corresponds to a previously-characterized long non-coding RNA (lncRNA) that represses HO transcription following pheromone treatment (YU *et al.* 2016). Expression of this lncRNA was previously shown to be Ste12-dependent and to require a perfect PRE sequence ~2.7 kb upstream of HO (YU *et al.* 2016). We identified Ste12 binding to this same site, which forms an H-T 4 motif with a nearby 2-mismatch PRE (TaAAACc) (**Appendix Figure A7a**). We additionally observed that both Ste12 binding at this site and transcription of the IT are Kar4-dependent (**Appendix Figure A7a** and see **Appendix Files A1** and **A2**). MSTRG.403 is Kar4-independent for Ste12 binding and transcription and is antisense to the gene *MCH2* (**Appendix Figure A7b**). However, *MCH2* expression does not change during pheromone treatment in our dataset. Non-coding RNA transcription at this locus has not, to our knowledge, previously been characterized. Although expression of this IT changes dramatically during pheromone treatment, this may represent a case of spurious transcription occurring at a good H-T 4 site that is coincidentally positioned in a genomic context favorable for transcription.

Lastly, we analyzed the Ste12 binding sites associated with pheromone-induced ITs for the presence of PRE di-motifs, as well as de novo motifs. This analysis was done for the Kar4-dependent and *kar4*Δ-only sites, which are more likely to be relevant to the pheromone response. For the sites exhibiting Kar4-dependent Ste12-binding near ITs, a weak H-T 4 PRE di-motif was identified by MEME in 100% of the sequences (**Appendix Figure A6c**). When explicitly searching for PRE di-motifs, the H-T 4 motif was the best match in 22% of sites (**Appendix Figure A6e** and **Appendix File A3**), a value somewhat lower than the 44% found for standard protein-coding genes. 33% of the best-match PRE di-motifs contained a perfect PRE, similar to our results for standard protein-coding genes (36%). For *kar4*Δ-only sites, only the PRE mono-motif (as opposed to any di-motif configuration) was found as significant de novo (**Appendix Figure A6d**). For the di-motif analysis, while 31% of the best di-motifs contained a perfect PRE, no di-motif was found more than once (**Appendix Figure A6f**). The T-T 3 motif, which was enriched in our previous *kar4*Δ-only Ste12 binding site analysis on protein-coding genes, was identified once, while the H-T 4 motif was not identified at all.

The significance and potential function of these pheromone-induced intergenic transcripts remain to be further explored. Although intriguing, it is possible that for most intergenic transcripts their transcription is largely fortuitous and non-functional. However, as identifying even a single intergenic transcript essential for mating would add to our understanding of transcriptional regulation during the pheromone response, exploring this further remains important.

### Appendix Materials & Methods

#### ESR strength prediction

To predict the magnitude of the environmental stress response (ESR) in our samples, we used a PCA-based method as previously described (O'DUIBHIR *et al.* 2014). Briefly, in this method, the environmental stress response was defined as the first principal component in a gene expression dataset comparing ~1500 single-deletion mutants to wild-type. Therefore, the contribution of every gene toward the ESR is captured in a single vector. The dot product was then calculated between this ESR vector and the $\log_2$ fold-changes in each of our samples and reported as the ESR strength. For this analysis, because there were minor differences in baseline ESR strength between our wild-type batches, we normalized each set of duplicates against its own wild-type control (namely, whereas in **Figure 2** samples were normalized against the average of four wild-type T0 controls, for **Appendix Figure A3** they were normalized against the average of two wild-type T0 controls each (either batch 1 or batch 2)).

#### Enrichment of known transcription factor motifs

Enrichment analysis of known motifs (**Appendix Figure A4)** was performed similarly to *de novo* motif identification, except instead of STREME we used SEA (https://meme-suite.org/meme/tools/sea) (BAILEY AND GRANT 2021) with default settings and the YEASTRACT database for reference motifs. Promoter sequence for all non-queried genes was used as background.

#### Identification of candidate novel intergenic transcripts

Candidate novel intergenic transcripts (ITs) were predicted using a custom pipeline (DOI: 10.5281/zenodo.17703432). Briefly, an initial set of transcripts were predicted genome-wide from RNA-seq BAM files with *stringtie* (v2.2.1) (PERTEA *et al.* 2015) in stranded mode and with a minimum transcript length cutoff of 200 bases. Next, a preliminary set of novel ITs per sample was generated by removing predicted transcripts overlapping known genes on the same strand or annotated LTR repeats with the program *bedtools intersect* (v2.31.1) (QUINLAN AND HALL 2010). Finally, all candidate ITs overlapping more than 10% from each sample were merged with *stringtie --merge* to generate a final set of candidate novel ITs. To further evaluate the validity of this final set of ITs, we searched for the presence of transcription start sites predicted by Cap analysis gene expression from the YeasTSS database (MCMILLAN *et al.* 2019) with *bedtools intersect*. Transcripts overlapping UTRs previously annotated (NAGALAKSHMI *et al.* 2008) were flagged.

**Differential expression analysis of predicted intergenic transcripts**

Fragment counts per predicted IT per sample were generated from the RNAseq BAM files previously used for gene differential expression analysis, with the *featureCounts* tool of the *Subread* package (v2.0.6) with the parameters "*-t exon -g gene_id -O -s 2 -J -R BAM -p --ignoreDup -M –fraction",* discarding duplicated and multimapping reads. Differential expression analysis of ITs was carried out with a custom R (v4.3.1) script (DOI: 10.5281/zenodo.17703432) based on the R package DESeq2 (v1.42.1). Classification as Ste12-dependent, Kar4-dependent, etc., was performed as described in the Methods in the main text.

**Associating Ste12 and Kar4 DNA binding sites with intergenic transcripts**

For ChIP-exo analyses related to ITs, ITs were included in the standard pool of candidate genes when associating binding sites with genes or LTRs. All downstream analyses were then unaltered. For analyses in the main results, ITs were not included in the gene association pool.

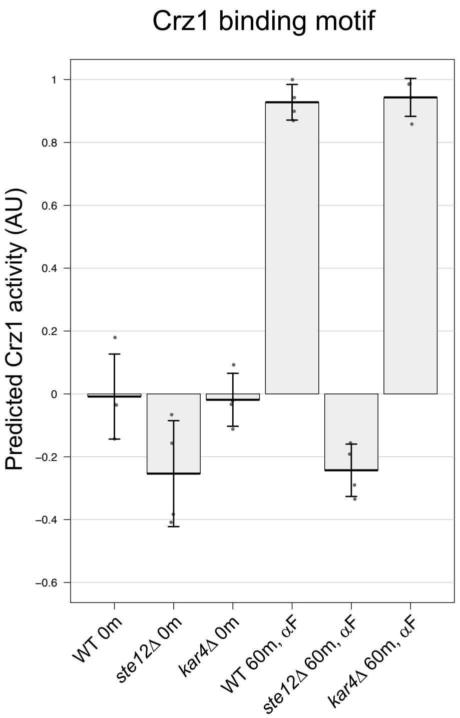
Crz1 binding motif
Predicted Crz1 activity (AU)
1
0.8
0.6
0.4
0.2
0
−0.2
−0.4
−0.6
WT 0m
ste12Δ 0m
kar4Δ 0m
WT 60m, αF
ste12Δ 60m, αF
kar4Δ 60m, αF

**Figure A1**

**Appendix Figure A1.** Crz1 predicted activity in the indicated samples, as **Figure 7b**. The full $\log_2$ fold-change gene expression matrix for all samples used in **Figure 2** was used as input. Units are derived from fitted score values (coefficients) and rescaled such that the largest absolute value is 1. Error bars indicate standard deviation. n = 4 samples per group.

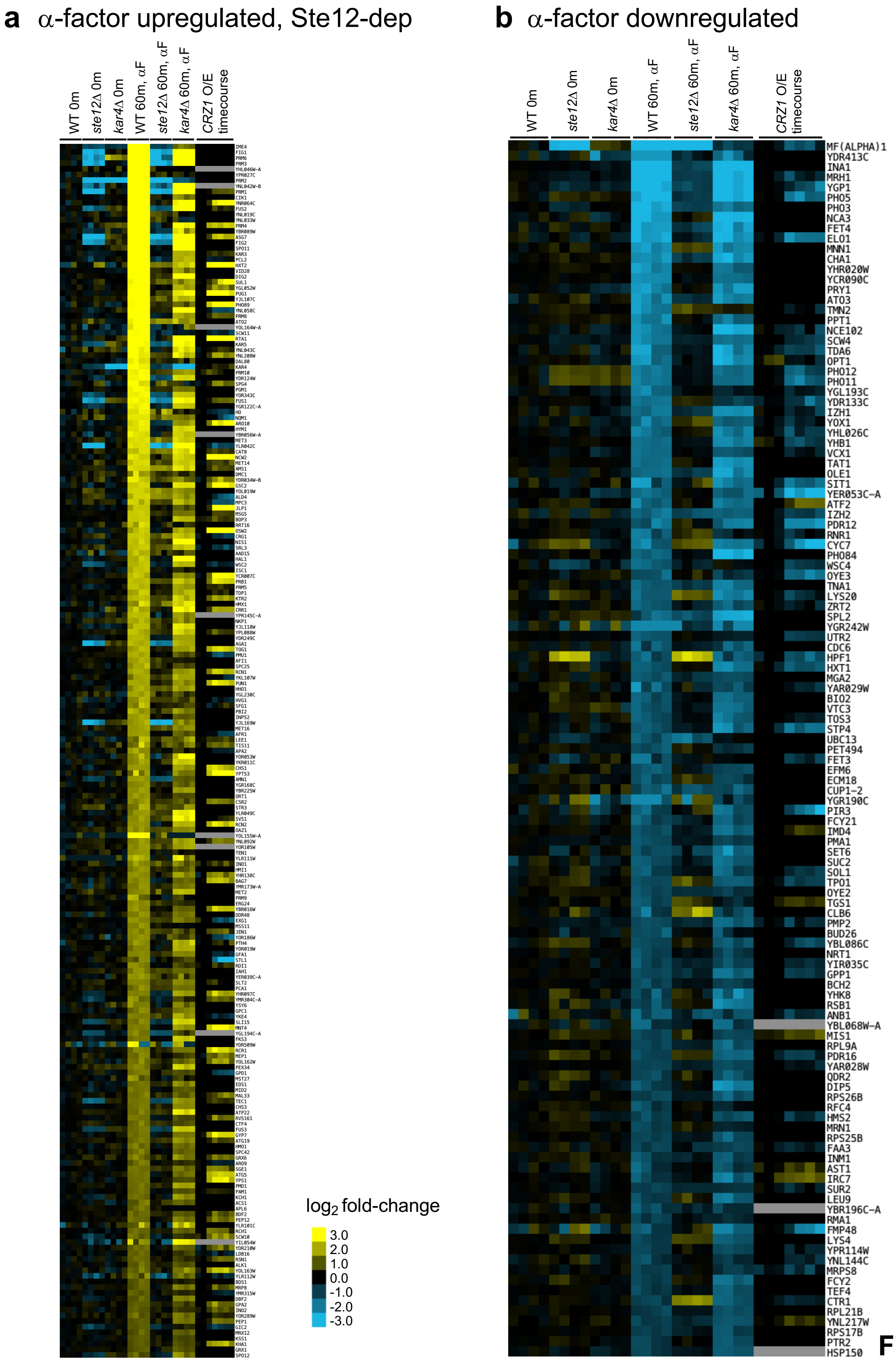
a α-factor upregulated, Ste12-dep
WT 0m
ste12Δ 0m
kar4Δ 0m
WT 60m, αF
ste12Δ 60m, αF
kar4Δ 60m, αF
CRZ1 O/E timecourse
log2 fold-change
3.0
2.0
1.0
0.0
-1.0
-2.0
-3.0
b α-factor downregulated
WT 0m
ste12Δ 0m
kar4Δ 0m
WT 60m, αF
ste12Δ 60m, αF
kar4Δ 60m, αF
CRZ1 O/E timecourse
MF(ALPHA)1
YDR413C
INA1
MRH1
YGP1
PHO5
PHO3
NCA3
FET4
ELO1
MNN1
CHA1
YHR020W
YCR090C
PRY1
ATO3
TMN2
PPT1
NCE102
SCW4
TDA6
OPT1
PHO12
PHO11
YGL193C
YDR133C
IZH1
YOX1
YHL026C
YHB1
VCX1
TAT1
OLE1
SIT1
YER053C-A
ATF2
IZH2
PDR12
RNR1
CYC7
PHO84
WSC4
OYE3
TNA1
LYS20
ZRT2
SPL2
YGR242W
UTR2
CDC6
HPF1
HXT1
MGA2
YAR029W
BIO2
VTC3
TOS3
STP4
UBC13
PET494
FET3
EFM6
ECM18
CUP1-2
YGR190C
PIR3
FCY21
IMD4
PMA1
SET6
SUC2
SOL1
TPO1
OYE2
TGS1
CLB6
PMP2
BUD26
YBL086C
NRT1
YIR035C
GPP1
BCH2
YHK8
RSB1
ANB1
YBL068W-A
MIS1
RPL9A
PDR16
YAR028W
QDR2
DIP5
RPS26B
RFC4
HMS2
MRN1
RPS25B
FAA3
INM1
AST1
IRC7
SUR2
LEU9
YBR196C-A
RMA1
FMP48
LYS4
YPR114W
YNL144C
MRPS8
FCY2
TEF4
CTR1
RPL21B
YNL217W
RPS17B
PTR2
HSP150

**Figure A2**

**Appendix Figure A2.** Comparison of pheromone-responsive upregulated (a) or downregulated (b) genes to *CRZ1* overexpression timecourse (HACKETT *et al.* 2020) as in Supplementary **Figure S4c**. Timecourse timepoints are (from left to right) 5, 10, 15, 20, 30, 45, and 90 minutes.

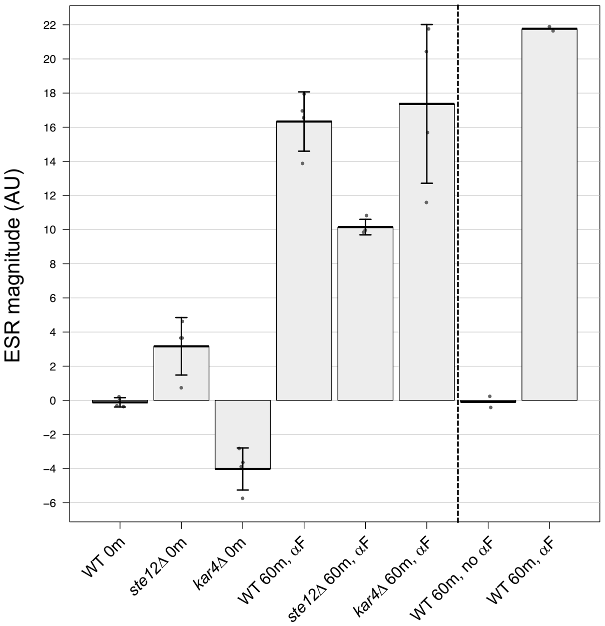
ESR magnitude (AU)
22
20
18
16
14
12
10
8
6
4
2
0
−2
−4
−6
WT 0m
ste12Δ 0m
kar4Δ 0m
WT 60m, αF
ste12Δ 60m, αF
kar4Δ 60m, αF
WT 60m, no αF
WT 60m, αF

**Figure A3**

**Appendix Figure A3.** Predicted ESR strength (see Methods) in indicated samples. Samples are normalized against their own wild-type controls (i.e., the left 6 conditions are normalized against “WT 0m” and the right 2 conditions are normalized against “WT 60m, no $\alpha$F). Samples from the left-group are additionally batch-corrected (i.e., samples from batch 1 are normalized against wild-type samples from batch 1, and samples from batch 2 are normalized against wild-type samples from batch 2). Error bars indicate standard deviation (n = 4). Bars indicate mean value and circles indicate raw data points. All t60 samples are significantly different from t0 samples (p-val < 0.01). P-values within the t0 group: wild-type vs. *ste12*Δ, p = 0.033; wild-type vs. *kar4*Δ, p = 0.008. P-values within the t60 group: wild-type vs. *ste12*Δ, p = 0.007; wild-type vs. *kar4*Δ, p = 0.699. P-values were calculated using a t-test with Bonferroni-Hochberg correction for multiple hypothesis testing.

| Motif | ID | P-value | E-value | Q-value | TP | FP | Enrichment Ratio |
|---|---|---|---|---|---|---|---|
| 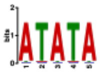 | Mot2 | 1.18e-3 | 8.65e-1 | 8.43e-1 | 51/108 (47.2)% | 1720/5840 (29.5%) | 1.62 |
| 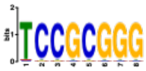 | Pdr1 | 5.36e-3 | 3.92e0 | 8.90e-1 | 30/108 (27.8%) | 959/5840 (16.4%) | 1.73 |
| 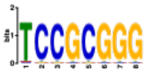 | Pdr3 | 5.36e-3 | 3.92e0 | 8.90e-1 | 30/108 (27.8%) | 959/5840 (16.4%) | 1.73 |
| 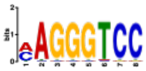 | Nrg2 | 7.31e-3 | 5.35e0 | 8.90e-1 | 18/108 (16.7% | 498/5840 (8.5%) | 2.04 |
| 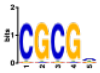 | Rsc3 | 9.12e-3 | 6.67e0 | 8.90e-1 | 41/108 (38.0%) | 1475/5840 (25.3%) | 1.52 |
| 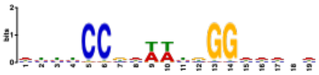 | Mcm1 | 9.39e-3 | 6.87e0 | 8.90e-1 | 42/108 (38.9%) | 1522/5840 (26.1%) | 1.51 |
| 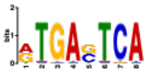 | Gcn4 | 1.09e-2 | 7.94e0 | 8.90e-1 | 26/108 (24.1%) | 845/5840 (14.5%) | 1.71 |
| 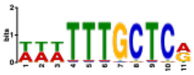 | Mac1 | 1.27e-2 | 9.30e0 | 8.90e-1 | 25/108 (23.1%) | 815/5840 (14.0%) | 1.71 |

**Figure A4**

**Appendix Figure A4.** Enrichment of known transcription factor motifs (see Methods) for genes downregulated in response to pheromone. “Logo” columns indicates motif logo, “Database” and “ID” columns indicate unique ID of the logo. “Alt ID” indicates the transcription factor associated with the motif. “TP” and “FP” indicate “true positive” and “false positive” rates, respectively. See SEA (Simple Enrichment Analysis) details for explanations on p-value, E-value (expected value), and Q-value (FDR-corrected p-value) calculations. Downregulated genes were used to determine true positives, and all other yeast genes (from the “simple” gene list, see Methods) were used as background to determine false positives. Note that in both sets 10% of genes were held-out for validation purposes to improve p-value accuracy (thus, the true positive denominator is 108 genes, not 119).

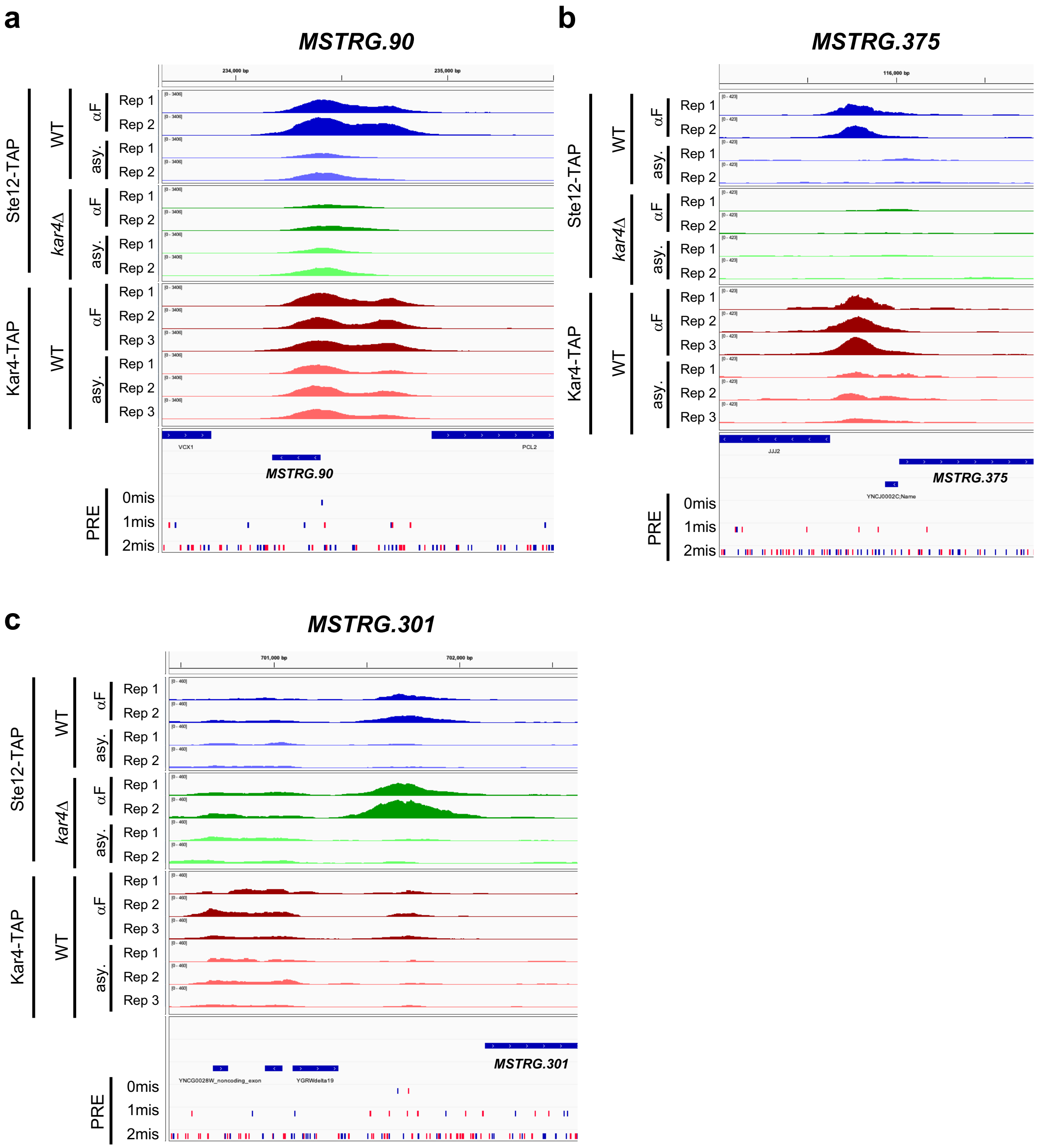
a
MSTRG.90
234,000 bp
235,000 bp
Ste12-TAP
WT
αF
asy.
kar4Δ
Kar4-TAP
Rep 1
Rep 2
Rep 3
VCX1
PCL2
MSTRG.90
PRE
0mis
1mis
2mis
b
MSTRG.375
116,000 bp
JJJ2
MSTRG.375
YNCJ0002C;Name
c
MSTRG.301
701,000 bp
702,000 bp
YNCG0028W_noncoding_exon
YGRWdelta19
MSTRG.301

**Figure A5**

**Appendix Figure A5.** Example loci as shown in **Supplementary Figure S7**, highlighting peaks associated with ITs during pheromone treatment. ITs are shown below genes but above PREs and begin with “MSTRG” (MSTRG is an automatically-assigned name prefix, meaning “Merged STRingtie Gene id”). a and b) Ste12 peaks that are close to protein coding gene promoters (*PCL2* in panel a and *JJJ2* in panel b) that are upregulated by pheromone. MSTRG.90 (a) is not significantly upregulated during the pheromone response in wild-type or *kar4*Δ cells. MSTRG.375 (b) is transcriptionally upregulated ~2.1-fold in wild-type cells and ~1.4-fold in *kar4*Δ cells (and not upregulated in *ste12*Δ cells, although it did not meet the qualifications for being Ste12-dependent by 2-fold, see Methods). c) A Ste12 binding site that is likely only associated with an intergenic transcript. MSTRG.301 (c) transcriptional upregulation is ~2.3-fold in both wild-type and *kar4*Δ cells (and not upregulated in *ste12*Δ cells).

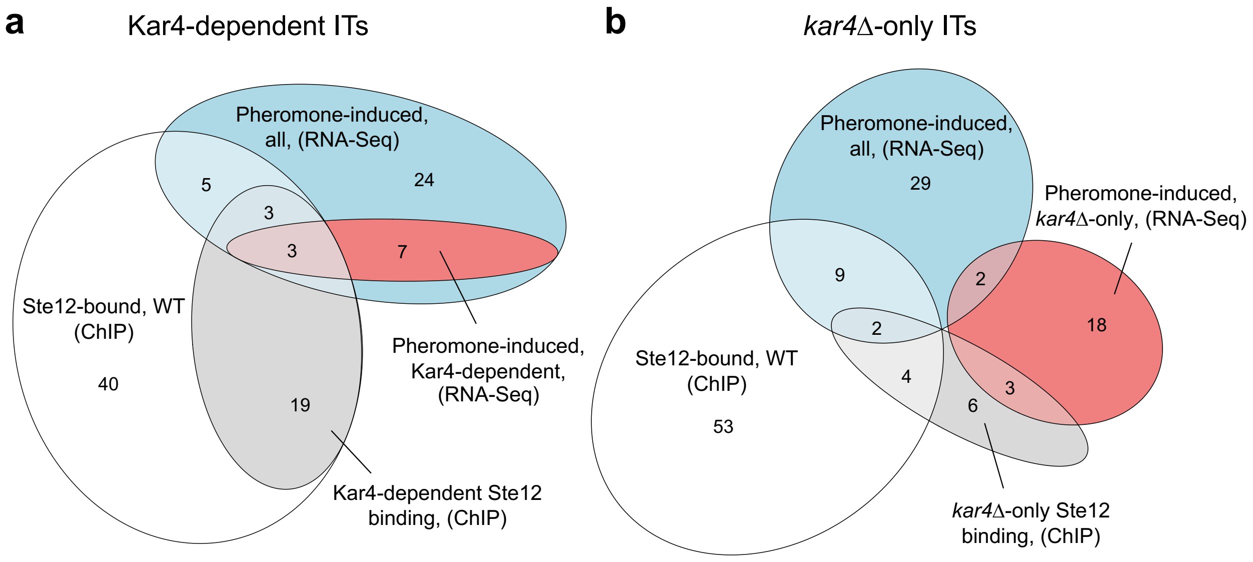


**c**

| Motif | Similar motifs | Sites |
|---|---|---|
| 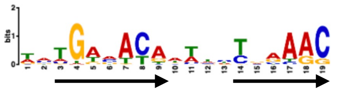 | Ste12 (H-T 4) | 34 (100%) |
| 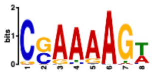 | Cup2 | 18 (52.9%) |
| 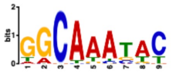 | Flo8/ Mss11 | 14 (41.2%) |
| 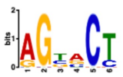 | Unknown | 16 (47.1%) |
| 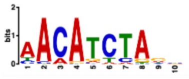 | Unknown | 16 (47.1%) |

**d**

| Motif | Similar motifs | Sites |
|---|---|---|
| 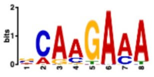 | Swi4/ Azf1 | 15 (78.9%) |
| 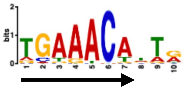 | Ste12 | 15 (78.9%) |
| 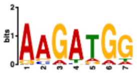 | Unknown | 14 (73.7%) |
| 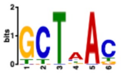 | Unknown | 8 (42.1%) |
| 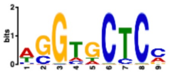 | Unknown | 8 (42.1%) |

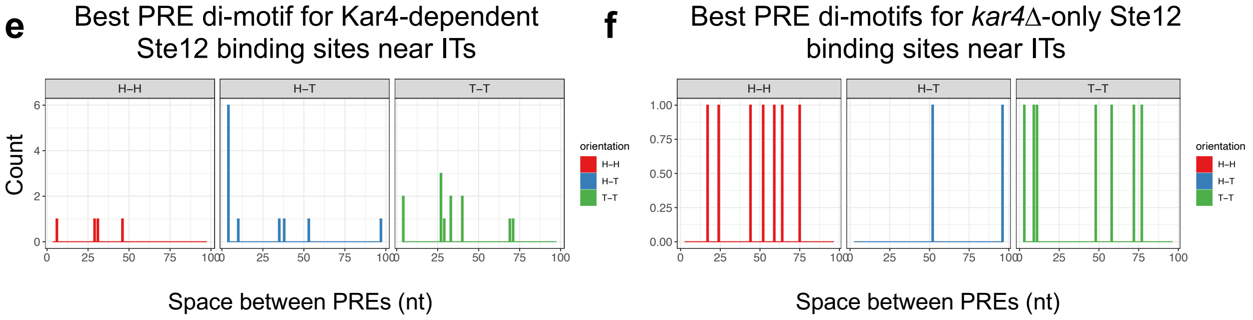


**Figure A6**

**Appendix Figure A6.** Characterization of Ste12-bound ITs during pheromone treatment. a) Venn diagram showing overlap between transcripts associated with Ste12-binding during pheromone treatment (by ChIP), either in total (white) or Kar4-dependent (grey), and transcriptionally pheromone-induced transcripts (by RNA-seq), either in total (blue) or Kar4-dependent (red). b) As in a, but rather than comparing to Kar4-dependent transcripts (wild-type vs. *kar4*Δ), comparing *kar4*Δ to wild-type (*kar4*Δ-only). Note that two small intersections could not be drawn: 1 transcript in common between the white (Ste12-bound, wild-type ChIP) and red (*kar4*Δ-only transcription) circles, and 1 transcript in common between the white (Ste12-bound, wild-type ChIP), red (*kar4*Δ-only transcription), and grey (Ste12-bound, *kar4*Δ-only) circles. c) de novo motif analysis as in **Figure 6b** for Kar4-dependent sites associated with ITs. d) As in c but for *kar4*Δ-only Ste12-binding sites. e) Best-match PRE di-motif histogram (as in **Figure 6c**) for Kar4-dependent sites associated with ITs (n = 27). f) As in e but for *kar4*Δ-only Ste12-binding sites (n = 16).

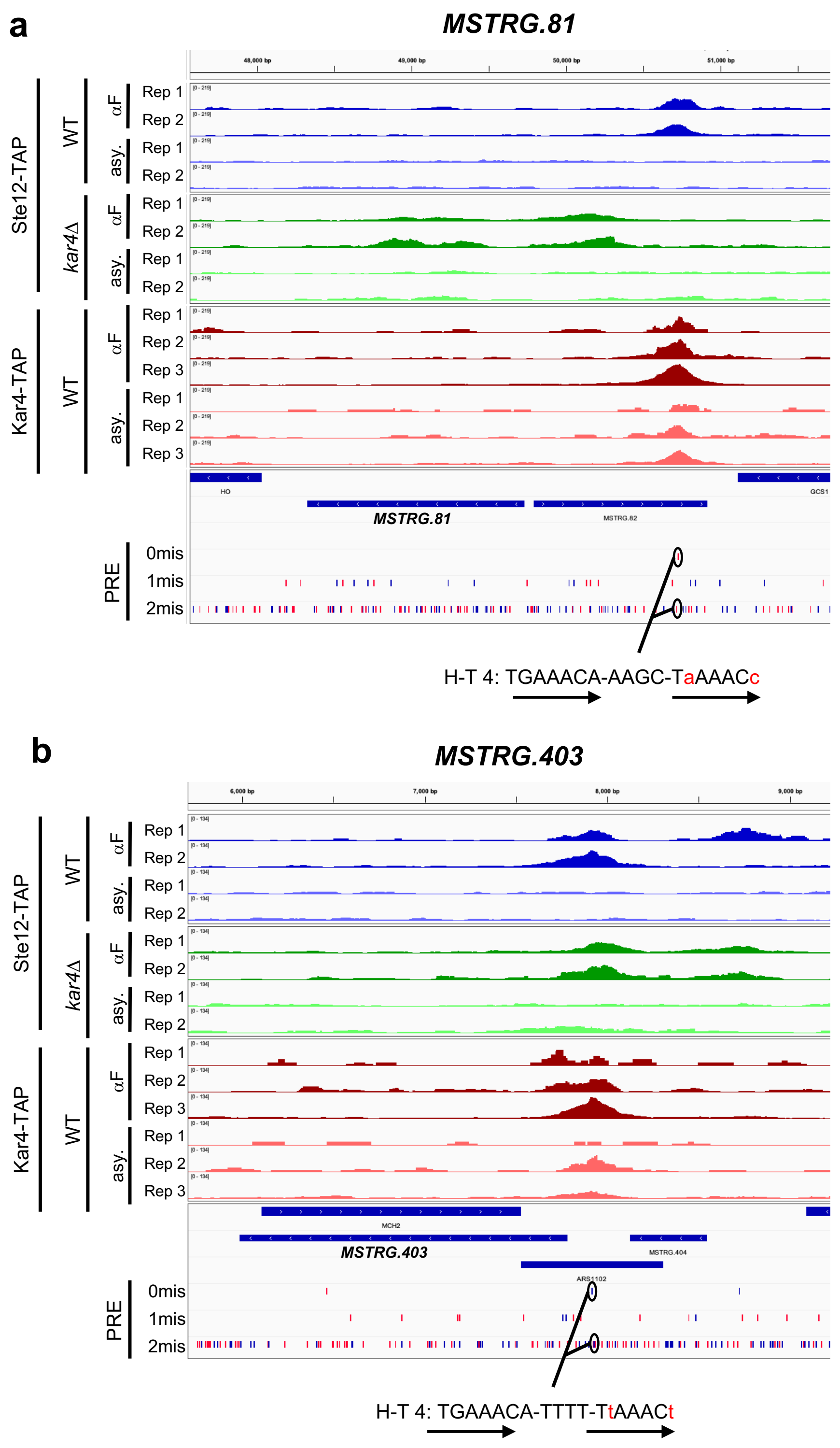
a
MSTRG.81
Ste12-TAP
WT
kar4Δ
αF
asy.
Rep 1
Rep 2
Rep 3
Kar4-TAP
HO
GCS1
MSTRG.81
MSTRG.82
PRE
0mis
1mis
2mis
H-T 4: TGAAACA-AAGC-TaAAACc
b
MSTRG.403
MCH2
MSTRG.403
MSTRG.404
ARS1102
H-T 4: TGAAACA-TTTT-TtAAACt

**Figure A7**

**Appendix Figure A7.** Top Ste12-bound and transcriptionally upregulated ITs as shown in **Appendix Figure A5**. H-T 4 motifs at the Ste12 binding site are given below the tracks. Note that for MSTRG.81 (a) the DNA sequence is reverse complemented from the reference sequence for clarity (i.e., the H-T 4 motif points toward MSTRG.81). Arrows indicate PRE locations and orientations.

**Appendix Supplemental File list**

File A1: Differential expression analyses for ITs

File A2: ChIP-exo peak annotations and differential contrasts including ITs

File A3: Best PRE di-motif annotations for ITs
